# Massively parallel interrogation of protein fragment secretability using SECRiFY reveals features influencing secretory system transit

**DOI:** 10.1101/241349

**Authors:** M. Boone, P. Ramasamy, J. Zuallaert, R. Bouwmeester, B. Van Moer, D. Maddelein, D. Turan, N. Hulstaert, H. Eeckhaut, E. Vandermarliere, L. Martens, S. Degroeve, W. De Neve, W. Vranken, N. Callewaert

## Abstract

While transcriptome- and proteome-wide technologies to assess processes in protein biogenesis are now widely available, we still lack global approaches to assay post-ribosomal biogenesis events, in particular those occurring in the eukaryotic secretory system. We here developed a method, SECRiFY, to simultaneously assess the secretability of >10^5^ protein fragments by two yeast species, *S. cerevisiae* and *P. pastoris,* using custom fragment libraries, surface display and a sequencing-based readout. Screening human proteome fragments with a median size of 50 - 100 amino acids, we generated datasets that enable datamining into protein features underlying secretability, revealing a striking role for intrinsic disorder and chain flexibility. SECRiFY is the first methodology that generates sufficient amounts of annotated data for advanced machine learning methods to deduce secretability predictors. The finding that secretability is indeed a learnable feature of protein sequences is of significant impact in the broad area of recombinant protein expression and *de novo* protein design.

## Introduction

The eukaryotic secretory system processes roughly a quarter of the proteome^1–3^, ensuring correct folding, assembly, and delivery of proteins to the extracellular environment, the plasma membrane, or membrane-bound organelles^4–6^. Model secretory cargos such as yeast carboxypeptidase Y (CPY), ɑ-1 antitrypsin (AAT), transthyretin (TTR), the Cystic Fibrosis transmembrane conductance regulator (CFTR), and the vesicular stomatitis virus G protein (VSVG) have been instrumental in understanding the function and regulation of many ER- or Golgi-resident proteins (for instance ^7–11^); yet, the precise features that enable or prevent secretory system transit of the thousands of other secretory proteins remain obscure. For example, it is generally unknown which chaperones are critical for assisting the folding of the different types of secretory protein domains, what sequence or structural features control ER export kinetics, or what determines glycan modification by Golgi glycosyltransferases. Studies that examine a broad range of proteins passing through the secretory system are integral to understanding how multiple processes integrate to produce the full set of secretory proteins. Unfortunately, most current approaches are unsuited to study a comprehensive range of protein folds after entry in the ER. Mass spectrometry (MS)-based proteomics is the predominant approach for the interrogation of post-translational events, but despite many technological advances, its breadth and depth is limited and decreases steeply with sample complexity; in routine MS setups, generally less than 70% of all transcribed mammalian protein-coding genes are detected^12,13^, and full protein coverage is rarely achieved.

In practice, the absence of such an integrated picture of the secretory system is most apparent in the unpredictability of heterologous protein secretion. Indeed, four decades after the recombinant DNA revolution, obtaining detectable levels of functional protein in a particular heterologous host system, secretory or not, is still principally a process of trial and error. Statistics from worldwide Structural Genomics Consortia initiatives suggest that the likelihood of progression from cloned target to soluble, purifiable protein lies between 20-50%, even with target selection^14,15^. These low and unpredictable expression success rates slow down progress in the many fields of basic and applied life science where recombinant protein production comes into play. Although expression screening of parallel constructs or variant libraries of a protein of interest to increase heterologous protein expression success rates has gained momentum^16–19^, it has to be repeated for each new target. Additionally, alternative comprehensive strategies to assess heterologous expression across entire proteomes have generally been limited to intracellular expression in *E. coli,* small proteomes, and cumbersome clone-by-clone strategies^20–23^. As the availability of large-scale protein expressability data will help to demystify which and why proteins fail to express, new methods for measuring expression in high throughput are needed.

We here developed an approach to evaluate the secretory potential (‘secretability’) of proteins on a proteome-wide scale. SECRiFY (<u>SEC</u>retability screening of <u>R</u>ecomb<u>i</u>nant <u>F</u>ragments in <u>Y</u>east) combines yeast surface display screening of protein libraries and a deep sequencing readout, enabling the systematic identification of heterologous polypeptides that can pass (or evade) the secretory quality control checkpoints of the yeast ER, Golgi, secretory vesicles and plasma membrane, and be secreted. As a first fundamental question to be addressed using this new methodology, we asked whether, given a particular sequence of a protein fragment, we could 1) learn what features contribute to its secretability and 2) generate machine-learned classifiers that predicts its secretability. Hence, we fragmented the human proteome and screened these fragments for secretability in two different yeast species, *Saccharomyces cerevisiae* and *Pichia pastoris (Komagataella phaffii).* We generated a large repository of more than 20,000 experimentally determined yeast-producible human protein fragments, available to the research community at https://iomics.ugent.be/secrify. These datasets of experimentally verified secretable fragments will assist experimental design of recombinant protein secretion. We used them to train (deep) machine learning models for secretability prediction, which unveiled sequence and structural determinants of productive secretory system transit, highlighting the utility of SECRiFY to provide further insight into the basic mechanisms of secretory processing. More application-focused implementation of SECRiFY (focusing on fixed-boundary protein domains or multi-domain fragments) should enable to generate databases of experimentally validated secretable native protein domain fragments, which could tremendously speed up experimental protein expression in many fields of study.

## Results

### Normalized fragment libraries for screening at domain-level resolution

Multi-domain proteins often fail to express or secrete in their entirety due to local issues with misfolding of particular protein areas, translation inhibitory sequences, protease susceptibility, the absence of stabilizing interaction partners or modifications, or toxicity. The structural, functional and evolutionary modularity of proteins in domains, however, implies that individual expression of certain protein parts, especially domains, can often nonetheless be achieved. Chopping up difficult proteins into experimentally tractable fragments has been exploited by structural biologists for years, both in rational target design as well as in random library screens for soluble expression^24–28^. Moreover, screening of protein domains or fragments can provide valuable information that is not immediately attainable or obvious from screens with full-length proteins^29^. Some domain-focused interactome studies, for example, have allowed immediate delineation of the minimal interacting regions and the detection of more interactors without increasing the number of false positives^30^. We thus rationalized that screening libraries of domains or domain-sized polypeptides, rather than full-length proteins, would allow for a higher resolution measurement of secretability across proteomes, and facilitate the identification of sequence or structural determinants of secretion.

Domain boundary prediction, however, is notoriously inaccurate, and even with a reliable estimate, small variations in the exact N- and C-terminus of the fragment can lead to dramatic differences in expressability^31^. Random approaches, on the other hand, can generate libraries of fragments that encompass most domains of a proteome by sheer oversampling. We therefore designed and built directional, randomly fragmented cDNA libraries covering the human transcriptome with fragments coding for approx. 50-100 amino acids, which is the median domain size of human proteins (**Fig. 1a, c**). Due to the large dynamic range of mRNA transcripts in human cells (in abundance of 4 orders of magnitude), however, capturing the full diversity of fragments would require unfeasibly large libraries, even at 100 bp resolution (**Fig. 1b**). We therefore reduced fragment abundance differences by relying on the second-order kinetics of nucleic acid rehybridization after denaturation, and subsequent digestion with the Kamchatka crab duplex-specific nuclease (DSN)^32,33^ (**Fig. 1c-d**). More abundant DNA species rehybridize faster and are therefore digested first; as such, even a single round of normalization substantially reduces abundance differences between DNA fragments (**Fig. 1g**). Crucially, this allowed us to downsize the libraries to a scale that is feasibly compatible with downstream cDNA library cloning and yeast transformation efficiencies (+/- 5*10^6^ - 5*10^7^).

**Fig. 1.**
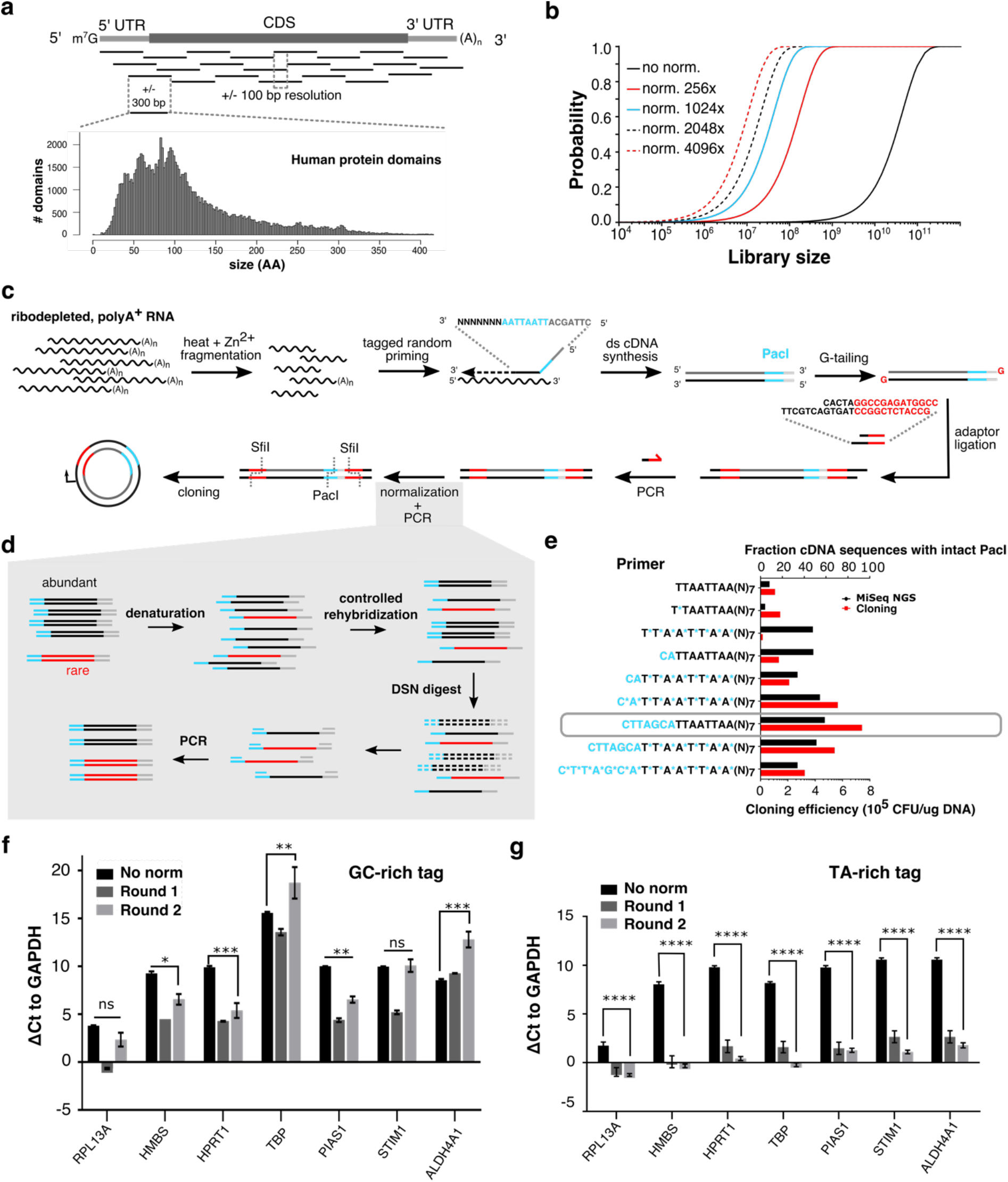
Capturing protein domains from transcriptomes with directional, normalized fragment libraries. **(a)** Most protein domains are between 50-150 amino acids (AA) long (lower left, Gene3D (v14.0.1) human protein domains, n= 104,734). Fragmentation of mRNA transcripts to 300 bp fragments should capture a substantial part of the domainome. At a +/100 bp resolution, on average 25 fragments would be sufficient to cover a typical transcript. (**b**). Estimated relationship between library size, at a hypothetical 100 bp resolution, and the probability of sampling any fragment, depending on the efficiency of fragment abundance normalization. **(c)** Fragment libraries are constructed by tagged random priming of fragmented polyA^+^ RNA, G-tailing, semi-single stranded adaptor ligation, PCR, and duplex-specific nuclease normalization before cloning into the yeast surface display vector. ds= doublestranded. (**d**) Abundant transcripts rehybridize faster than rare ones during kinetically controlled rehybridization after denaturation, and as such, digestion of double-stranded DNA with duplex-specific nuclease (DSN) can be used to normalize fragment abundance. (**e**) Effect of phosphorothioate bonds (blue stars) and buffer sequences (blue nucleotide sequence) on degradation of the PacI sequence in the tag, as measured by deep sequencing (black bars, upper axis) and restriction enzyme/ligase-based cloning into the surface display vector (red bars, bottom axis). The design with buffering bases alone (grey box) was the most effective. Primers are written from 5’ to 3’. CFU= colony-forming units, NGS= next-generation sequencing. (**f, g**) Abundance differences of various gene fragments compared to *GAPDH,* as ΔCt +/− SEM. All sequence abundances are nearly equalized when using a TA-rich (g), instead of a GC-rich (f) tag in the random primer, with normalization efficiencies up to a +/− 1000-fold (ΔCt of 10) for *HPRT1.* Two-way ANOVA with Tukey post-hoc, ns: non-significant, * p<0.05, ** p <0.01, *** p<0.001, **** p<0.0001. n= 9.

To ensure directionality, random primers were tagged with a rare-cutter restriction site (PacI), which is distinct from the SfiI site incorporated in the library adapters (**Fig. 1c**). We initially observed that the random primer tag sequence is susceptible to degradation due to endo- and exonuclease activity of the *E. coli* DNA polymerase I during second strand synthesis^34–37^. As a result, less than 20% of fragment sequences contained a full-length PacI site, negatively affecting ligation into the surface display vector (**Fig. 1e**). Both nuclease-resistant phosphorothioate bonds and buffer sequences could partially protect the tag from degradation, and for the final library design, we settled on the primer where protection efficiency was maximal (**Fig. 1e, grey bar**).

Tag composition also affected abundance normalization efficiencies. In an earlier design with a GC-rich tag, normalization was less effective (**Fig. 1f**) than the design with PacI tag (**Fig. 1g**), where an approx. 1000-fold normalization could routinely be obtained. The tag sequence is present on all sequence fragments, and most likely, when using a GC-rich tag, rehybridization kinetics (and therefore degradation) is dominated by the tag rather than by the sequence of the fragments themselves.

In all, this library construction protocol allows for efficient capture of protein-coding fragments tiled along eukaryotic transcriptomes. It is, to our knowledge, the first time that an effective method for normalization of tagged random-primed cDNA fragment libraries is reported, and it should find many applications in areas where the protein-coding potential of a cell needs to be effectively covered in expression libraries.

### SECRiFY as a platform for secretability screening in yeast

Relying on the sophisticated quality control (QC) machinery of the eukaryotic secretory system, which ensures efficient degradation of unstable or misfolded proteins before reaching the plasma membrane^19,20^, we further reasoned that surface display could be used as a proxy for productive secretion. As such, once cloned into the surface display vector and transferred to yeast, library polypeptides are directed to the secretory system by an N-terminal secretory leader sequence derived from the yeast a mating factor (MFα1 prepro), and furthermore on the yeast cell wall via C-terminal fusion to the GPI-anchoring region of the *S. cerevisiae* cell wall protein Sag1 (**Fig. 2a-b**). Fragments for which the fragment-Sag1 fusion successfully passes (or escapes) secretory system QC without proteolytic degradation are recognized through their N- and C-terminal epitope tags (FLAG and V5, resp.), and are segregated from the rest using iterations of high-efficiency magnetic- and fluorescence-activated cell sorting (MACS/FACS) (**Fig. 2b-c**). Finally, fragment identification and classification are achieved by deep sequencing of fragment amplicons from the unsorted and sorted cell populations. In short, SECRiFY assesses secretability, i.e. the potential of a polypeptide to transit through the secretory system of ER, Golgi, vesicles, and PM without degradation, in a manner that is independent of the original endogenous localization of the protein of interest. For the present study, we focus on basic principles of secretability. While in practice, any protein-coding mRNA pool is compatible with SECRiFY, considering its biomedical importance and structural complexity, we here focused on the human proteome for our screens, as encoded by the transcriptome of various human cell lines.

**Fig. 2.**
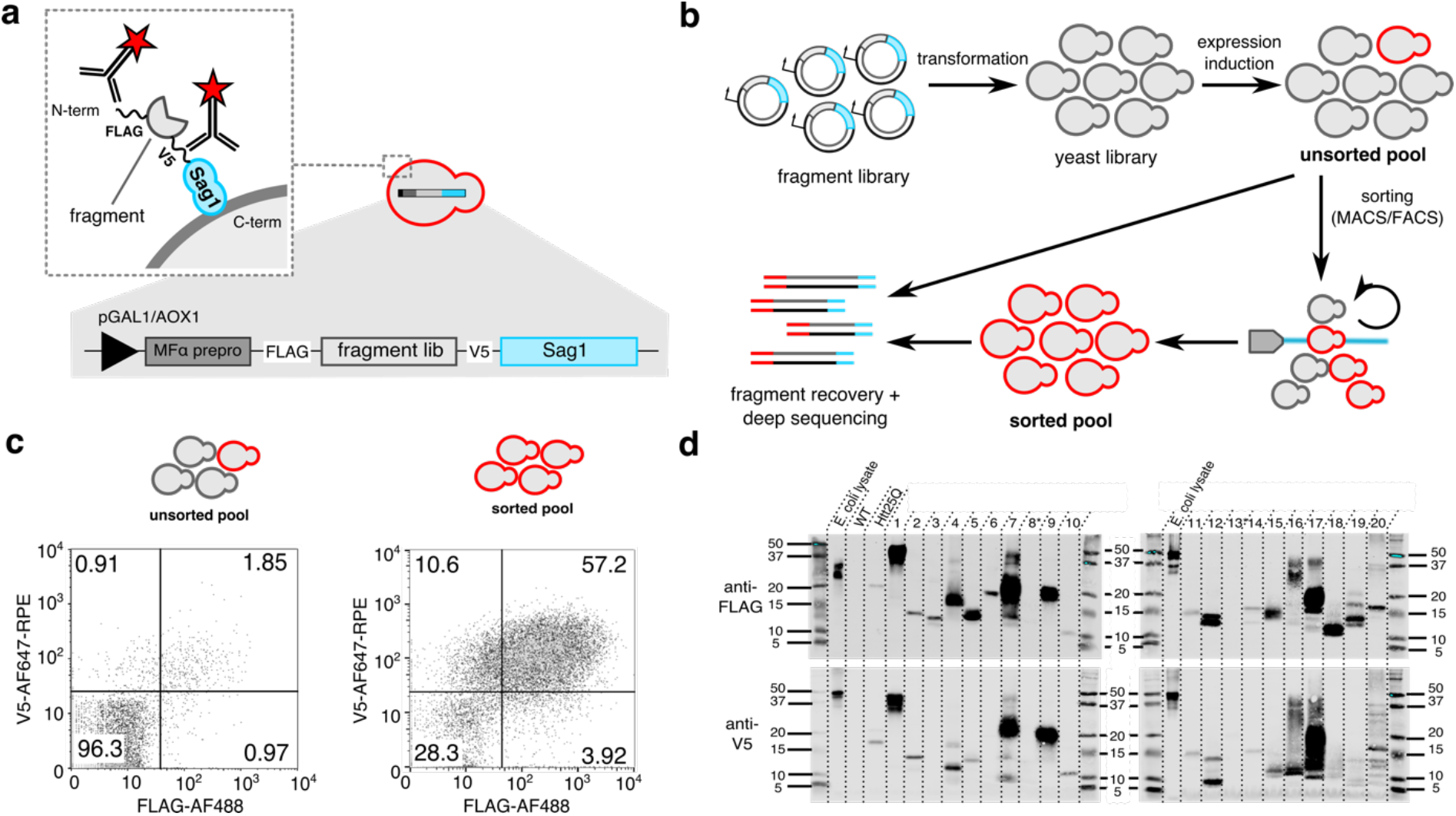
Screening for secretable protein fragments with the SECRiFY surface display platform. (**a**) Surface display as a proxy for secretion. Libraries are cloned downstream of an inducible promoter (pGal1 for *S. cerevisiae,* and pAOX1 for *P. pastoris),* a secretory leader sequence (MFa prepro), and a FLAG tag; and upstream of a V5 tag and the Sag1 anchor. Productive passage of library polypeptide fragments through the yeast secretory system leads to incorporation into the yeast cell wall, and displaying clones are identified through antibody-based labeling of the epitope tags. (**b**) SECRiFY screening workflow. Fragment libraries are transformed to yeast. After fragment expression induction, displaying FLAG^+^V5^+^ clones are sorted in multiple rounds of MACS/FACS. Fragments are identified by PCR recovery and deep sequencing of both sorted and unsorted cell pools. (**c**) Representative flow cytometry plots for SECRiFY screening of the human proteome in *S. cerevisiae.* After 3 rounds of enrichment (MACS/FACS/FACS), the fraction of double-positive (FLAG^+^V5^+^) clones increases roughly 30fold. (**d**) Western blot validation of fragment secretability after SECRiFY screening of the human proteome in *S. cerevisiae.* The majority of human protein fragments from sorted yeast cells can be expressed and secreted into the yeast medium in a Sag1-independent manner. Note that several fragments run as multiple species, likely due to heterogeneous processing and modifications such as O-glycosylation*. E. coli* lysate: antibody positive control, WT: *S. cerevisiae* R1158 medium (neg. control), Htt25Q: medium from *S. cerevisiae* secreting human Htt25Q (pos. control).

We first benchmarked the method by building a 1.96*10^6^ clone fragment library of the HEK293T transcriptome and performed triplicate screens in *S. cerevisiae.* On average, 1.76% ± 0.12% of library cells displayed a fragment with an intact N-terminus (FLAG-tag) and intact C-terminus (V5-tag) (**Supplementary Fig. 1**). Accounting for a 1/9 chance of up- and downstream in-frame cloning, this means that approximately 15.8% of in-frame fragments are detectably displayed and hence, potentially secretable. After a 32-fold enrichment of these double positive cells through a single round of MACS and two subsequent rounds of FACS (**Supplementary Fig. 1**), both pre- and post-sort population were sequenced at a per-base average coverage of minimally 150 reads. On average, 1.12*10^6^ unique fragments/replicate were detected, covering on average 26.45% ± 0.86% of the human canonical transcriptome with at least three reads (**Supp. Table 1-3**). To assess the secretion-predictive value of the method, we picked random clones from the sorted population of a single experiment (**Supplementary Fig. 2, Supplementary Table 4**) and tested the secretion of their encoded fragments when not fused to the anchor protein Sag1. The N- or C-terminal tags of 18/20 (90%) fragments could be reliably detected on western blot from the growth medium, and for 16/20 (80%) fragments, both tags were recognized (**Fig. 1d, Supp. Table 5**). As such, fragments displayed by sorted cells are indeed ‘secretable’ with a high probability. We further classified fragments into those that were enriched (also referred to as secretable) and those that were passively depleted (hence, not detected as secretable) by sorting, setting a cut-off on the enrichment factor 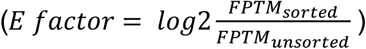 at 1 and −1, respectively, reflecting a minimal 2-fold increase and decrease in normalized sequence read counts after sorting. Of 170,226 inframe fragments commonly detected in the three experiments, 6.83% were consistently enriched in all three replicates, and 80.21% consistently depleted (**Supp. Table 6, Supplementary Fig. 3**). Thus, using this metric, these screens were reproducible with an 87.03% concordance between replicates. These final stratified groups of fragments, which were concordantly enriched or depleted, will further be referred to as secretable and depleted, respectively.

Since we only performed positive selection for secretable fragments during screening, the depleted fraction contains only passively depleted fragments, and the negative predictive value is relatively low (40-73%, **Supplementary Fig. 4**). In light of this, interpretation of features affecting secretion must focus on those that affect secretability, and not non-secretability. However, as there are ± 15 times as many fragments (data points) in this depleted set, this relatively low negative predictive value still provides for sufficient signal to allow machine learning methods to learn (see below).

Although we initially tested our method in the model yeast *S. cerevisiae,* in practice, the methylotroph *Pichia pastoris (Komagataella phaffii)* is an increasingly popular choice of host for recombinant protein production. Mostly, this has been attributed to this yeast’s formidable capacity for high-density growth, the secretion of relatively few endogenous secreted proteins, and the availability of very tightly repressed and extraordinarily strong inducible promoters derived from the yeast methanol metabolism genes^38,39^. Key to the adaptation of SECRiFY for use in *P. pastoris* was the development of a modified protocol for high-efficiency large-scale *P. pastoris* transformation, which resulted in an improvement in transformation efficiency of 2 – 3 orders of magnitude (**Online Methods** and **Supplementary Fig. 5**). While we previously observed a slight bias towards detecting small fragments in both enriched and depleted classes in our *S. cerevisiae* pilot screen, reducing the number of PCR amplification cycles during library generation for sequencing largely eliminated this trend, although small skews occurring during both cloning and sequencing were still observed (**Supplementary Fig. 6**). For the *P. pastoris* screens presented here, we first generated a new fragment library with slightly larger fragment insert sizes from the pooled transcriptome of four different human cell lines (SK-N-SH_RA, GM12878, HepG2 and MCF-7) originating from diverse human tissues (brain, blood, liver and breast) and selected to maximize the number of expressed human genes based on ENCODE transcriptome data^40^. Our high-efficiency transformation to *P. pastoris* generated a library with an estimated diversity of 9.8*10^6^ clones. Averaged over three replicate screens, 4.06% ± 0.68% of cells from this library were FLAG^+^V5^+^ (**Supplementary Fig. 7**), which, accounting for the frequent presence of multi-copy inserts (**Supplementary Fig. 8**), suggests that 12.18% of in-frame fragments are displayed and hence, potentially secretable. Sequencing the fragments of unsorted cells and cells sorted after 1 round of MACS and 1 round of FACS, we detected ± 1.5 million unique fragments per replicate, either in the enriched protein-displaying library, in the non-enriched starting library, or in both, covering 38.38% ± 2.25% of the human canonical transcriptome with at least three reads (**Suppl. Table 7-9**). Of the 215,004 in-frame fragments detected in all three of the replicates, 4.84% were classified as consistently enriched, and 65.75% as consistently depleted, leading to a 71% concordance between replicates (**Suppl. Table 10, Supplementary Fig. 9**).

Overall, these data show that SECRiFY is a reproducible and a reliable method to estimate the secretability of protein fragments. This dataset now represents by far the largest resource on eukaryotic secretability of protein fragments. To enable researchers to easily access and analyze the data from the screens in this study, we built an interactive database (http://iomics.ugent.be/secrify/) allowing visualization of the protein fragments detected in these screens and their mapping on available PDB structures (**Supplementary Fig. 10,** Figshare links to data in **Online Methods**).

### Secretable fragments are more flexible and disordered

Just as cytosolic protein expression is influenced by a variety of DNA, mRNA, and protein sequence or structural features and their complex interplays^41–44^, secretion of polypeptides will depend on a combination of multiple parameters, some of which are related to the unique environment and QC machinery of the ER and beyond. Even already at the simple level of general averaged parameters over our secretable vs depleted protein fragment collections, several intriguing observations emerged from our data.

We first examined whether secretable fragments differed from depleted ones in their probability to form secondary structures. To maximize the accuracy of feature prediction, fragments were filtered for size and exact match to Uniprot proteins, and condensed to an unambiguous subset of consolidated sequences in order to reduce sequence redundancy (see **Online methods**). Secondary structure prediction of this consolidated subset shows that secretable fragments most prominently have a lower propensity to form a-helical structures (p= 2.95*10^-124^, Mann-Whitney-Wilcoxon test) (**Fig. 3a, Supplementary Fig. 11a)**. Indeed, when clustering overlapping sequences to representative fragments and mapping these to solved structures in PDB (roughly 50% of representative fragments, **Supplementary Fig. 12**), secretability similarly inversely correlates with a-helical content (**Fig. 3b, Supplementary Fig. 13a**) (p= 2.35*10^-8^, Mann-Whitney-Wilcoxon test). In contrast, differences in β-sheet content are only minimal (**Fig. 3a-b**).

**Fig. 3.**
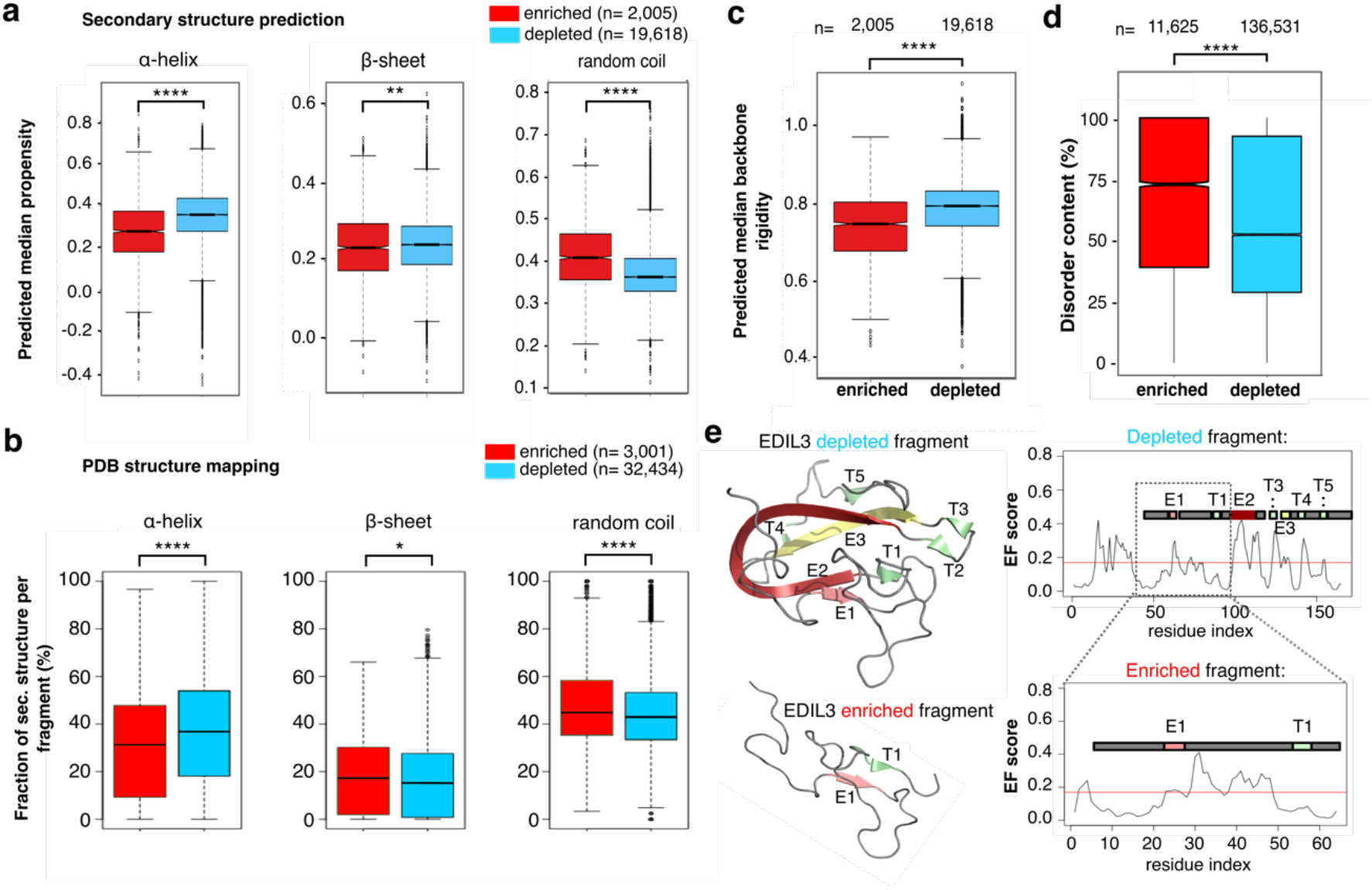
Patterns in secretable fragments. **(a)** Dynamine predictions of secondary structure propensity in subsets of consolidated enriched and depleted fragments in *S. cerevisiae.* Enriched fragments have a lower helical content (p= 2.95*10^-124^) and a higher random coil (p= 1.99*10^-127^) propensity, which is confirmed further by mapping representative fragments to known structures in PDB (a-helix p= 2.35*10^-8^, random coil p= 1.24*10^-4^) (**b**). Beta sheet differences are not as pronounced (p= 1.26*10^-3^ for Dynamine prediction and p= 0.02 for PBD mapping). (**c**) Enriched consolidated fragments are also predicted to be more dynamic than depleted ones (p= 5.08*10^-118^). (**d**) Similarly, the predicted disorder content in the total set of enriched fragments is significantly higher than in depleted fragments (p= <2.2*10^-16^). Twosided Mann-Whitney-Wilcoxon tests in (**a-d)**. * p < 0.05, ** p <0.01, **** p<0.0001. (**e**). Two overlapping fragments of the human protein EDIL3 differ in secretability outcome. Early folding (EF) propensity predictions suggest that for the depleted fragment regions E2, T3/E3 and R4 are likely the regions driving folding of the depleted fragment, and lack of these regions in the enriched fragment result in a change in secretability.

Since secretable fragments also tend to more readily form random coils than depleted fragments, based on secondary structure predictions (p= 1.99*10^-197^, Mann-Whitney-Wilcoxon test) as well as PDB mapping (p= 1.24*10^-4^, Mann-Whitney-Wilcoxon test) (**Fig. 3a-b**), we further examined how backbone dynamics and intrinsic disorder relate to secretability. As predicted using Dynamine^45,46^, secretable fragments are distinctly more flexible (p= 5.08*10^-118^, Mann-Whitney-Wilcoxon test) (**Fig. 3c**, **Supplementary Fig. 14a**). Disorder calculations on the full secretable vs depleted sets with RAPID^47^ also confirmed a higher average intrinsic disorder content in secretable fragments (p < 2.2*10-^16^, Mann-Whitney-Wilcoxon test) **(Fig. 3d**, **Supplementary Fig. 15**). In line with this, on average, fragments from both subsets appear compositionally biased. A larger fraction of secretable fragments has a higher proportion of negatively charged residues and prolines, and a tendency towards lower hydrophobicity (**Supplementary Fig. 16**). Possibly, this increase disorder in secretable fragments reflects how unstructured fragment sequences that lack typically exposed hydrophobic amino acids are missed by ER chaperones and can subsequently travel downstream. This is particularly striking since endogenous secretory system proteins in both human and yeast are, on average, less disordered than the whole proteome, both when considering overall disorder content (p < 2.2*10^-16^ and p= 2.46*10^-5^ resp., Fisher exact test) as well as absolute number of disordered amino acids (p < 2.2*10^-16^ and p= 9.44*10^-10^ resp., Fisher exact test) (**Suppl. Table 11-12**), suggesting evolutionary counterselection.

Increasing the fidelity of the above findings, all feature enrichment observations were reproduced in the *P. pastoris* SECRiFY screens (**Supplementary Tables 7-10, Supplementary Figs. 6-18**). In addition, our conclusions remained unchanged when choosing alternative criteria for defining secretable vs depleted fragment sets, illustrating the robustness of our observations.

### In silico secretability prediction with machine learning

For the first time, our SECRiFY method generates secretability data at a scale at which training of predictive machine learning classifiers becomes feasible. To study the presence of discriminatory features in the dataset, we explored two distinct approaches: one based on feature engineering together with gradient boosted decision tree modeling^48^, and a deep learning approach based on convolutional neural networks (CNNs)^49^.

The gradient boosting classifier requires a fixed input size. Therefore, a series of manually engineered input features were proposed, based on physicochemical properties, sequence length and amino acid frequencies. Ten individual classifiers were trained using different properties, and an ensemble of those was constructed using another gradient boosted classifier taking the outputs of the individual classifiers as input. The deep learning approach involved a CNN taking a one-hot encoding as input, followed by three blocks of convolutional, max pooling and dropout layers. We explored different strategies to deal with the variable input size, as this is not supported by standard CNN architectures. A global max pooling layer yielded the best overall results. This layer is finally connected to a dense layer, followed by an output layer with a softmax.

Fragments shorter than 50 amino acids were removed from both the *S. cerevisiae* and *P. pastoris* datasets, as those are likely not long enough to properly fold, which mitigates their relevance. Using a restrictive 10-fold cross-validation scheme, where we made sure that protein fragments originating from the same gene were included in the same fold, we compared the classifiers based on the area under the receiver operating characteristic curve (AUROC). Gradient boosting achieved an AUROC of 0.781 and 0.772 on the *S. cerevisiae* and *P. pastoris* datasets, respectively, whereas the CNNs achieved AUROCs of the same magnitude, 0.779 and 0.768 (**Fig. 4a**). Classification results of both classifiers thus confirmed the presence of distinctive features within both secretable and depleted subsets of the data. We observed a strong correlation between the predicted values for the two approaches, with Pearson correlation coefficients of 0.810 and 0.887 on the respective datasets (**Fig. 4b**), which suggests that the two models learned to use similar distinctive features in the data.

**Fig. 4.**
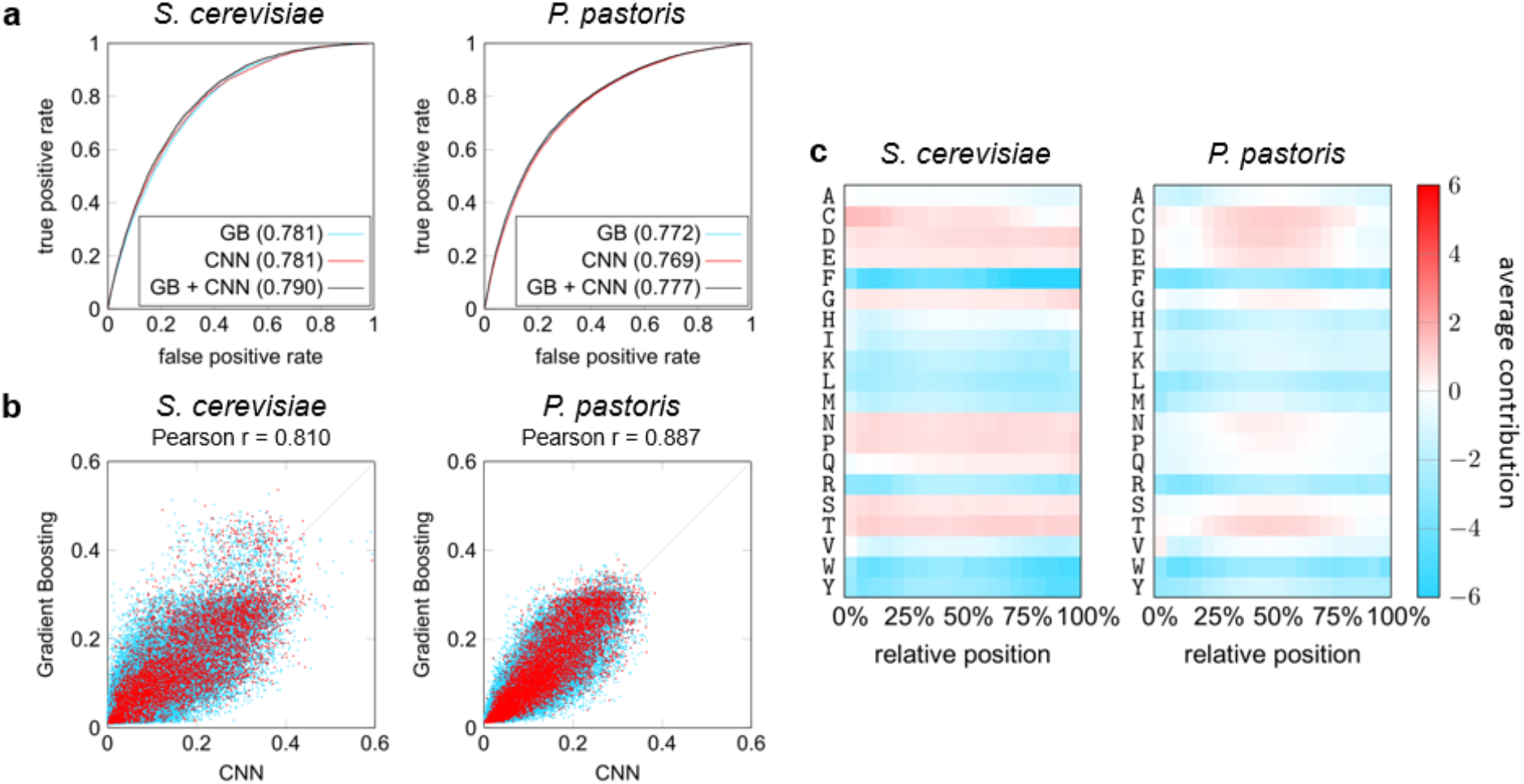
Machine learning for secretability prediction. (**a**) Evaluation (expressed in AUROC) of the gradient boosting (GB) and convolutional neural network (CNN, one model randomly selected out of the ten trained models) approaches, as well as an ensemble taking the average predictions of both. (**b**) Correlation between the predicted values of the GB and CNN models. Secretable data samples are shown in red, depleted samples in blue. (**c**) Average contribution of individual residues when occurring in different parts of the sequence. For each sequence in a test set, the contribution towards a *positive* prediction (secretable) is calculated for each individual residue. Contributions are then normalized, with absolute values of all contributions in a sequence adding up to 100 on average.

Feature importance analysis using attribution methods led to compelling insights in the decisions of the CNN (**Fig. 4c-d**). Aggregation of individual attribution maps on the amino acid residue level indicated that the influence of individual residues on secretability is largely independent of their position in the sequence. Strikingly, there is a positive bias towards smaller residues, in line with our biophysical predictions that random coils are more readily formed in secretable fragments. Negatively charged residues also seemed to substantially contribute to secretability, confirming the pattern we picked up looking at simple averaged parameters across the full dataset. Similarly, a negative bias was observed towards all hydrophobic amino acids, affirming our earlier observations.

### Secretable fragments are enriched in certain folds and domains

Other features that affect folding in the ER might influence secretability. Although increased presence of N-glycosylation sequons or uneven number of cysteines could increase ER retention of fragments, we did not observe clear differences in number of Cys or N-glycosylation sequons (**Supplementary Fig. 16f-g**). Furthermore, secreted and depleted fragments do not substantially differ in their predicted propensity to collapse into folded structure (EFoldMine^50^ prediction, **Supplementary Fig. 14b**). Nonetheless, in select cases where depleted and enriched fragments overlap in sequence on the same protein, the presence or absence of regions that are most likely to fold rapidly often correlated with secretability (**Fig. 3e**).

Beyond these parameters, we noticed clear differences in protein folds and domain architectures represented in both fragment groups. Of those fragments that mapped to known structures, secretable fragments are enriched in distorted sandwich and β complex folds compared to depleted fragments, suggesting that these folds are potentially more stable in the secretory environment, while the opposite is true for proteins with, for example, an a horseshoe architecture (**Supplementary Table 13**, Figshare links to data in **Online Methods**). Similarly, certain Pfam domains, such as the AAA18-domain (PF13238), are more prominent in enriched fragments than depleted fragments, while many typically cytoplasmic domains such as ribosomal protein domains or the tetratricopeptide repeat (TPR, PF13181) are found exclusively in depleted fragments (Figshare links to data in **Online Methods,** see also **Supplementary Fig. 18**). This illustrates that sequence- and fold-contextual patterns of features still contain much information that is not apparent from averaged parameters.

### Protein secretability does not correlate with endogenous secretion

Most patterns found to be enriched in secretable fragments are not recapitulated in endogenous secretory proteins. We therefore further evaluated whether the human proteins from which secretable fragments were derived, were enriched in secretory proteins. Since many proteins produced both secretable and depleted fragments in our screens, we considered only those proteins for which no depleted fragments were found. In both *S. cerevisiae* and *P. pastoris* screens, the proportion of secretory proteins (i.e., with a signal peptide in the endogenous setting) in this set was not significantly higher than the fraction of secretory proteins in the human proteome (Fisher’s one-sided exact test, p=0.9183 (*S. cerevisiae)* and p=0.421 (*P. pastoris),* **Table 1**). This disconnect between highly secretable fragments and endogenous secretory proteins likely reflects how features that determine highest efficiency passage through the secretory system are not the most important components of evolutionary selective pressure for secretory proteins.

**Table 1.** Secretory enrichment of secretable proteins.

|  | **with signal peptide** | **no signal peptide** | **p-value one-sided**<br>**Fisher exact test** |
| --- | --- | --- | --- |
| **Human proteome** | 3574 | 16548 | NA |
| **SECRiFY secretable**<br>**(*S. cerevisiae*)** | 17 | 109 | 0.9183 |
| **SECRiFY secretable**<br>**(*P. pastoris*)** | 24 | 104 | 0.421 |

## Discussion

Despite the tremendous strides made in the field of recombinant protein production, heterologous expression remains highly unpredictable. A deeper understanding of the intricate ways in which different processes integrate to produce the full set of secretory system proteins, heterologous or not, is only slowly emerging. Although the study of model secretory proteins has led to substantial progress in the field, a more global approach is needed to gain a more profound and comprehensive characterization of the factors that influence secretion.

Our SECRiFY method assesses the secretability of proteins on a proteome-wide scale and at domain-sized resolution by yeast. To this end, inspired by developments in the field of massive parallel sequencing library construction and random approaches to protein engineering, we first developed a streamlined method for the construction of a directionally cloned, normalized, and random primed cDNA fragment library. This novel combination of features enabled us to screen the human proteome for secretability at an unprecedented scale and depth compared to previous studies^52,53,19^. By extension, libraries of this type could be valuable alternatives in other applications where fragment screening is beneficial, such as high throughput proteinprotein interactomics.

In SECRiFY, yeast surface display screening of these libraries is combined with high efficiency cell sorting and deep sequencing to segregate and identify protein fragments that can productively pass through the yeast secretory system. As such, we demonstrated that the secretability of protein fragments across entire proteomes can be verified experimentally in an efficient, systematic, high-throughput and reproducible manner. Although we here used the human proteome as a testcase, our approach is generic and can be used to screen any eukaryotic, or with minor adaptations, even prokaryotic proteome. Already, the databases we have generated in this work constitute by far the largest resources of such yeast-secretable human protein segments, which will be useful in structural studies (where high levels of proteins are required for crystallization), immunological experiments (where recombinant production is needed for vaccine development or antigen discovery), and biochemical characterizations of particular proteins (which also necessitates a minimal protein amount). Remarkably, using both a gradient boosted method based on feature engineering, and an end-to-end trainable convolutional neural network approach, we achieved an AUROC of up to 0.790 for *S. cerevisiae* and 0.777 for *P. pastoris.* Practically, this means that secretable and depleted fragments have properties that allow for discrimination, even without prior knowledge of their nature. This unbiased approach confirmed our hypothesis-driven observations that biophysical features relating to secondary structure and flexibility affect secretability. Secretability is thus indeed a learnable feature of protein sequences, a finding that will benefit recombinant protein expression and *de novo* protein design.

From a fundamental biology perspective, it is likely that SECRiFY will provide a means to characterize the substrate scope of secretory system processes that regulate secretory protein passage through the eukaryotic secretory system in a proteome-wide manner. This complements existing methods, such as ribosome profiling^54^, which deal with protein biogenesis prior to passage through the secretory system.

Our screens, and the combined hypothesis-driven and unbiased data mining of the data, uncovered that secretable fragments tend to be less a-helical, more flexible and more intrinsically disordered than fragments without significant display. Chaperones responsible for secretory system quality control are generally poised to recognized mostly exposed hydrophobic stretches^55–57^, so conceivably, flexible yet polar and charged fragments would avoid these interactions and quickly progress towards cell wall incorporation. Limited proteolysis measurements also recently confirmed the inverse correlation of in-cell thermal stability with intrinsic disorder, a-helical secondary structure, and aspartate content^58^. Although these measurements involved proteins in their endogenous context in lysed cells, and lack of dataset overlap precluded direct comparison of secretability and thermal denaturation, it would be intriguing to investigate the relationship between secretability and stability. Even though our results show a clear correlation between disorder and secretability, for ordered proteins, quality control mechanisms in the secretory system will generally efficiently remove unfolded or unstable proteins. Indeed, recent limited proteolysis screens of known small protein domains present in PDB or Pfam suggest that structurally defined surface-displayed domains have an overall high stability score^59^.

Fragment detectability effects may also contribute to the observed enrichment of aspartates and glutamates. Phosphomannose proteins of the yeast cell wall, which confer it a net negative charge, may be repelled by negatively charged displayed fragments, leading to more efficient antibody staining due to increased accessibility. Our observation that more fragments with a high overall positive charge (Lys + Arg content) are not secreted could relate to translational ribosomal stalling at polybasic stretches^60^. Although proline-rich proteins are generally localized in the nucleus or cytoplasm^61^, and are associated with ribosome slowdown^62^, it is possible that the intrinsic ability of prolines to ‘lock’ conformations or reduce conformational freedom enhances secretory protein stability and hence, display and secretion. Polyproline or pro-rich stretches are also known motifs for binding to a wide variety of other proteins^29,63,64^; as such, the human oxidoreductase ERp57 binds calnexin and calreticulin at their pro-rich motif, and peptidyl-prolyl isomerases often act in complexes associated with multiple chaperones. Prolines are also used as gatekeeper residues against aggregation^65^, and perhaps by extension, also against degradation and therefore display.

Secretable fragments are not enriched in secretory proteins. SECRiFY detects secretion at the fragment level, potentially causing some features affecting full-length protein secretion to be missed. Nonetheless, the absence of correlation also underlines that SECRiFY assesses secretability, i.e. the capacity to be secreted, rather than actual endogenous cellular localization to the secretory system. Indeed, just like most proteins are only marginally stable, endogenous secretory proteins evolved for function, not secretability.

Arguably, our method also has its limitations. In the current SECRiFY setup, secretability was measured in the sequence context of the a mating factor prepro sequence at the N-terminus, and the Sag1 cell wall protein at the C-terminus. While results from our and other labs have indicated that for several single proteins, display efficiency correlates with relative secretion levels, it cannot be excluded that, at least for certain fragments, both leader sequence and the +/− 300 amino acid Sag1 anchor might differentially influence fragment folding, solubility, or stability. In *E. coli,* fusion to large proteins such as SUMO, the *T. harzanium* cellulose binding domain (CBD), or to maltose binding protein (MBP) is an often used strategy to promote ‘passenger solubilization’, although again, effects vary depending on the protein^66,67^. Considering the vectorial nature of translation, a C-terminal fusion, as is the case in our setup, is nevertheless generally deemed less perturbing than an N-terminal fusion, although this is not absolute. Sag1 is also a GPI-anchored protein, affecting the entry pathway into the ER^68–70^. Similarly, the prepro leader sequence, with its multi-step processing and preference for posttranslational translocation^71–73^, may bias secretability of certain fragments. It remains to be determined whether similar patterns will emerge with different secretory leaders, anchors, promoters, untranslated regions, or growth conditions.

Display also imposes limitations on the dynamic range of the method, as there is an upper limit to the number of molecules that the cell wall can accommodate. Generally, this is in the range of about 10^4^ molecules per cell^74,75^. Thus, perturbations affecting secretion efficiency in these higher ranges may be missed.

In all, with SECRiFY, we here show that our fragment sequence library is proof-of-concept for massively parallel assessment of passage through the secretory system, providing the opportunity the learn which features influence secretability, and what rules sequences must abide by for successful transit through the yeast secretory system. We anticipate that our method and its next-generation derivatives will be of great value in both protein engineering and fundamental studies of the secretory system.

## Acknowledgements

The authors thank the VIB Nucleomics Core for Illumina sequencing, as well as M. Vuylsteke for statistical insights, K. Vandewalle for help with western blot sample preparation, and Y. Dondelinger for help with plasmid construction. We thank Lennart Martens for freeing up time to discuss the project and to allow D.M, D.T., E.V. and P.R. to spend time on this project away from their main tasks. This work was supported by a Ghent University BOF PhD Fellowship (M.B., H.E.), a PhD Fellowship from the Research Foundation Flanders (FWO) (M.B., H.E.), an FWO research grant (G.0276.13N) (N.C.) and an ERC Consolidator grant no. 616966 (N.C.). E.V. is a postdoctoral research fellow of the Research Foundation Flanders. P.R. and W.V. acknowledge support from the Research Foundation Flanders through grant number G.0328.16N. W.D. and J.Z. are funded by the Ghent University Global Campus.

## Author Contributions

M.B. performed library constructions, methods development and optimization, screenings, sequencing data processing and analyses, and wrote the manuscript. P.R. performed structure-based data-interpretation under the supervision of E.V. and W.V. W.V. ran, analyzed and visualized the biophysical predictions. S.D. and R.B. developed the gradient boosting classifiers, W.D. and J.Z. developed the deep learning classifiers, and together they analysed the prediction results and feature contributions. D.M. and D.T. developed the website interface and the underlying database. B.V.M. and H.E. provided additional experimental support. N.C. conceived the project, and assisted in experimental design, interpretation, and manuscript writing. All authors read, revised and approved the final version of the manuscript.

