## Supplementary material for "Massively parallel interrogation of protein fragment secretability using SECRiFY reveals features influencing secretory system transit": Materials and Methods

### Online methods

##### Plasmid construction

All restriction digests, PCRs, plasmid preparations and DNA purifications were performed according to the reagent/kit manufacturer's guidelines unless stated otherwise. Transformations to chemically competent *E. coli* MC1061 cells were done by heat shock, and cells were plated on LB agar (5 g/l bacto yeast extract, 10 g/l bacto tryptone, 10 g/l NaCl, 15 g/l agar) with the appropriate antibiotics unless noted otherwise. When working with plasmids containing the zeocin resistance cassette, *E. coli* TOP10 cells were used and plated on low salt LB (5 g/l bacto yeast extract, 10 g/l bacto tryptone, 5 g/l NaCl, 15 g/l agar) agar plates containing 50 μg/ml zeocin. After initial restriction digest/colony PCR/insert sequencing checkups of constructed plasmids, final plasmids were fully sequenced by the VIB Genetic Sequencing Facility using Sanger sequencing before use.

The *S. cerevisiae* surface display plasmid (pSSDSfiIPacI-FLAGV5-Gal1) was generated by recombination-based assembly of 3 fragments: linearized p415-Gal1-noLac as vector backbone (*GAL1* promoter, *CYC1* TT, CEN/ARS, *LEU2* marker), a PCR product of pBluescript-ScCatch (FLAG-ministuffer-V5-Sag1), and a PCR product of the MFα1 prepro signal from pGal1-MF. PCR products were fused by overlap extension PCR, and the resulting product was recombined with linearized vector in a 30 min RT CloneEZ reaction (GenScript) and transformed to *E. coli*. To facilitate subsequent cloning in pSSDSfiIPacI-FLAGV5-Gal1, the small stuffer between FLAG and V5 was further replaced by a large stuffer fragment via Gibson Assembly, generating pSSDSfiIPacI-FLAGV5-Gal1-stuffer. For this, pSSDSfiIPacI-FLAGV5-Gal1 was digested with an equimolar amount of a SfiI-site containing oligo (A136) using restriction enzyme SfiI (NEB) for 50°C. The reaction was cooled and PacI (NEB) was added, and digestion was continued at 37°C for 1h. Purified vector fragment was combined with the PCR amplified stuffer fragment for Gibson Assembly. Similarly, an insertless display vector, in which the Sag1 is preceded by in-frame FLAG and V5 tags, was also constructed to function as 'empty display' control for subsequent flow cytometry experiments (pSSDSfiIPacI-FLAGV5-Gal1-EV). pSSDSfiIPacI-FLAGV5-Gal1 was thus digested by BamHI/XhoI (Promega), and purified backbone was combined with amplified FLAG-V5 in a Gibson Assembly reaction. For Sag1-less secretable expression, we also constructed a vector similar to pSSD but lacking the Sag1 coding sequence. This plasmid, pSCASfiIPacI-FLAGV5-Gal1, was constructed by PCR, phosphorylation and blunt relegation. A long stuffer-containing version of this plasmid, pSCASfiIPacI-FLAGV5-Gal1-stuffer, was constructed using the same procedure as for pSSDSfiIPacI-FLAGV5-Gal1-stuffer construction.

The *P. pastoris* surface display vector pPSDZeoSfiIPacI-FLAGV5-AOX1 was made by switching up the pPSDSfiIPacI-FLAGV5-AOX1 backbone for the pPICZ backbone through HindIII and NotI digest (both Promega), purification from gel, dephosphorylation of the pPICZ backbone, and ligation. Vector pPSDZeoSfiIPacI-FLAGV5-AOX1-stuffer was furthermore constructed by inserting part of the sequence for α-galactosidase from pPICZαGalMycHis between FLAG and V5 using SfiI/PacI restriction digest and ligation. An insertless display vector, in which the Sag1 is preceded by in-frame FLAG and V5 tags, was also constructed to function as 'empty display' control for subsequent flow cytometry experiments (pPSDZeoSfiIPacI-FLAGV5-AOX1-EV).

##### Yeast strains

*S. cerevisiae* strain R1158 (*MATa URA3*::*CMV-tTA his3Δ1 leu2Δ0, met15Δ0*) was obtained from Open Biosystems, frozen as slant in 15% glycerol at -80°C, and grown on SD-Ura (0.67% yeast nitrogen base w/o amino acids, with ammonium sulfate; 2% dextrose; 0.077% CSM-Ura dropout mix; 17g agar; pH 5.8) plates unless noted otherwise.

All *Pichia pastoris* work was performed in strain GS115 (*his4*)[1], grown in YPD media (10 g/L yeast extract, 20 g/L dextrose, 20 g/L peptone) supplemented with various concentrations of zeocin and set to various pHs as indicated, and supplemented with 17 g/L agar for plates. All plates were always freshly cast or kept in the dark at 4°C for maximum one week.

##### Human cell lines

HEK293T cells were cultured at 37°C in Dulbecco's Modified Eagle Medium (DMEM) supplemented with 10% (v/v) fetal calf serum, 2 mM L-glutamine and 110 mg/l sodium pyruvate. All cells were PPLO negative during cultivation. Cells were routinely split 1/20 with trypsin/EDTA every 3 days or when reaching max 80% confluency.

The cell lines HepG2, MCF7-AZ, GM12787 and SK-N-SH were obtained from the VIB IRC cell bank (HepG2, MCF7-AZ, and SK-N-SH) or from the Coriell Institute (GM12787). All cells were PPLO negative throughout cultivation and were grown without antibiotics at 37°C in 5% $CO_2$ humidified incubators. Cells were split when reaching 70% confluency (HepG2, MCF7-AZ, SK-N-SH_RA) or when reaching max. 1 million cells/ml (GM12878). HepG2 cells were grown in Dulbecco's Modified Eagle Medium (DMEM) supplemented with 10% (v/v) fetal calf serum (FCS), 2 mM L-glutamine, 10 mM sodium pyruvate and 100 μM non-essential amino acids. MCF7-AZ, G12878 and SK-N-SH_RA cells were grown and propagated according to the UW ENCODE Cell culture SOPs (http://genome.cse.ucsc.edu/ENCODE/protocols/cell/human/Stam_15_protocols.pdf). For MCF7-AZ, this was in Eagle's Minimal Essential Medium with 10% FCS, 2 mM L-glutamine, and 100 μM non-essential amino acids; splitting with Accutase (Thermo Fisher). For GM12878, this was in RPMI 1640 with 2 mM L-glutamine, 15% FCS. For SK-N-SH_RA, undifferentiated SK-N-SH cells were grown in RPMI 1640 with 2 mM L-glutamine, 10% FCS, and 10 mM sodium pyruvate and split with Accutase (Thermo Fisher). Prior to harvesting, cells were treated for 48h with medium containing 6 μM *all-trans* retinoic acid for differentiation to cells with a neural phenotype.

##### Human cDNA fragment library construction

Human cell line total RNA was isolated using the Innuprep RNA MIDI Direct kit (Analytik-Jena) according to the manufacturer's instructions, additionally digesting potentially remaining genomic DNA with DNase (Turbo DNA-free kit, Ambion) for 1h at 37°C. RNA integrity was checked on the Agilent BioAnalyzer; all samples always had a RIN of 9 or higher. For the library screen in *P. pastoris*, samples from the HepG2, MCF7-AZ, GM12878 and SK-N-SH_RA cell lines were pooled in equal amounts. Next, poly-A$^+$ transcripts were selected with the Oligotex mRNA midi kit (Qiagen) and precipitated overnight at -20°C in 100% RNase-free EtOH (3x initial volume) with RNase-free NaOAc pH 5.2 (0.3 M final) containing RNase-free glycogen (100 ng/μl final). Poly-A$^+$-selected RNA was recovered by pelleting for 1h at 4°C at 14000 x *g*, washing with 70% RNase-free EtOH, and resuspended in RNase-free water (Ambion). Samples were further depleted of ribosomal RNA with the Ribo-Zero Gold (human/mouse) magnetic kit (Epicentre) following the manufacturer's instructions but using up to 7.5 μg of polyA$^+$ RNA per reaction. Ribodepleted samples were then purified with the RNeasy MinElute Cleanup kit (Qiagen). The RNA was furthermore diluted to 37.5 ng/μl in 16 μl reactions, and fragmented with 1.8 μl of $Zn^{2+}$ fragmentation buffer (100 mM $ZnCl_2$ in 100 mM Tris-HCl pH 7.0) in a PCR machine with heated lid at 70°C for 1 min 45s. These conditions were optimized to yield fragments with a Poisson-distributed length around 500 bp. Fragmentation was stopped with 1.8 μl 0.5 M EDTA pH 8.0, and samples were pooled and purified once more with the RNeasy MinElute Cleanup kit (Qiagen). RNA quality and size distribution was monitored at each step on a 2100 BioAnalyzer using RNA 6000 pico chips (Agilent Technologies).

In subsequent steps, contamination with environmental human genomic DNA was avoided as much as possible until after the adaptor ligation step. Fragmented RNA was transcribed to double stranded cDNA using the Maxima H minus Double-Stranded cDNA synthesis kit (Thermo Fisher Scientific) according to the manufacturer's instructions, but swapping the first strand random primer for our nuclease-protected PacI-tagged random primer (primer A196). After RNase treatment, the cDNA was purified using RNase-free DNA cleanup beads (either AMPure XP beads (Agencourt) or CleanPCR beads (CleanNA), following manufacturer's instructions) with a 1.6:1 ratio beads:sample (v/v). The cDNA was G-tailed using Pyrophage 3137 DNA polymerase exo minus (Lucigen) in a reaction with 0.2 mM dGTP and corresponding Pyrophage polymerase buffer for 30 min at 70°C. After DNA cleanup with beads (1.8x volume), G-tailed cDNA was ligated to the SfiI-adaptor (A188_F and A188_R) in 1x Rapid Ligation buffer and 30 U/μl of T4 UltraPure DNA Ligase (Enzymatics), using 100 pmoles of adaptor per 60 μl reaction, for 15 min at room temperature. Samples were purified twice in DNA cleanup beads (1.6x volume). Before normalization, samples were PCR amplified using primer A141_F (final 600 nM), which hybridizes to the adaptor, and 1x KAPA HiFi HotStart mix (KAPA Biosystems) by denaturation for 3 min at 95°C, and 20 cycles of 98°C for 20s, 67°C for 15s, 72°C for 30s. Samples were purified with DNA cleanup beads (1.6x) and normalized with the Kamchatka crab duplex specific nuclease (DSN) (Evrogen) as in Bogdanov *et al.*[2]. Briefly, per 4 μl reaction, 200 ng of cDNA is mixed with DNase-free water and 1x hybridization buffer (4x stock: 200 mM HEPES pH 7.5 with 2 M NaCl), denatured for 2 min at 98°C, and allowed to hydridize for 5h at 68°C in a PCR machine with heated lid. Avoiding sample cooling, the cDNA is combined with 5 μl of pre-heated 2x DSN Master buffer (Evrogen) and equilibrated at 68°C for 10 min, after which 0.5 μl (1 DSN unit) of DSN enzyme is added, digestion then proceeds for 25 min at 68°C. The reaction is stopped through the addition of 10 μl of preheated 2x EDTA stop solution (Evrogen), and after a brief incubation for 5 min at 68°C, the sample is diluted with 20 μl of

DNase-free water. The single-stranded sample is then PCR amplified using 10 μl of template per 50 μl reaction with 1x KAPA HiFi HotStart mix and primer A141_F (final 600 nM) (3 min at 95°C, and 15 cycles of 98°C for 20s, 67°C for 15s, 72°C for 30s). A second round of normalization is performed after sample cleanup using beads (1.6x), using the same protocol (hybridization + DSN digest + PCR + bead cleanup) but allowing hybridization for 15h and overlaying the hybridization reaction with 10 μl of mineral oil to counter evaporation. cDNA library size distribution was monitored at each step of the procedure on a 2100 BioAnalyzer using DNA high Sensitivity chips (Agilent Technologies). Normalization efficiency was controlled by qPCR comparing the levels of a set of reference genes with various expression levels (*GAPDH* (B002 primers), *RPL13A* (B005), *HMBS* (B003), *HPRT1* (B004), *TBP* (B009), *PIAS1* (B012), *STIM1* (B013), and *ALDH4A1* (B014)) in non-normalized, single-round normalized, and two-round normalized samples. All samples including controls were diluted to 5 ng/μl in DNase-free water, with final 10 μl qPCR reactions containing 2.5 ng DNA, 1x SensiFast SYBR No-ROX qPCR mix (Bioline), 300 nM forward primer and 300 nM reverse primer. Reactions were run on a LightCycler 480 (Roche) with 3 min denaturation at 95°C, followed by 45 cycles of 95°C for 3s, 65°C for 30s (ramp rate 2.5°C/s), and 75°C for 1s. Melting curves were generated to check the specificity of the reactions.

**Human cDNA library cloning and plasmid library preparation**

The cDNA fragment libraries were cloned in the *S. cerevisiae* pSSDSfiIPacI-FLAGV5-Gal1 and *P. pastoris* pPSDZeoSfiIPacI-FLAGV5-AOX1 surface display vector (for the *S. cerevisiae* and *P. pastoris* screens, resp.) using SfiI/PacI restriction digestion and ligation on a preparative scale. 200 μg of vector was first digested overnight at 50°C with SfiI (NEB) in CutSmart buffer (NEB) and an equal molar amount of SfiI-site containing oligo (A136) according to the manufacturer's protocol, in 50 μl aliquots. After cooling to room temperature, PacI (NEB) was added and digestion was allowed to proceed for 1h at 37°C. The backbone band was purified from agarose gel, and dephosphorylated for 1h at 37°C using a thermolabile alkaline phosphatase FastAP (Thermo Scientific) that was inactivated at 75°C for 5 min after dephosphorylation. The cDNA library was also digested sequentially with SfiI and PacI, without A136 oligo, and purified with the NucleoSpin kit (or DNA Clean and Concentrator 500 kit (ZymoResearch) for larger scale purifications) and desalted using CleanPCR beads. Digested library and dephosphorylated vector were combined in a 20:1 molar ratio for ligation with T4 DNA ligase (Thermo Scientific) using the provided T4 Ligase buffer (which was aliquoted to avoid multiple freeze-thaw cycles), aliquoted in 50 μl reactions in a PCR plate, overnight at 16°C in a PCR machine with cooled lid. Prior to electroporation, the reactions were pooled, purified over 1.4x CleanPCR beads, eluted in purified water (3/8ths the original ligation reaction volume), and kept on ice until electroporation.

For electroporation, freshly streaked *E. coli* MC1061 (*S. cerevisiae* screen) or TOP10 (*P. pastoris* screen) cells were grown in 5 ml of liquid LB medium (5 g/l bacto yeast extract, 10 g/l bacto tryptone, 10 g/l NaCl) at 37°C for 1 day. The stationary culture was inoculated the following morning 1/100 in fresh LB in shake flasks of appropriate size for proper aeration, and grown while shaking at 37°C until an $OD_{600}$ of 0.5 (about 2h). The culture was chilled on ice for at least 30 min, pelleted for 15 min at 4000 x *g* at 4°C and washed twice with ice-cold sterile water (first using 1x the original culture volume, then

1/2x), each time pelleting for 15 min at 4000 x *g* at 4°C. A last wash was done in 1/50$^{th}$ of the original culture volume of ice-cold sterile 10% glycerol, to resuspend the now electrocompetent cells in ice-cold sterile 10% glycerol (600 µl per 200 ml of starting culture). Electroporation was performed in pre-chilled 96-well electroporation plates (HT-200 system from BTX), using 40 µl electrocompetent cells with 2.5 µl of purified ligation reaction per well (mix well), with the Gene Pulser electroporation system (BioRad) set at 200 Ω, a capacitance of 25 µF, a capacitance extension of 125 µF, and a voltage of 2.5 kV. Cells were immediately transferred and pooled in SOC medium (5 g/l bacto yeast extract, 20 g/l bacto tryptone, 0.5 g/l NaCl, 2.5 mM KCl, 10 mM $MgCl_2$, 20 mM dextrose set to pH 7.0) at 1 ml SOC per reaction, and allowed to recover for 1h at 37°C. A serial dilution of these recovered cells was plated on agar plates with the appropriate antibiotic to assess transformation efficiency, and the rest of the culture was spread on large agar + antibiotic 24.5 cm x 24.5 cm bioassay dishes (3-4 ml per dish) using plastic sterile drigalski spatulas. After 16-24h growth in a 37°C incubator, all the colonies were scraped from the agar and pooled. The pellet was washed with sterile water, and weighed to assess cell number and the appropriate plasmid extraction scale, as described in the manual of the plasmid extraction kit used. The plasmid library was then extracted from the bacterial cells using one or multiple NucleoBond Xtra Midi preps (Macherey-Nagel) or QIAfilter Plasmid Giga preps (Qiagen) and eluted in Tris-HCl pH 8.5. The QIAfilter Giga preps give the overall best yield and purity. All reactions and electroporations were scaled or repeated as necessary.

Library diversity was estimated assuming equally probable variants as described in Bosley *et al.*[3], which states that the diversity $D = D_{max}*(1-e^{-T/D_{max}})$ with $D_{max}$ being the maximal diversity (given an infinite number of transformants), and T the number of transformants obtained. Note that this number does not reflect the *probability* that a randomly picked fragment is present in the library, nor does it reflect the completeness of the library, but merely the maximal diversity possible given a particular number of transformants. In the case of our human cDNA fragment libraries, we approximate $D_{max}=5*10^7$ (assuming a normalization factor of 1024 and based on a 100 bp resolution). Note that $D_{max}$ is larger in reality as fragmentation is random. For the *S. cerevisiae* screen, we obtained an estimated $2.66*10^6$ *E. coli* transformants (transformation efficiency $1.21*10^5$ CFU/µg vector DNA) collected from 72 large agar dishes after 216 transformation reactions, and thus calculate a diversity of $2.59*10^6$ plasmid clones. For the *P. pastoris* screen, we obtained a total of approximately $1,28*10^7$ *E. coli* transformants (transformation efficiency on average around $10^5$ CFU/µg vector DNA used in the ligation reaction) collected from 318 large agar dishes after 1148 transformation reactions, and thus calculate a diversity of $1,13*10^7$ plasmid clones.

##### *S. cerevisiae* library generation

The human cDNA-surface display plasmid library was transformed to *S. cerevisiae* strain R1158 using the large-scale high-efficiency LiAc/SS carrier DNA/PEG heat shock method described in the Nature Protocols paper by Gietz and Schiestl[4] (120x scale). A small fraction of cells was serially diluted, plated and grown on SD-Leu-Ura agar plates at 28°C for 3 days to assess transformation efficiency. The rest of the cells were immediately inoculated 1/20 in liquid SD-Leu-Ura medium (6.7% yeast nitrogen base w/o amino acids, with ammonium sulfate; 2% dextrose; 0.077% CSM-Leu-Ura dropout mix; pH 5.8) in shake flasks of the appropriate size after heat shock, and transformants were selected for 48h at 30°C while shaking. After selection, a small aliquot of cells was serially diluted and plated on YPD plates (10 g/l yeast extract, 20 g/l peptone, 20 g/l dextrose, 17 g/l agar) for colony PCR-based assessment of selection efficiency. The rest of the library was aliquotted and frozen at -80°C in 15% glycerol. Transformations were scaled up or repeated as necessary.

For the library used in this screen, we obtained $3.68*10^6$ yeast transformants (the transformation efficiency was $3.06*10^5$ CFU/µg plasmid DNA), and with a $D_{max}$ of $2.59*10^6$ (the plasmid library diversity), the estimated diversity of this yeast library is thus $1.96*10^6$ clones. As is customary in the field[5–7], to ensure recovery of virtually all clones in downstream steps, we always worked with at least 10x as many cells as the estimated library diversity.

##### *P. pastoris* optimized transformation procedure

Plasmids or plasmid libraries were linearized within the AOX1 promoter with *MssI* (NEB, Ipswich, USA), checked for complete digestion on agarose gel and purified with CleanPCR beads (CleanNA). We modified the high-efficiency *P. pastoris* electroporation protocol as described in Wu and Letchworth[8]. Briefly, cells are grown from subcultures to an $OD_{600}$ of 1.5, pelleted at room temperature at 1500g for 5', and resuspended in 200 ml of sterile LiAc/DTT solution (100 mM LiAc, 10 mM DTT (from fresh 1 M stock), 600 mM sorbitol, 10 mM Tris-HCl pH 7.5) per 250 ml culture. The suspension is incubated for 30' at 28°C with gentle shaking (100 rpm). Pellets (1500 x *g* for 5' at 4°C) are subsequently washed 3 times with ice-cold and sterile 1 M sorbitol (37.5 ml per 250 ml starting culture), and kept on ice as much as possible. The pretreated cells are finally reconstituted in 1 M ice-cold sorbitol (1.875 ml per 250 ml starting culture) and kept on ice until electroporation. For electroporation, 80 µl of pretreated *P. pastoris* cells are mixed with 100 ng – 1 µg (range tested during optimization experiments) of desalted, linearized library DNA (reconstituted in MQ) in an ice-cold 0.2 cm electroporation cuvette or electroporation 96-well plate. These mixes are electroporated at 200 Ω, a capacitance of 25 µF and capacitance extension of 125 µF, and a voltage of 1.5 kV using the Gene Pulser electroporation system (BioRad, Hercules, USA), connected to a HT-200 plate handler (BTX, Holliston, USA) for high-throughput electroporations. Immediately after electroporation, 1 ml of ice-cold YPD pH 8.0 is added and cells are transferred to appropriate flasks of tubes. The $OD_{600}$ is measured before and after a 6h recovery with incubation at 28°C while shaking. Cells are subsequently plated onto fresh YPD pH 8.0 agar plates containing 20 µg/ml of zeocin using glass beads to ensure uniform dispersion and grown for 3 days at 30°C. Transformation efficiencies are calculated based on the number of colony forming units per µg of vector DNA, corrected with the factor of growth that occurred during recovery.

##### *P. pastoris* library generation

We transformed the linearized large human cDNA-surface display plasmid library to *P. pastoris* strain GS115 using the optimized library transformation procedure described above, in 184 transformations using 96-well format electroporation cuvettes (BTX) with 1 µg per transformation. A small fraction of cells was serially diluted after electroporation and recovery, and plated and grown on fresh YPD pH 8.0

agar plates containing 20 µg/ml zeocin for 2-3 days at 28°C in order to assess transformation efficiency. The rest of the cells were inoculated 1/25 in liquid YPD pH 8.0 with 20 µg/ml zeocin, and grown at 28°C while shaking for 2 days. In order to determine the fraction of transformed cells, a serial dilution of the selected culture was plated on non-selective YPD plates and grown for 2 days at 28°C for colony PCR. The rest of the cells was stored at -80°C in aliquots with 15% sterile glycerol. Corrected for the 2.74x factor growth occurring during recovery, transformation efficiency was estimated at $1.23*10^5$ CFU/µg DNA, thus obtaining $2,28*10^7$ transformants and an estimated maximal diversity of $9.8*10^6$ clones.

As for the *S. cerevisiae* library, we always worked with at least 10x as many cells as the estimated library diversity.

##### *S. cerevisiae* cell sorting

For the first round of sorting, $6.89*10^7$ library yeast cells were resuscitated from frozen aliquots in 10 ml of SRaf-Leu-Ura (6.7% yeast nitrogen base w/o amino acids, with ammonium sulfate; 2% raffinose; 0.077% CSM-Leu-Ura dropout mix; pH 5.8) and grown for 24h at 28°C while shaking. The control strain with FLAG-V5-Sag1 was inoculated from plate in 5 ml SRaf-Leu-Ura and grown under the same conditions. Expression was induced at $OD_{600}$=5 in 10 ml (library) or 5 ml (control strain) SRaf/Gal-Leu-Ura (6.7% yeast nitrogen base w/o amino acids, with ammonium sulfate; 1% raffinose; 1% ultra-pure galactose; 0.077% CSM-Leu-Ura dropout mix; pH 5.8) for 24h, again at 28°C while shaking. Cell pellets from two 1.5 ml aliquots of induced library culture were stored at -80°C for plasmid extraction. The remaining cells were kept on ice or at 4°C during the entire staining procedure. Cells were first washed 3x in ice-cold wash buffer (PBS+1mM EDTA, pH 7.2 + 1 Complete Inhibitor EDTA-free tablet (Roche) per 50ml buffer, freshly made and filter sterile), each time spinning down at 4°C for 3min at 3000xg, and stained at $OD_{600}$= 4 with mouse monoclonal anti-V5 (1/500, AbD Serotec MCA2892) and/or rabbit polyclonal anti-FLAG (1/200, Sigma-Aldrich F7425) in ice-cold staining buffer (wash buffer + 0.5 mg/ml of Bovine Serum Albumin) on a rotating wheel for 45' at 4°C, aliquoted in 2 ml tubes. Cell aliquots were washed 2x with 2ml ice-cold staining buffer, and secondary staining was done with goat anti-mouse AF647-RPE (1/250, Life Technologies A20990) and/or goat anti-rabbit AF488 (1/500, Life Technologies A11008) and/or anti-mouse IgG microbeads (50 µl per ml of cells, Miltenyi Biotec 130-048-401), on a rotating wheel for 45' at 4°C in the dark. Cells that underwent MACS enrichment were washed 2x in MACS buffer (MACS BSA stock solution (Miltenyi Biotec) 1/20 in autoMACS rinsing solution (Miltenyi Biotec) + 1 Complete Inhibitor EDTA-free tablet (Roche) per 50 ml buffer, freshly made and filter sterile). MACS enrichment was performed according to the manufacturer's protocol on a single LS column. After elution, cells were pelleted for 3 min at 3000xg at 4°C, and recovered in 350 µl staining buffer. Cell samples that were not subjected to enrichment were washed 2x with ice-cold staining buffer. All samples were filtered over 35 µm cell strainer caps before measurement. Flow cytometry and cell sorting was performed on a MoFlo Legacy sorter (Beckman Coulter) accompanied by FlowJo v10.1 for data analysis. Fluorophores were excited at 488 nm, and fluorescence was collected through 605 short pass +530/40 band pass filters (AF488) and/or a 670/30 band pass filter (AF647-RPE). Cells were gated for a uniform SSC vs FSC single-cell population, and fluorescence quadrant gates were chosen as such that, after compensation, max. 5% of cells of unstained and single stained controls appeared above the background. We sorted out roughly 350 000 MACS-enriched $FLAG^{+}V5^{+}$ cells per screen (>10x library diversity was screened), adding 9ml of SD-Leu-Ura + Pen/Strep (6.7% yeast nitrogen base w/o amino acids, with ammonium sulfate; 2% dextrose; 0.077% CSM-Leu-Ura dropout mix; pH 5.8 + 100 U/ml penicillin and 100 µg/ml streptomycin (Thermo Fisher Scientific)) to the collected cells for recovery. Sorted cells were then grown for 3 days at 28°C while shaking, and frozen at -80°C in 15% glycerol aliquots.

For the second round of sorting, round 1 sorted cells and control strains were grown, induced, stained and sorted as in the first round but omitting MACS pre-enrichment and choosing a slightly more stringent gate to increase specificity. Cells were recovered for 4 days, part of the culture was frozen as slants at -80°C in 15% glycerol aliquots, and part of it was frozen as pellets for plasmid DNA isolation. A dilution series of these round 2 sorted cells was plated out on SD-Leu-Ura agar plates (SD-Leu-Ura + 1.7% agar) for single clone analysis. Purity of the two-round sorted cells was verified by growing +/- $2,5*10^{7}$ cells in 20 ml SRaf-Leu-Ura + Pen/Strep (100 U/ml penicillin and 100 µg/ml streptomycin) for 48h at 28°C while shaking, and again inducing expression at at $OD_{600}$=5 in SRaf/Gal-Leu-Ura + Pen/Strep for 24h at 28°C while shaking. Cells were stained as described for the first and second sorting round, data was again collected on the MoFlo Legacy flow cytometer, and analyzed using FlowJo v10.1. The entire sorting of this yeast library was replicated three times on separate days.

##### *P. pastoris* cell sorting

For the sorting of protein fragment displaying *P. pastoris* cells, $2.2*10^{8}$ library yeast cells were resuscitated from frozen aliquots in 100 ml of buffered complex glycerol medium (BMGY) (10 g/l bacto yeast extract, 20 g/l bacto peptone, 100 mM potassium phosohate buffer pH 6.0, 1.34% yeast nitrogen base with ammonium sulfate; $4*10^{-5}$% biotin, 1% glycerol) and grown for 24h at 28°C while shaking. The control 'empty vector (EV)' strain with FLAG-V5-Sag1 was inoculated from plate in 5 ml of BMGY and grown under the same conditions. Expression was induced at $OD_{600}$=10 after switching the medium to buffered complex methanol medium (BMMY) (10 g/l bacto yeast extract, 20 g/l bacto peptone, 100 mM potassium phosohate buffer pH 6.0, 1.34% yeast nitrogen base with ammonium sulfate; $4*10^{-5}$% biotin, 1% methanol), in 25 ml for the libraries and 5 ml for the control strain. Induction was allowed for 48h at 28°C while shaking, spiking in methanol to 1% every 8-12h. At this point, a few ml of culture was subjected to genomic DNA extraction for downstream sequencing using the MasterPure Yeast DNA purification kit (Epicentre) following the manufacturer's instructions. The remaining cells were then stained, keeping samples on ice or at 4°C during the entire procedure. Cells were first washed 3x in ice-cold wash buffer (PBS+1mM EDTA, pH 7.2 + 1 Complete Inhibitor EDTA-free tablet (Roche) per 50ml buffer, freshly made and filter sterile), each time spinning down at 4°C for 3 min at 1500 x g, and stained at $OD_{600}$=2 with mouse monoclonal anti-V5 (1/500, AbD Serotec MCA2892) and/or rabbit polyclonal anti-FLAG (1/200, Sigma-Aldrich F7425) in ice-cold staining buffer (wash buffer + 0.5mg/ml Bovine Serum Albumin) on a rotating wheel for 45' at 4°C. Cells were washed 2x with ice-cold staining buffer, and secondary staining was done with goat anti-mouse AF647-RPE (1/250, Life Technologies A20990) and/or goat anti-rabbit AF488 (1/500, Life Technologies A11008) and/or anti-mouse IgG MACS microbeads (50µl per ml cells, Miltenyi Biotec 130-048-401), on a rotating wheel for 45' at 4°C in the dark. Cells that underwent MACS enrichment were washed 2x in MACS buffer (MACS BSA stock solution (Miltenyi Biotec) 1/20 in autoMACS rinsing solution (Miltenyi Biotec) + 1 Complete Inhibitor EDTA-free tablet (Roche) per 50ml buffer, freshly made and filter sterile). MACS enrichment was performed according to the manufacturer's protocol on two LS columns. After elution, cells were pelleted for 3 min at 1500 x *g* at 4°C, and recovered in 2.5 ml of staining buffer. Cell samples that were not subjected to enrichment were washed 2x with ice-cold staining buffer. All samples were filtered over 35 µm cell strainer caps before measurement. Flow cytometry and cell sorting was performed on a MoFlo Legacy sorter (Beckman Coulter) accompanied by FlowJo v10.1 for data analysis. Fluorophores were excited at 488 nm, and fluorescence was collected through 605 short pass +530/40 band pass filters (AF488) and/or a 670/30 band pass filter (AF647-RPE). Cells were gated for a uniform SSC vs FSC single-cell population, and fluorescence quadrant gates were chosen as such that, after compensation, max. 5% of cells of unstained and single stained controls appeared above the background. We sorted out approximately 5 million MACS-enriched $FLAG^{+}V5^{+}$ cells per screen (a number of events >10x library diversity was screened in total). Sorted cells were spun down at 1500 x g for 5 min at 4°C, and recovered in 20 ml of YPD pH 8.0 + Pen/Strep (100 U/ml penicillin and 100 µg/ml streptomycin (Thermo Fisher Scientific)). After 12h, zeocin was added to 20 µg/ml. Sorted cells were grown for 36h in total at 28°C while shaking, and frozen at -80°C in 15% glycerol aliquots. For genomic DNA isolation, cells were recovered in YPD pH 8.0 with Pen/Strep and zeocin, and genomic DNA was extracted using the MasterPure Yeast DNA purification kit. Library sorting was replicated 3 times on three different days.

##### *S. cerevisiae* deep sequencing library preparation

Plasmid isolation of sorted and non-sorted *S. cerevisiae* yeast libraries was performed as in Whitehead *et al.*[6] using the ZymoPrep Yeast Plasmid Miniprep II kit (Zymo Research). Briefly, $9\text{-}20*10^{7}$ pelleted frozen cells were resuspended in 400 µl of Solution I with 50 U Zymolyase and incubated for 4h at 37°C. After a flash freeze in liquid $N_2$ and thawing at 42°C, plasmid extraction was continued as described in the manufacturer's protocol, but eluting in 30 µl of 10 mM Tris-HCl pH 8.0. Genomic DNA was digested with 60 U of exonuclease I (NEB) and 7.5 U lambda exonuclease (NEB) in lambda exonuclease buffer (NEB) for 90 min at 30°C, followed by inactivation for 20 min at 80°C. Library plasmids were purified from the buffer using CleanPCR beads (2x reaction volume) (GC Biotech) and eluted in 22 µl MilliQ water. Next, the human cDNA fragments on the plasmids were recovered by PCR using two pools of 'frameshifting' primers in analogy to Lundberg *et al.*[9], so as to equalize base distribution at the first sequenced positions in order to take maximal advantage of the sequencing chip capacity. Pools of equal molar concentration were made for A247_F*x* and for A247_R*x*. PCR reactions were set up using 20 µl purified plasmid DNA, 300 nM of each primer pool, and 1x KAPA HiFi HotStart Readymix in a final volume of 50 µl, and run for 3 min at 95°C, followed by 25 cycles of 98°C for 20s, 61°C for 15s, 72°C for 30s. Samples were purified using CleanPCR beads (1.6x reaction volume) and eluted in 40 µl of 0.1x TE buffer (1mM Tris-HCl + 0.1mM EDTA, pH 8.0). Illumina adaptor sequences and barcodes were added using the NEBNext Ultra DNA library prep kit for Illumina (NEB) largely according to the manufacturer's protocol, except that the samples were purified using two rounds of 1.6x volume

CleanPCR beads after adaptor ligation to remove adaptor dimers, and that the final PCR was performed with custom primers (A237_F and A237_R_bcx, with bcx indicating different barcodes), desalted Ultramers from IDT) and for 25 cycles. After PCR, the 500-1200 bp fragments were purified from 2% agarose gel using the Nucleospin gel and PCR cleanup kit (Macherey-Nagel), specifically solubilizing the agarose blocks overnight at 4°C in NT buffer to avoid fragment denaturation and reduce GC-bias. After elution in NE buffer, samples were purified a second time using CleanPCR beads (1.6x volume) and finally eluted in 25 µl of 0.1x TE buffer in DNA LoBind tubes (Eppendorf). Reasoning that the reduced complexity of the sorted fragment pool would require less depth than that of the unsorted fragments, samples were pooled in a 2.5/1 molar ratio of unsorted/sorted libraries. Concentrations were determined using Nanodrop, Qubit, and the KAPA Library Quantification kit for LC480 on an Lightcycler 480 (Roche) according to the manufacturer's instructions. Size distributions were assessed on a 12-capillary Fragment Analyzer (Advanced Analytical) with their High Sensitivity NGS kit (DNF-474, Advanced Analytical), and the BioAnalyzer (Agilent) with the DNA High Sensitivity kit (Agilent).

##### *P. pastoris* deep sequencing library preparation

The cDNA fragments of sorted and unsorted *P. pastoris* library were picked up from genomic DNA by PCR (500 nM A149_F, 500 nM A149_R, 1x KAPA HiFi HotStart master mix, 70 ng genomic DNA per 20 µl reaction – 95°C for 3 min, followed by 20 cycles of 98°C for 20s, 61°C for 15s, 72°C for 30s before cooling). PCR fragments between 300-1000 bp in length were isolated from a 2% agarose gel using the NucleoSpin Gel and PCR cleanup kit (Macherey-Nagel) and CleanPCR beads (CleanNA), solubilizing the plugs at 4°C to avoid denaturation of AT-rich fragments, eluting in 30 µl purified water. This pool of fagments was then further subjected to a second short PCR for the addition of frameshifting bases (500 nM A247_F primer pool, 500 nM A247_R primer pool, 1x KAPA HiFi HotStart master mix, 20 µl DNA per 50 µl reaction – 95°C for 3 min, followed by 5 cycles of 98°C for 20s, 61°C for 15s, 72°C for 30s before cooling) and was purified with CleanPCR beads (1.6:1 ratio beads:reaction volume) and eluted in 45 µl of purified water. Illumina sequencing library construction was done with the NEBNext Ultra DNA library prep kit (NEB) largely according to the manufacturer's protocol, except that the samples were purified using one rounds of 1.2x volume CleanPCR beads after adaptor ligation to remove adaptor dimers, and that the final PCR was performed with custom primers (A237_F and the barcoded A237_R_bcx, desalted Ultramers from IDT) and for 7 cycles. This number of PCR cycles was found to be optimal after a prior optimization experiment in which we followed the PCR reactions in real time in a qPCR with SYBR Green, to determine the maximal number of cycles until an amplification plateau is reached. Fragments were purified using CleanPCR beads (0.7x volume) and finally eluted in 25 µl of 0.1x TE buffer in DNA LoBind tubes (Eppendorf). To increase sample yields, we did an additional 4-cycle PCR with primers against the P5 and P7 sequences (500 nM A240_F, 500 nM A240_R, 1x KAPA HiFi HotStart master mix, 2.5 µl DNA per 100 µl reaction - 95°C for 3 min, followed by 4 cycles of 98°C for 20s, 63°C for 15s, 72°C for 30s before cooling). Fragments were again purified using CleanPCR beads (0.7x volume) and eluted in 30 µl 0.1x TE buffer in DNA LoBind tubes (Eppendorf). Samples were pooled in a 4.3/1 molar ratio of unsorted/sorted libraries. Concentrations were determined using Nanodrop, Qubit, and the KAPA Library Quantification kit for LC480 on an Lightcycler 480 (Roche) according to manufacturer's instructions. Size distributions were assessed on a 12-capillary Fragment Analyzer (Advanced Analytical) with their High Sensitivity NGS kit (DNF-474, Advanced Analytical).

##### Illumina sequencing, read processing, and sequencing data analysis

For each screen, the pooled sample was paired-end sequenced (2x150bp) on an Illumina NextSeq 500 mid-throughput or high-throughput (*S. cerevisiae* or *P. pastoris* screen, resp.) chip. Raw demultiplexed Illumina sequencing data was processed using a combination of publicly available tools and custom scripts. Raw reads were first trimmed with Trim Galore! version 0.4.1 (www.bioinformatics.babraham.ac.uk/projects/trim_galore) to remove Illumina adapter sequences. Next, FLAG/V5 and frameshifting sequences were trimmed off with Cutadapt version 1.10 (ref. 10), discarding all untrimmed pairs to only keep correctly cloned cDNA fragments. Quality control of raw and processed fastq files was performed using FastQC version 0.11.3 (www.bioinformatics.babraham.ac.uk/projects/fastqc). Processed reads were mapped to the human transcriptome of known protein-coding genes as downloaded from Ensembl's BioMart[11] using BBMap v35.40 (sourceforge.net/projects/bbmap). Count tables were built and analyzed from the properly paired mapped reads using SAMtools[12] v1.2 and v1.3, BEDtools[13] v2.24.0 and v2.25.0, EMBOSS[14] v6.6.0, R project 3.3.0 (www.R-project.org) and the R packages plyr, ggplot2, alakazam, stringr, and UpSetR[15]. A summary of the most important scripts can be found on Figshare (figshare.com/s/5dba6b512fa74ef68a40). Fragments were considered detected when fragment count > 0 in either the unsorted sample, or the sorted sample. Enrichment factors (E factors) were calculated as $\log_2\left(\frac{FPTM\ sorted}{FPTM\ unsorted}\right)$, with FPTM being our custom Fragment count Per Ten Million fragments which is defined as the number of read pairs with the same start and end position per 10 million read pairs. For the concordance calculations, *FPTM unsorted* was calculated over the merged replicate unsorted samples, and from the fragments detected in all 3 replicates (sorted sample or merged unsorted), only the fragments that were in-frame with both the N-and C-term fusion parts in the surface display construct were considered (as we used random priming, there is an expectable 1/9 chance that a cloned fragment is in the same reading frame with both the N-and C-term fusion parts).

##### Flow cytometry of randomly picked sorted *S. cerevisiae* clones

To assess the correlation between sequencing count and surface display fluorescence signal, 47 two-round sorted *S. cerevisiae* single clones and the control strain with FLAG-V5-Sag1 were inoculated in 2 ml of SRaf-Leu-Ura in deep 24-well plates and grown for 24h at 28°C while shaking. Cells were pelleted at 4°C at 3000 x *g* for 3 min, supernatans was removed, cells were resuspended in 2 ml SRaf/Gal-Leu-Ura, and induced for 24h at 28°C. Cell staining was performed as done for the library round 2 sorts, without MACS enrichment, but working in 96-well V-bottom Nunc microwell plates (Thermo Fisher). Cells were finally diluted ¼ in staining buffer, and measured on an LSR-II HTS flow cytometer (BD). Fluorophores were excited at 488 nm, and fluorescence was collected through 550 long pass + 525/50 band pass filters (AF488), and/or 670 long pass + 685/35 band pass filters (AF647-RPE). Compensation, gating and further data analysis were done in FlowJo v10.1.

For identification, the same clones were subjected to a colony PCR targeting the human cDNA fragment that each clone encodes. For each clone, a single colony was picked from plate and resuspended in 20 µl of freshly-made 20 mM NaOH and incubated for 5 min at room temperature (RT). Lysis was stopped by adding 80 µl of sterile water, and 5 µl was used in a 25 µl PCR reaction with 0.5 U Phusion High Fidelity polymerase (NEB), 500 nM of forward primer A207_F, 500 nM of reverse primer A221_R, 1x Phusion HF buffer, and 200 µM dNTPs (Promega). PCR cycling conditions involved a 98°C denaturation for 30s; 30 cycles of 98°C for 10s, 52°C for 15s, 72°C for 45s; finishing off with 5 min at 72°C before cooling. PCR fragments were purified by CleanPCR beads (GC Biotech), Sanger sequenced from both ends using primers A149_F and A149_R, and obtained sequences were mapped to the human reference transcriptome sequences using BLAST and reconstructed from there. NGS fragment counts in background and sorted cell libraries were obtained by searching the count tables for fragments with the same gene symbol and amino acid sequence.

##### Secretion validation via western blot

For the validation of fragment secretability, 20 random single *S. cerevisiae* clones from the replicate 1 screen sorts were grown in 2 ml of SD-Leu-Ura, and plasmids were isolated with the ZymoPrep Yeast Plasmid Miniprep II kit (Zymo Research) according to the manufacturer's instructions. For each clone, the encoded cDNA fragments were isolated via PCR with 300 nM of A262_F and 300 nM of A262_R primers (with homologous overhangs for downstream cloning in pSCA-stuffer), in a 1x KAPA HiFi PCR reaction using 4 ng of plasmid DNA per 20 µl reaction. Samples were denatured for 3 min at 95°C, and cycled 25x at 98°C for 20s, 57°C for 15s, 72°C for 20s, before cooling. Amplified DNA was extracted from gel and purified. Secretion vector pSCASfiIPacI-FLAGV5-AOX1-stuffer was digested with SfiI and PacI (both NEB), and the vector backbone was also isolated from gel and purified. Fragment and backbone were assembled using Gibson Assembly for 30 min at 50°C, and transformed to *E. coli*. Plasmids were verified by sequencing. These 20 pSCA-fragment vectors were transformed to yeast using the LiAc/PEG method by Gietz and Schiestl[17], transformed clones were checked by colonyPCR as described above, but using primers A221_R and A221_F. For secretion induction, these single yeast clones were first grown in SRaf-Leu-Ura for 48h at 28°C while shaking, pelleted, and induced in SRaf/Gal-Leu-Ura for 24h at 28°C. Medium was collected and frozen at -20°C until protein extraction.

Secreted proteins were pelleted from the medium by precipitation with DOC and TCA. Briefly, for each sample, 10% of the sample volume of 5 mg/ml deoxycholate (DOC) was added, the sample was incubated on ice for 10 min, 13.54 M trichloroacetic acid (TCA) was added at 10% sample volume, the sample was incubated on ice for 20 min, and the precipitate was pelleted at 4°C in a centrifuge at max. speed for 30 min. Supernatans was removed, and the pellet was washed twice with ice-cold acetone, and once with 70% ethanol, each time pelleting the sample for 20 min at 14 000 x *g* at 4°C. The pellets were dried at 37°C and resuspended in 1x phosphate buffered saline PBS. Total protein concentration was estimated with the microBCA kit (Pierce) according to the manual's instructions. For each sample, 10 µg of protein was additionally PNGase F digested (NEB) overnight, according to the manufacturer's protocol. Finally, equal amounts of protein for each sample were denatured in 1x Laemmli buffer (10%

glycerol, 0.1% DTT, 63 mM Tris-HCl pH 6.8, 2% SDS, 0.0005% bromophenol blue) for 10 min at 98°C, run on a 15% Tris-Glycine SDS-PAGE gel, and semi-dry blotted for 1h30 on PVDF membranes at 75 mA per 45 $cm^2$ blot. Blots were blocked with 3% milk powder solution for 2h at RT or 4°C overnight and stained with polyclonal rabbit anti-FLAG antibody (1/2000, Sigma, F7425) + anti-rabbit IgG-Dylight800 antibody (1/15000, Thermo Scientific, #35571), or mouse anti-V5 monoclonal antibody (1/3000, AbD Serotec, #MCA1360) + anti-mouse IgG-Dylight8000 (1/15000, Thermo Scientific, #35521). The ladder was the BioRad Precision Plus Dual Xtra ladder. Blots were imaged with the Li-Cor Odyssey system.

##### Feature enrichment analysis

Protein and fragment structural disorder prediction was done using RAPID[16]. To assess whether secretable fragments were more likely to be derived from endogenously secreted proteins than by chance, human proteins and human secretory proteins (ie with signal peptide) were downloaded from Uniprot (release 2018_11) and intersected with the lists of secretable and depleted fragments. Only proteins for which no depleted fragments were found were retained for analysis. For analysis of N-glycan sequons, we evaluated the presence of the sequon NXS or NXT but not NPS or NPT using custom awk code.

##### Structural bioinformatics

For the biophysical predictions, sequences were first filtered for a 100% sequence match to the UniProt protein and a length longer than 30 amino acids. Secondary structure (a-helix, b-sheet and random coil) and early folding propensities were predicted as described for EFoldMine[21], but only retaining residues in the full protein sequence that are unambiguously 'secreted' or 'depleted' across overlapping fragments. From this, contiguous regions of secreted or depleted residues were assembled into consolidated fragments, onto which the average of all predictions for that fragment from the original fragment is condensed. Backbone dynamics of sequenced fragments were predicted using Dynamine[19,20]. Plots were generated in R (www.R-project.org) using custom Python scripts.

For PDB mapping, protein fragment sequences were first clustered into representative fragments using the CD-HIT package[18] with an identity parameter of 100%. This clusters all shorter sequences with fully overlapping longer sequences into a single longer representative fragment. The representative fragments from each dataset were used as queries to perform a blast against PDB database using standalone blast (ncbi-blast-2.6.0+). The percentage of secondary structural elements for each fragment with a PDB hit was calculated from its corresponding DSSP coordinates. Domain architectures (Pfam and Gene3D) were retrieved using InterProScan[22] (v 5.24-63.0). Frequency of a particular domain in a dataset was obtained by removing duplicate entries (if a particular domain is present more than once for a particular fragment) in the dataset.

##### Statistical hypothesis testing

Replicability was assessed through calculation of Spearman correlation factors for single-replicate enrichment factors. Hypothesis testing of feature distributions in enriched vs depleted fragments was carried out using non-parametric two-sided Mann-Whitney-Wilcoxon tests. For endogenous secretory protein enrichment, we used a Fisher's One-Sided Exact test. In case of more than 10 comparisons the significance of p-values was corrected using Benjamini-Hochberg multiple hypotheses testing. All analyses were performed using the R programming language (www.R-project.org), except for the correlation calculations of flow cytometric median fluorescence intensity vs enrichment factors of single clones, which were calculated using GraphPad Prism 7.

##### Datasets for binary classification

Two machine learning approaches were explored to investigate to what extent secretable and non-secretable fragments can be distinguished by primary sequence: a gradient boosted decision tree[23] model, and a convolutional neural network model[24]. Both approaches were constructed to perform this binary classification task, and were trained and evaluated on the same datasets.

The *S. cerevisiae* and *P. pastoris* datasets contain 148,156 and 151,761 protein fragments respectively, of which the (non-)secretability was consistent across the three replicates of the sorting experiment. In *S. cerevisiae* a total of 11,625 fragments were found to be secreted (or enriched), and 136,531 fragments were found to be non-secreted (or depleted). *For P. pastoris*, 10,404 secreted and 141,357 non-secreted fragments were found. Furthermore, we only retained fragments with a sequence length of at least 50 amino acids in the dataset, as we consider shorter sequences irrelevant because they do not fold properly. This resulted in the final dataset properties as shown in **Supplementary Table 14**.

Due to the imbalance between positive and negative samples in the dataset, the performance of the models was evaluated using the area under the curve of the receiver operating curve (AUROC) metric, as it is relatively insensitive to changes in class distribution. Instead of working with a fixed class probability threshold, the AUROC takes the ratio of detected enriched fragments (true positive rate) against the ratio of correctly assigned depleted fragments (false positive rate) for all possible thresholds. The AUC of this curve determines the performance, where a value of 1.0 indicates the best achievable performance and random prediction achieves a value of 0.5. The AUROC can also be seen as the probability that a randomly sampled enriched fragment has a higher predicted value than a randomly sampled depleted fragment.

A 10-fold cross-validation (CV) scheme was deployed to calculate the performance over the full dataset. To avoid bias between training and test data during this CV, folds were constructed in a way that all fragments originating from one gene belong to the same fold. If this measure would be disregarded, correct predictions on test data might be a result of sequence similarity and the model overfitting on training data, resulting in overly optimistic results. Simultaneously, folds were constructed to maintain similar class distributions.

##### Gradient boosting

The dataset consists of protein fragments of variable length. Traditional machine learning techniques typically rely on equal sized feature vectors and do not support variable input sizes. To overcome this problem, we extracted feature vectors from primary sequence to ensure a fixed size of the feature vector.

Multiple physicochemical properties were considered when extracting the feature vectors. For each property, the data extraction was performed and a separate model was trained. Amino acid scales were collected for the following properties: polarity[25], hydrophobicity[26], average area buried[27], buried residues[28], bulkiness[29], molar refraction[30], recognition factors[31], molecular weight, transmembrane tendency[32], and HPLC[33].

For each property, five groups of features are extracted, resulting in a vector of 40 values for each data sample:

- Relative amino acid frequency (20 features, independent of the property).
- Sequence length (1 feature, independent of the property).
- The values of the property for the first six (at the N-terminus) and the last six (at the C-terminus) amino acids (12 features).
- The average value of the property over the entire sequence (1 feature).
- The average value of that property per region, when dividing each fragment into six equal-length regions (6 features). Shorter sequences will have shorter regions.

We then built a gradient boosting classifier that takes these feature sets as input. One classifier takes the 40 features per protein fragment as input, and produces a probability for the secretability of that fragment. After training a classifier for each property, an ensemble model is constructed, taking the ten probabilities of the individual classifiers as input for a new gradient boosting classifier, which then produces a final probability.

The hyperparameters for the gradient boosting decision trees were determined for each fold of the cross-validation individually, using a randomized search. For this search, the data from the training set in this fold was used. The results for the gradient boosting classifiers are listed in Supplementary Table 15.

**Convolutional neural network**

In recent times, deep learning techniques have been widely adopted in proteomics[34,35,36]. Especially convolutional neural networks (CNN) have been successfully applied in this context, given their ability to be trained end-to-end from primary sequence (preventing the need for manual feature engineering), their ability to learn spatial relations independent of position, and their intuitive way of encoding sequence motifs in the first convolutional layer.

A potential hindrance of the typical CNN architecture is that it expects a fixed-size input to produce a fixed-size output. Given the variability of sequence lengths in the secretability datasets, we explored four strategies to deal with this variation. After a one-hot encoding and three blocks, each consisting of a convolutional layer, rectified linear unit (ReLU) activation function, dropout layer and max pooling layer, the output of the last block is transformed using one of the following methods:

- Global max pooling, being a max pooling operation over the full sequence.
- K-max pooling[37], where the K highest activations are kept (in their respective order) per channel.
- A bidirectional gated recurrent unit (GRU), where the last hidden states of each direction are concatenated.

As a baseline, we also pad the input sequence with zeros until a fixed length is reached, and truncate any proteins that go beyond this length. After doing this, no transformation to a fixed length is necessary anymore. We choose a maximum length of 200 amino acids, as this covers 99.8% of all considered fragments.

Finally, this fixed-size output is followed by a fully-connected layer, which is then connected to an output layer with a single neuron. A sigmoid is used to generate probabilities from the final activation. The final hyperparameters of the architecture were determined using a grid search, and are given in Supplementary Table 16. The results for each architecture are given in Supplementary Table 17.

##### Identifying decisive input features

A challenge for neural networks, and various other machine learning techniques for that matter, is their lack of inherent interpretability. Attribution methods have been developed to combat this issue. Here, we use the integrated gradients[38] method, which is based on the backpropagation algorithm. The principle of backpropagation-based attribution methods is to first do a forward pass through the network, generating an output signal, and to then backpropagate that signal back to the input to see which parts of the input sequence were responsible for that prediction. This yields a so-called attribution (or saliency) map, with a positive or negative contribution per amino acid towards the predicted secretability of the fragment. The magnitude of the contribution indicates how strongly it directs the the network towards secretable (positive contribution) or non-secretable (negative contribution) prediction. The overall magnitude of contributions scales with the confidence of the model.

For each protein fragment in the test set of a given fold, we calculated the attribution map for the optimal model (with global max pooling). To investigate the general behavior of the model, we then aggregated them using two strategies:

- We calculated the average contribution per amino acid, regardless of where in the sequence it occurred.
- We divided each sequence into twenty regions, and calculated the average contribution per amino acid per region. This means that the first region contains the average contribution of amino acids that occurred in the first 5% of their respective sequences, the second region from 5% to 10%, etc.

**Data availability**

Unprocessed fastq files of both screens have been deposited in the Sequence Read Archive (SRA) under SRP094995. Lists of all detected, enriched or depleted fragments can be found on Figshare (figshare.com/s/82bb61370d7024f6fb09 for *S. cerevisiae* screens and figshare.com/s/cace104b0ffc5a57811f for *P. pastoris* screens), as can the lists of Pfam hits for all representative fragments (figshare.com/s/052370ec40154c09fb68), and CATH/Gene3D hits for those fragments mapping to PDB structures (figshare.com/s/5a8ca88d27168243c9fe). The data has been integrated in a webinterface for easy browsing by biologists interested in secretability of fragments of particular proteins of interest. These fragments are visually mapped to the PDB model of the protein's structure, where such structure is available. The web interface which can be accessed via http://iomics.ugent.be/secrify/search.

**Primers used in this study**

*= phosphorothioate

Bold underlined=barcode

| Primer name | Sequence (5'->3') |
|---|---|
| 3' AOX | GCAAATGGCATTCTGACATCC |
| 5' AOX | GACTGGTTCCAATTGACAAGC |
| A135_F | ATGTTAGGCCGAGATGGCCCAGAATTACCCTCCACGTT |
| A135_R | CAGTGCTTAATTAATTCCGGCTCCTATGTTGT |
| A136_F | CATCCAGTGGCCACTTAGGCCAAAGTCCTGA |
| A136_R | TCAGGACTTTGGCCTAAGTGGCCACTGGATG |
| A141_F | CACTAGGCCGAGATGGCC |
| A149_F | ATGACGATAAGGCCGAGAT |
| A149_R | GGAGAGGGTTAGGGATAGG |
| A188_F | CACTAGGCCGAGATGGCC |
| A188_R | GCCATCTCGGCCTAGTGACTGCTT |
| A196 | CTTAGCATTAATTAANNNNNNN |
| A202_F | TCAAGGAGAAAAAACTATATCTAGATGAGATTTCCTTCAATTTT |
| A204_F | CTCGAGAAAAGAGAGGCC |
| A204_R | GACATAACTAATTACATGACTCGATTAGAATAGCAGGTACGACAAA |
| A207_F | GTTCGATTGCCTGTTTCT |
| A217_R | CGCTTCGGCCTCTCTTTTCTCGAGAGATACCCCTTCTTCTTTAG |
| A220_F | ATGACGATAAGGCGAGCGCCAAAAGCTCTTTTATCTCAACC |
| A220_R | GAGCTTTTGGCGCTCGCCTTATCGTCATCGTCTTTGTAAT |
| A221_F | GACATAACTAATTACATGACTC |
| A221_R | TCAAGGAGAAAAAACTATATC |
| A223_F | GATGACGATAAGGCCGAGATGGCCTAAGGACAAGGGAAAACGCAA |
| A223_R | GTTAGGGATAGGCTTACCATTAATTAACCCTCCTCACGTGCTATT |
| A237_F | AATGATACGGCGACCACCGAGATCTACACTCTTTCCCTACACGACGCTCTTCCGATC*T |
| A237_R_BC 1 | CAAGCAGAAGACGGCATACGAGAT**TTACCGACG**GTGACTGGAGTTCAGACGTGTGCTCTTCCGATC*T |
| A237_R_BC 2 | CAAGCAGAAGACGGCATACGAGAT**ATTGGACAC**GTGACTGGAGTTCAGACGTGTGCTCTTCCGATC*T |
| A237_R_BC 3 | CAAGCAGAAGACGGCATACGAGAT**TCGCATGGA**GTGACTGGAGTTCAGACGTGTGCTCTTCCGATC*T |
| A237_R_BC 4 | CAAGCAGAAGACGGCATACGAGAT**AGCGAACCT**GTGACTGGAGTTCAGACGTGTGCTCTTCCGATC*T |
| A237_R_BC 5 | CAAGCAGAAGACGGCATACGAGAT**AGCTTCGAC**GTGACTGGAGTTCAGACGTGTGCTCTTCCGATC*T |

| | |
|---|---|
| A237_R_BC6 | CAAGCAGAAGACGGCATACGAGAT**GTCAGCCGT**GTGACTGGAGTTCAGACGTGTGCTCTTCCGATC*T |
| A240_F | AATGATACGGCGACCACCGA |
| A240_R | CAAGCAGAAGACGGCATACGA |
| A241_F | GCTAAAGAAGAAGGGGTAT |
| A241_R | AGTTTTGTGCTGGTAGTC |
| A247_F1 | GATGACGATAAGGCCGAGAT |
| A247_F10 | CTACTGACTGATGACGATAAGGCCGAGAT |
| A247_F2 | TGATGACGATAAGGCCGAGAT |
| A247_F3 | CTGATGACGATAAGGCCGAGAT |
| A247_F4 | ACTGATGACGATAAGGCCGAGAT |
| A247_F5 | GACTGATGACGATAAGGCCGAGAT |
| A247_F6 | TGACTGATGACGATAAGGCCGAGAT |
| A247_F7 | CTGACTGATGACGATAAGGCCGAGAT |
| A247_F8 | ACTGACTGATGACGATAAGGCCGAGAT |
| A247_F9 | TACTGACTGATGACGATAAGGCCGAGAT |
| A247_R1 | TGGAGAGGGTTAGGGATAGG |
| A247_R10 | GTGACATCCTGGAGAGGGTTAGGGATAGG |
| A247_R2 | CTGGAGAGGGTTAGGGATAGG |
| A247_R3 | CCTGGAGAGGGTTAGGGATAGG |
| A247_R4 | TCCTGGAGAGGGTTAGGGATAGG |
| A247_R5 | ATCCTGGAGAGGGTTAGGGATAGG |
| A247_R6 | CATCCTGGAGAGGGTTAGGGATAGG |
| A247_R7 | ACATCCTGGAGAGGGTTAGGGATAGG |
| A247_R8 | GACATCCTGGAGAGGGTTAGGGATAGG |
| A247_R9 | TGACATCCTGGAGAGGGTTAGGGATAGG |
| A256_F | GGATTACAAAGACGATGACGATAAGGCCGAGATGGCCGATGAGATTTCCTTCAATTTTTACTGCAGTTTTATTCGCATTAATTAATGGTAAGCCTATCCCTAACCCTCTCCTCGGTCTCGATTCTACGTAAGCGGCCGCGAATTAATTCGCCTTAGACATGACTGTT |
| A256_R | AACAGTCATGTCTAAGGCGAATTAATTCGCGGCCGCTTACGTAGAATCGAGACCGAGGAGAGGGTTAGGGATAGGCTTACCATTAATTAATGCGAATAAAACTGCAGTAAAAATTGAAGGAAATCTCATCGGCCATCTCGGCCTTATCGTCATCGTCTTTGTAATCC |
| A257_F | CGTAGAATCGAGACCGAG |
| A257_R | TAATCGAGTCATGTAATTAGTTATGT |
| A262_F | GAGGGTTAGGGATAGGCTTAC |
| A262_R | ACAAAGACGATGACGATAAG |
| B002_F | TGCACCACCAACTGCTTAGC |
| B002_R | GGCATGGACTGTGGTCATGAG |
| B003_F | GGCAATGCGGCTGCAA |
| B003_R | GGGTACCCACGCGAATCAC |
| B004_F | TGACACTGGCAAAACAATGCA |
| B004_R | GGTCCTTTTCACCAGCAAGCT |
| B005_F | CCTGGAGGAGAAGAGGAAAGAGA |
| B005_R | TTGAGGACCTCTGTGTATTTGTCAA |
| B009_F | CGGCTGTTTAACTTCGCTTC |
| B009_R | CACACGCCAAGAAACAGT |
| B012_F | ATTTGCCTTGACACCACAACA |
| B012_R | CCTTAACTGGACCTGTACTGTGA |
| B013_F | AGTCACAGTGAGAAGGCGAC |
| B013_R | CAATTCGGCAAAACTCTGCTG |
| B015_F | TACCAAGTGTCGCCTTTTAACC |
| B015_R | GCTCTTGTCTGCATAACAGAACT |
