## Supplementary Information for "Massively parallel interrogation of protein fragment secretability using SECRiFY reveals features influencing secretory system transit"

7. Present address: Department of Biochemistry and Biophysics, UCSF, San Francisco, CA, USA

* corresponding authors

This pdf file includes

Suppl. Figs 1-18

Suppl. Tables 1-11

Suppl. References

### Supplementary Figures

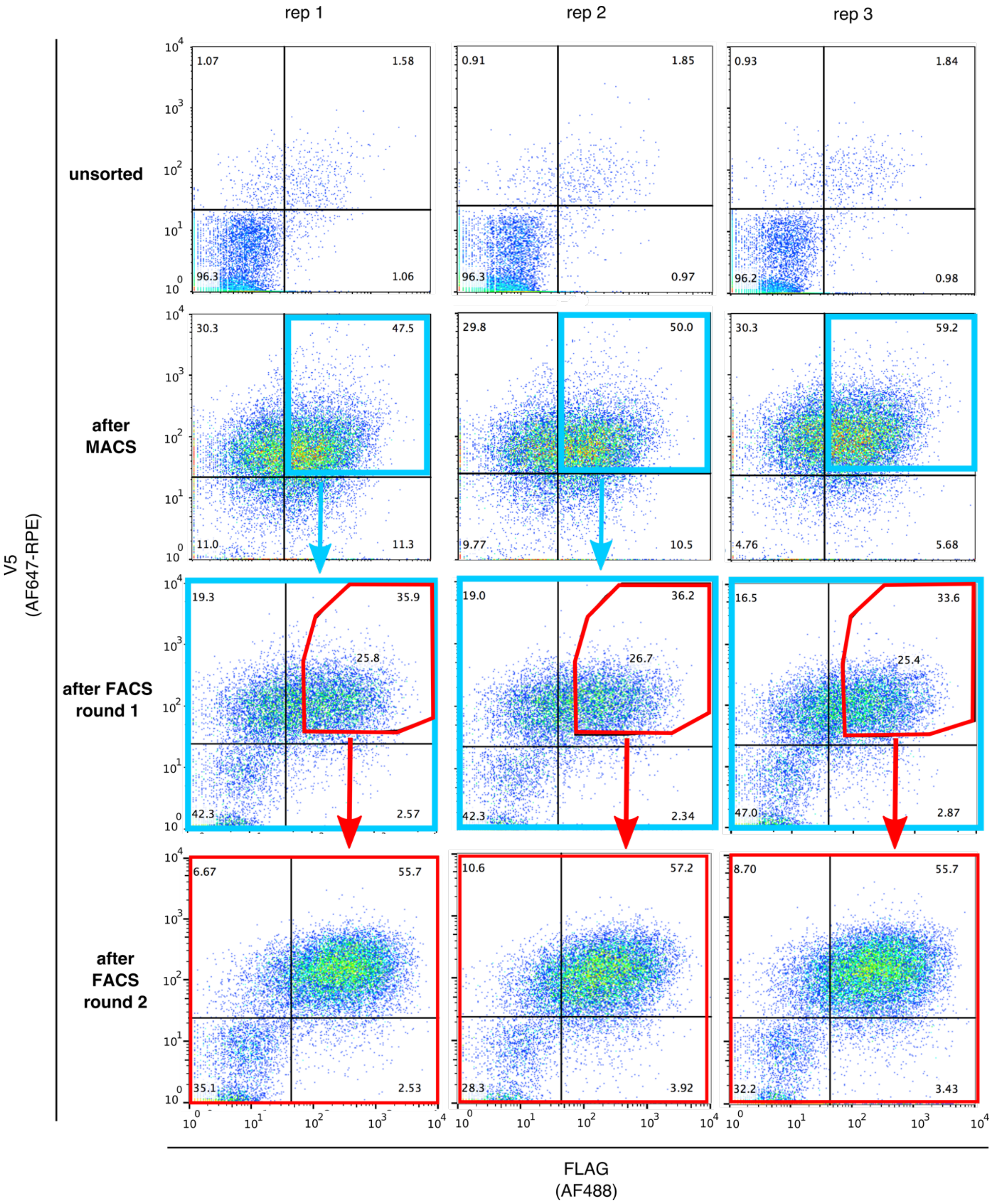


**Supplementary Fig. 1. Sorting of fragment-displaying *S. cerevisiae* library clones.** The *S. cerevisiae* cDNA fragment yeast library was sorted for FLAG$^+$V5$^+$ cells in three replicate experiments (rep 1, rep2 and rep 3). In the first round, V5$^+$ cells were enriched by MACS (row 2, 'after MACS'), and further directly sorted out for FLAG$^+$V5$^+$ cells by FACS. Sorted cells were frozen after recovery, regrown for induction and staining, and subjected to another round of FACS with a slightly more stringent (red) gate (row 3, ‘after FACS round 1’). These round 2 sorted cells were frozen for storage, again regrown for induction and staining, and examined for purity by flow cytometry (row 4, ‘after FACS round 2).

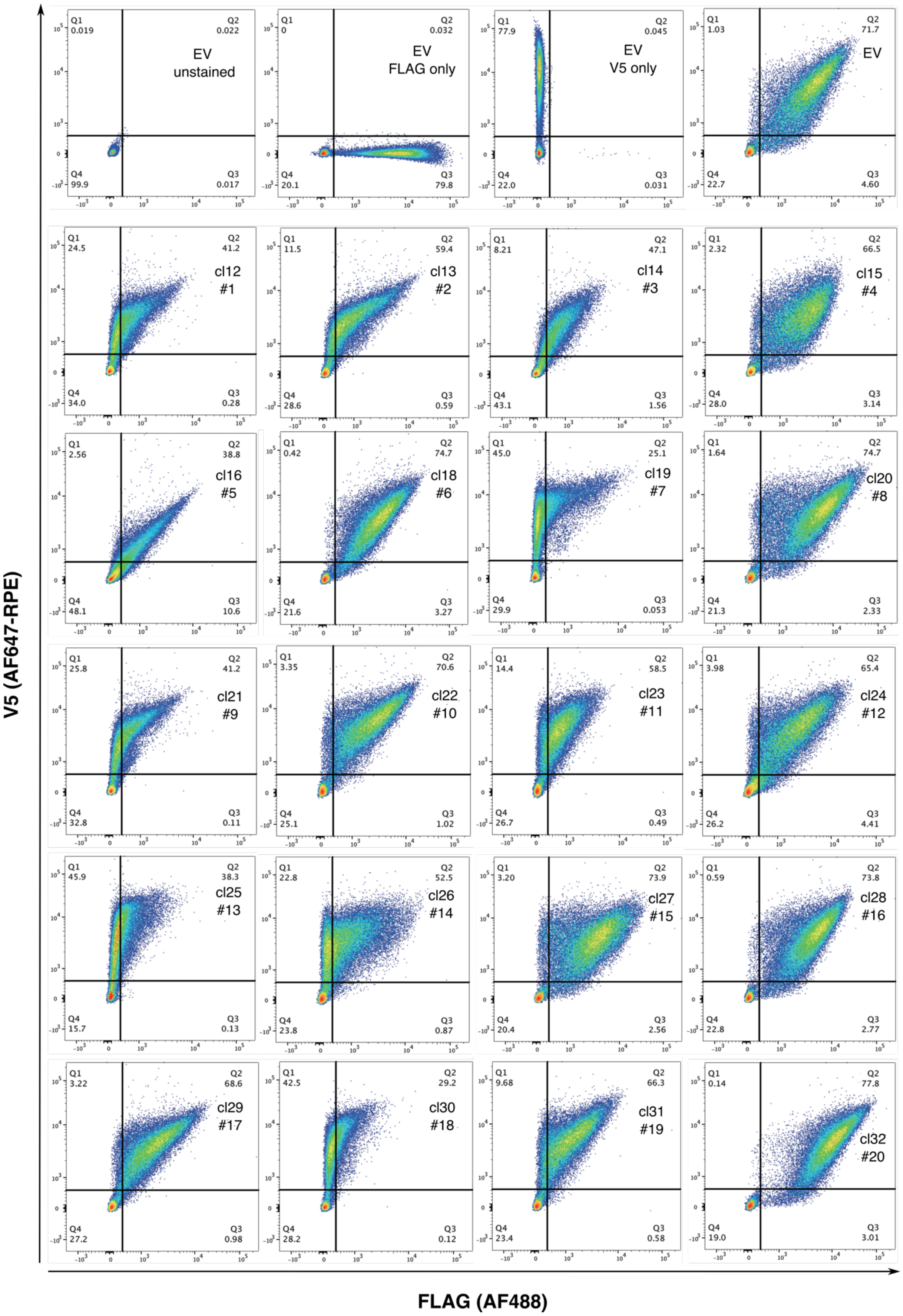
V5 (AF647-RPE)
FLAG (AF488)
EV unstained
Q1 0.019 Q2 0.022 Q3 0.017 Q4 99.9
EV FLAG only
Q1 0 Q2 0.032 Q3 79.8 Q4 20.1
EV V5 only
Q1 77.9 Q2 0.045 Q3 0.031 Q4 22.0
EV
Q1 1.03 Q2 71.7 Q3 4.60 Q4 22.7
cl12 #1
Q1 24.5 Q2 41.2 Q3 0.28 Q4 34.0
cl13 #2
Q1 11.5 Q2 59.4 Q3 0.59 Q4 28.6
cl14 #3
Q1 8.21 Q2 47.1 Q3 1.56 Q4 43.1
cl15 #4
Q1 2.32 Q2 66.5 Q3 3.14 Q4 28.0
cl16 #5
Q1 2.56 Q2 38.8 Q3 10.6 Q4 48.1
cl18 #6
Q1 0.42 Q2 74.7 Q3 3.27 Q4 21.6
cl19 #7
Q1 45.0 Q2 25.1 Q3 0.053 Q4 29.9
cl20 #8
Q1 1.64 Q2 74.7 Q3 2.33 Q4 21.3
cl21 #9
Q1 25.8 Q2 41.2 Q3 0.11 Q4 32.8
cl22 #10
Q1 3.35 Q2 70.6 Q3 1.02 Q4 25.1
cl23 #11
Q1 14.4 Q2 58.5 Q3 0.49 Q4 26.7
cl24 #12
Q1 3.98 Q2 65.4 Q3 4.41 Q4 26.2
cl25 #13
Q1 45.9 Q2 38.3 Q3 0.13 Q4 15.7
cl26 #14
Q1 22.8 Q2 52.5 Q3 0.87 Q4 23.8
cl27 #15
Q1 3.20 Q2 73.9 Q3 2.56 Q4 20.4
cl28 #16
Q1 0.59 Q2 73.8 Q3 2.77 Q4 22.8
cl29 #17
Q1 3.22 Q2 68.6 Q3 0.98 Q4 27.2
cl30 #18
Q1 42.5 Q2 29.2 Q3 0.12 Q4 28.2
cl31 #19
Q1 9.68 Q2 66.3 Q3 0.58 Q4 23.4
cl32 #20
Q1 0.14 Q2 77.8 Q3 3.01 Q4 19.0

**Supplementary Fig. 2. Flow cytometry plots of select randomly picked clones after two rounds of sorting**. Top row: FLAG-V5-Sag1 control strain (EV= empty vector) unstained, single stained, and double stained controls. Other rows are the double-stained single clones. Clone number (cl...) matches the clone identity in **Suppl. Table 4**, and the number below (#...) indicates the fragment as used for secretion check on western blot (**Fig. 1d** and **Suppl. Table 5**). Clone 17 was omitted as it was not detected in the sequencing data.

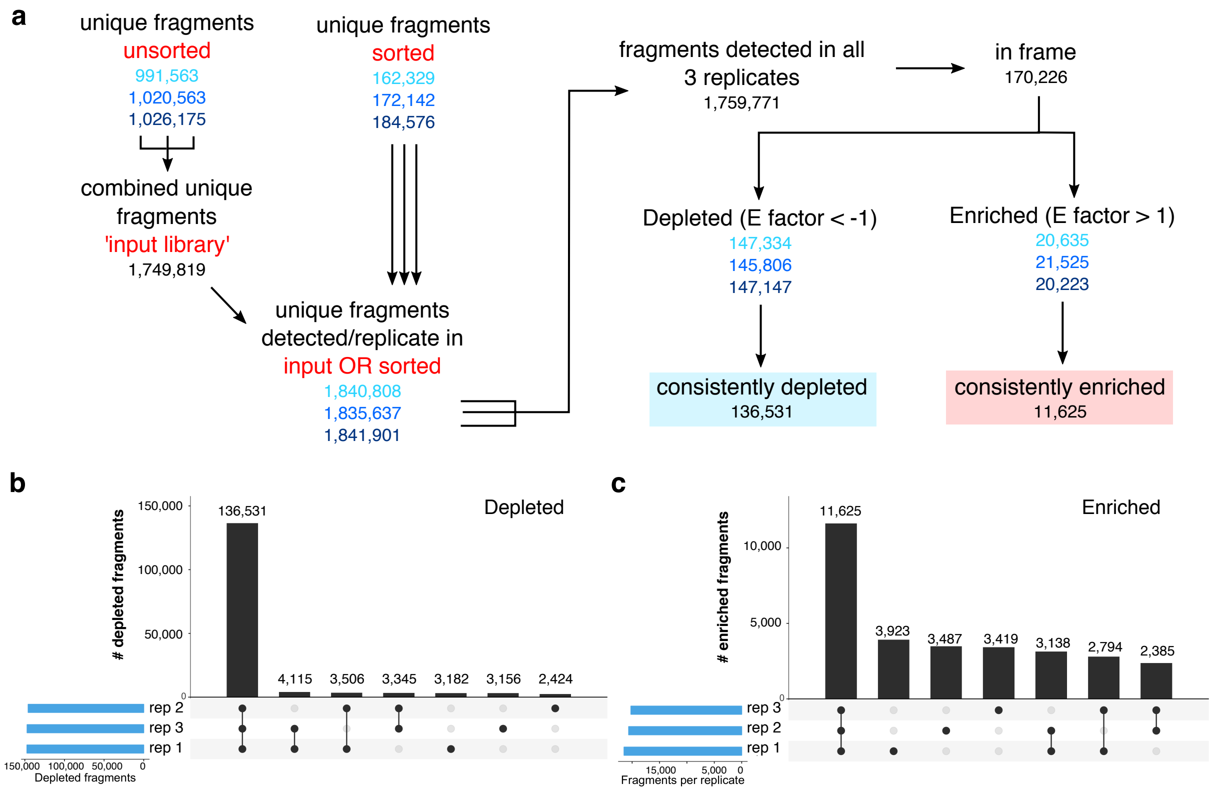


**Supplementary Fig. 3. Enriched/depleted fragment classification and replicate statistics in *S. cerevisiae* screens. a)** Fragment processing workflow for classification into enriched/depleted fragments. The number of fragments in each of the three replicate screens is shown in different shades of blue. A fragment is considered detected in a replicate when it is detected in either the sorted sample of that experiment, and/or in the input library. Three independent samples of this input library were sequenced, and we combined all of the fragments detected in the total sequence read pool to enhance the sequencing depth of this highly diverse unsorted input fragment library. Consistently depleted (or enriched) fragments are in-frame fragments, detected in all three experiments, for which the E factor is < -1 (or > 1) in all experiments. **b-c**) Replicate experiment overlap. Left (b): depleted fragments. Right (c): enriched fragments.

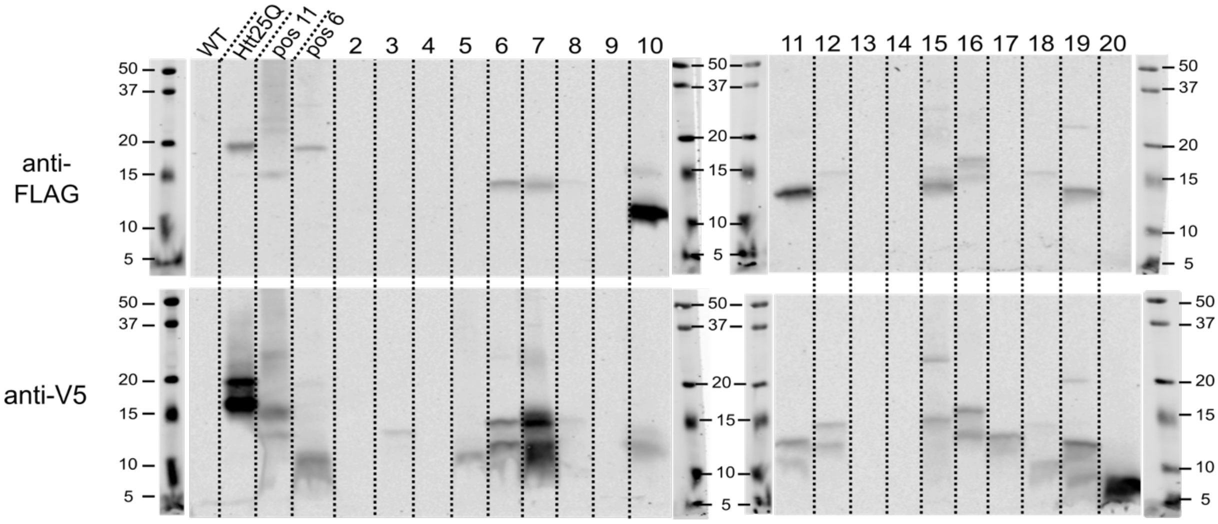


**Supplementary Fig. 4. A fraction of depleted fragments are still secretable.** Twenty random human fragments that were depleted after SECRiFY screening were expressed in *S. cerevisiae* with N-terminal FLAG and C-terminal V5 tag but without Sag1 anchor, and presence in medium was assessed by Western Blot. Although most fragments remained undetected, several fragments still show some level of secretability (ie bands close to the expected molecular weight detectable by FLAG and V5), suggesting a relatively low negative predictive value, as expected for a screen with focus on positive selection. DNA for fragment 1 could not be synthesized. WT: *S. cerevisiae* R1158 medium (neg. control), Htt25Q: medium from *S. cerevisiae* secreting human Htt25Q (pos. control), pos 11: enriched fragment 11 (pos. control), pos 6: enriched fragment 6 (pos. control).

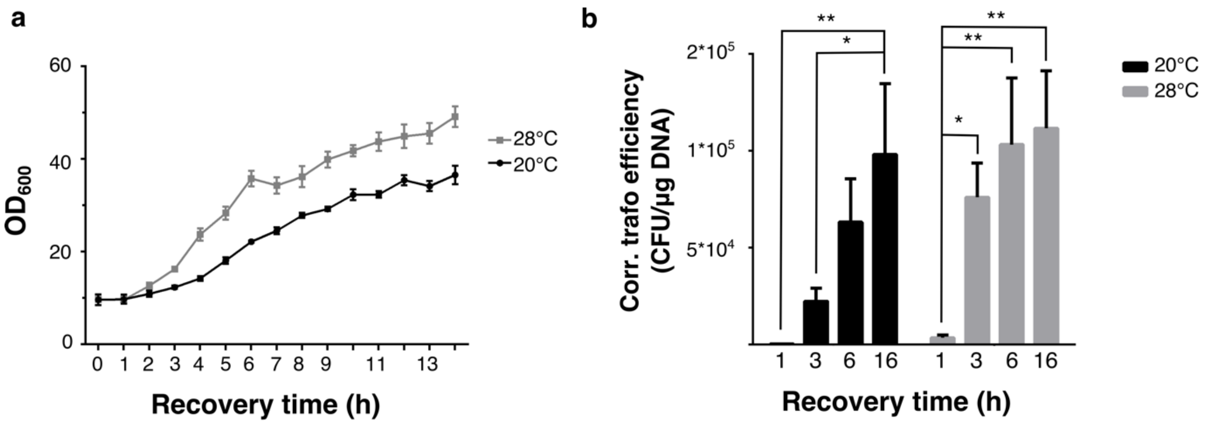


**Supplementary Fig. 5. Extending recovery time of transformed *P. pastoris* cells in YPD improves transformation efficiency. a**) Exponential growth of *P. pastoris* library transformants takes off after about 3h into recovery in YPD medium at 28°C after electroporation, suggesting it takes several hours for cells to recover from electroporation. Mean $OD_{600}$ ± SEM (n= 3). **b**) Transformation efficiency, corrected for growth occurring during recovery, significantly increases with the length of the recovery period at both 28°C and 20°C. Two-way repeated measures ANOVA with Tukey's post-hoc test for multiple comparisons, mean corrected transformation efficiency ± SEM (n=4). * $p<0.05$, ** $p<0.01$. Electroporation was done with 100 ng library DNA per reaction and selection on YPD pH 8.0 + 20 µg/ml zeocin agar plates.

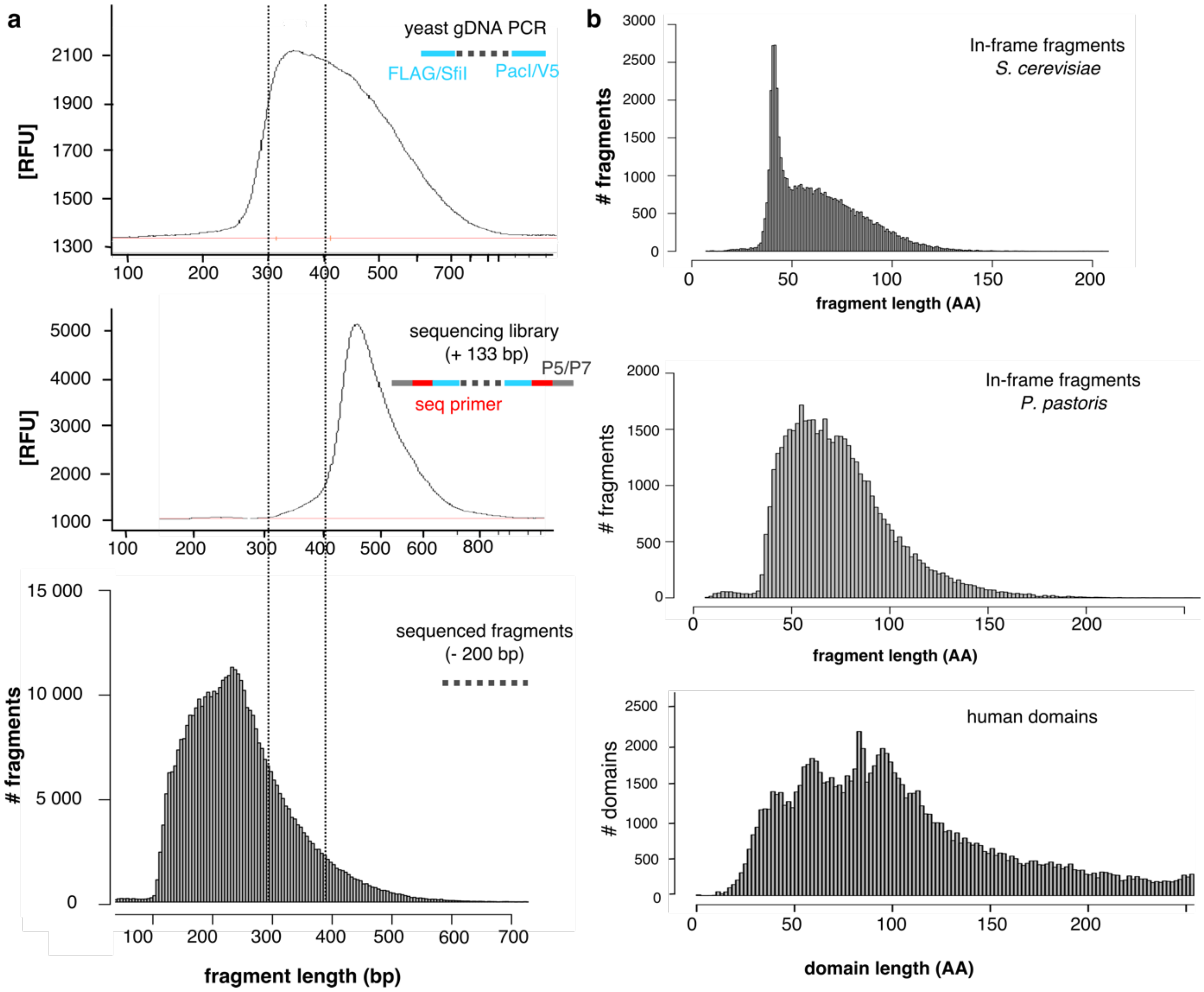


**Supplementary Fig. 6. Size distributions of library cDNA fragments. a)** Illumina sequencing of pools of sequences with a broad size distribution leads to a bias towards smaller fragments**.** Fragments from unsorted and sorted *P. pastoris* cells are picked up by PCR from plasmids integrated in gDNA. These fragments are between 300-500 bp in length (upper panel), including the additional 67 bp from the common priming sequences (blue). After sequencing library construction, adding the sequences necessary for paired-end sequencing (red) and flow-cell binding (gray) (+ 133 bp in total compared to the amplified fragments), total fragment length shifts accordingly, though towards a slightly narrower distribution partially depleted in the longer sequences (middle panel). The cDNA fragment distribution after sequencing and read processing (which removes 200 bp of method-related sequences) shows that most sequenced fragments are between 150-400 bp long (bottom panel, n= 423,692, *P. pastoris* experiment). Thus, we observe a bias towards smaller fragments (100-200 bp), especially during sequencing. This likely has to do with biases in bridge amplification inherent to Illumina sequencing. Such bias can likely be overcome in future experiments by using mate-pair sequencing protocols. This would also enable the analysis of libraries encoding much larger protein fragments or intact proteins **b**) Final sequenced in-frame fragment size distribution in our *S. cerevisiae* and *P. pastoris* experiments compared to that of human protein domains in Gene3D (v14.0.0, n= 104,734). The overrepresentation of fragments smaller than 50 AA in the *S. cerevisiae* experiments (upper panel) was resolved in the *P. pastoris* experiments (middle panel) by avoiding overamplification during sequencing library prep. Nevertheless, both fragment distributions largely overlap with the size distribution of human protein domains.

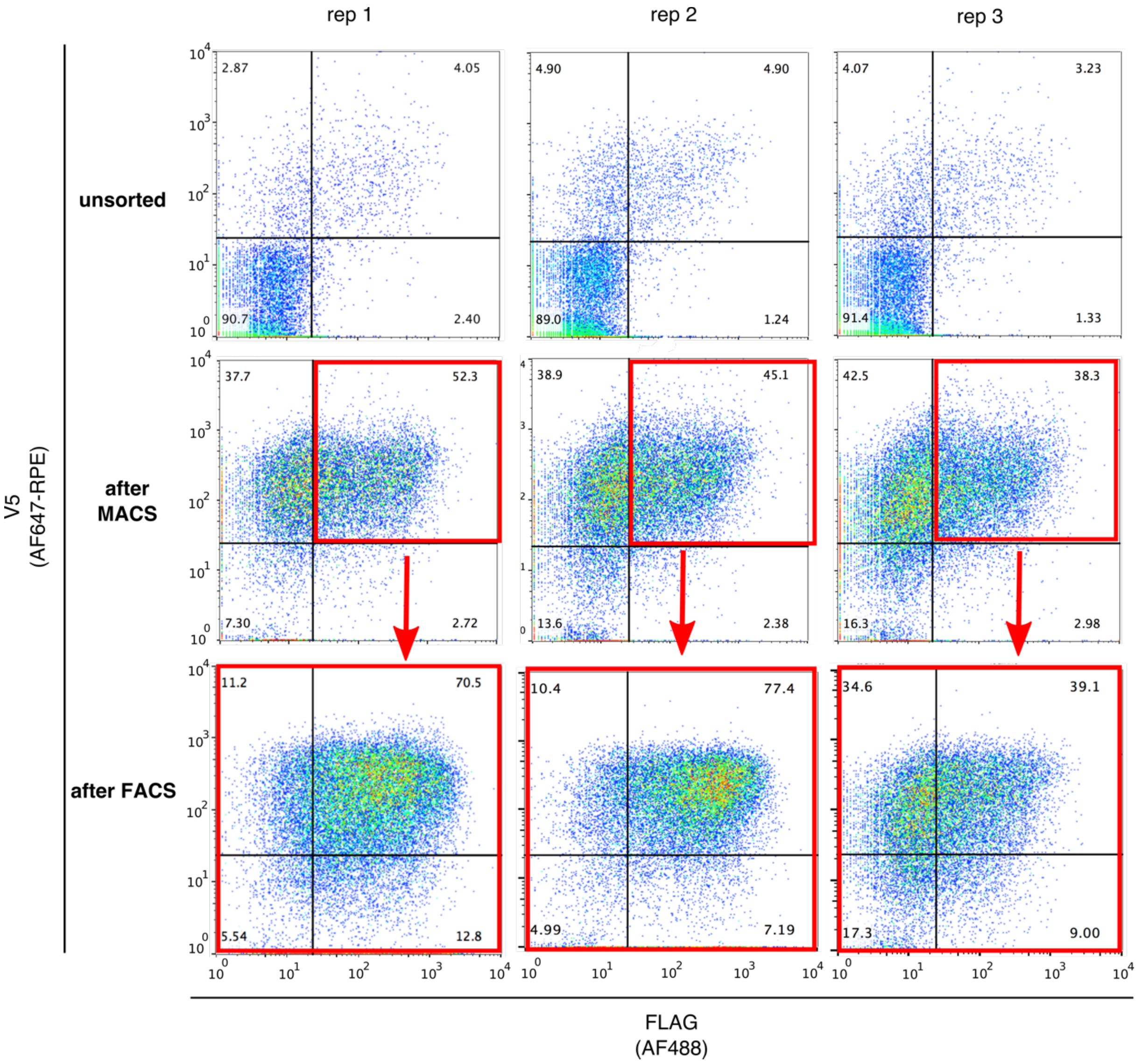


**Supplementary Fig. 7. Sorting of fragment-displaying *P. pastoris* library clones.** Expression of the human fragment library in *P. pastoris* was induced (upper row panels) and the pool was first enriched for $V5^+$ cells, displaying a fragment with an intact C-terminus, using MACS (middle row panels). This enriched subset was immediately sorted for $FLAG^+V5^+$ cells using FACS, representing display of fragments with both termini intact. Sorted cells were allowed to recover and were frozen for storage. The purity of these sorted cells was checked by regrowing the cells, inducing fragment expression, cell staining, and flow cytometry (lower row panels, ‘after FACS’). The sorting of the library was replicated independently three times.

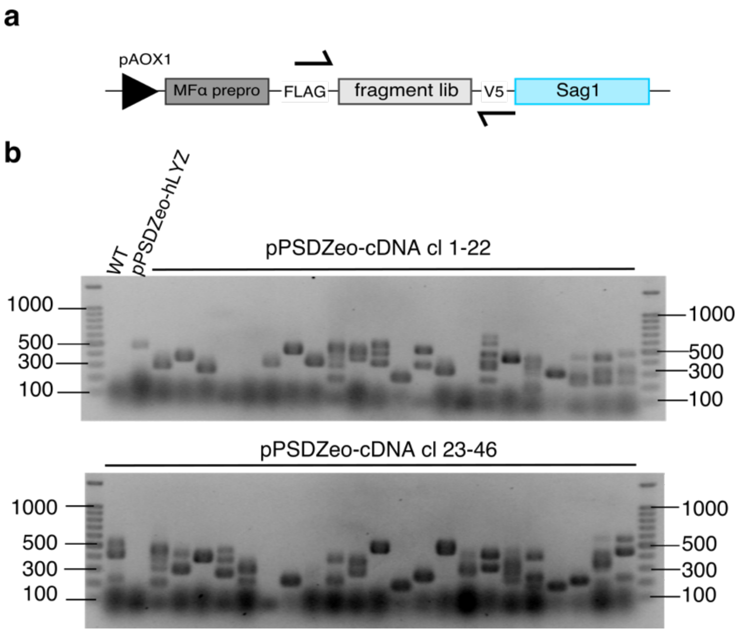


**Supplementary Fig. 8. Co-integration of multiple fragments in the *P. pastoris* genome as shown by colony PCR on randomly picked library clones. a)** The pPSDZeo surface display cassette in the *P. pastoris* yeast library clones. Primers for the colony PCR to amplify fragment inserts are represented by black arrows. **b)** Colony PCR of single library clones. WT= *P. pastoris* untransformed strain GS115. A strain displaying human lysozyme, marked as pPSDZeo-hLYZ, was used as a positive control and generates a band of 447 bp. Band sizes reflect the size of the fragment insert + 40 bp primer sequence. 41/46 clones contain at least one insert, 22 of those contain more than one insert. The average clone has 3 inserts. This accounts for the higher % of clones (+/- 4%, see **Supplementary Fig. 7**) that express a detectable protein fragment in the unsorted library than is the case for *S. cerevisiae* (+/- 1.7%). However, this co-integration of several fragment expression constructs will not significantly affect the reliability of determining which of these fragments are secretable: as only a tiny fraction (about 1.7%) of inserts yield a detectably surface-displayed protein fragment, the chance of co-occurrence of two such surface-displayable fragments is very low (about 1.7% of the sorted cells). This issue is further mitigated because we replicate the sorting experiment 3 times and only consider those fragments that are observed to be enriched consistently across the 3 experiments.

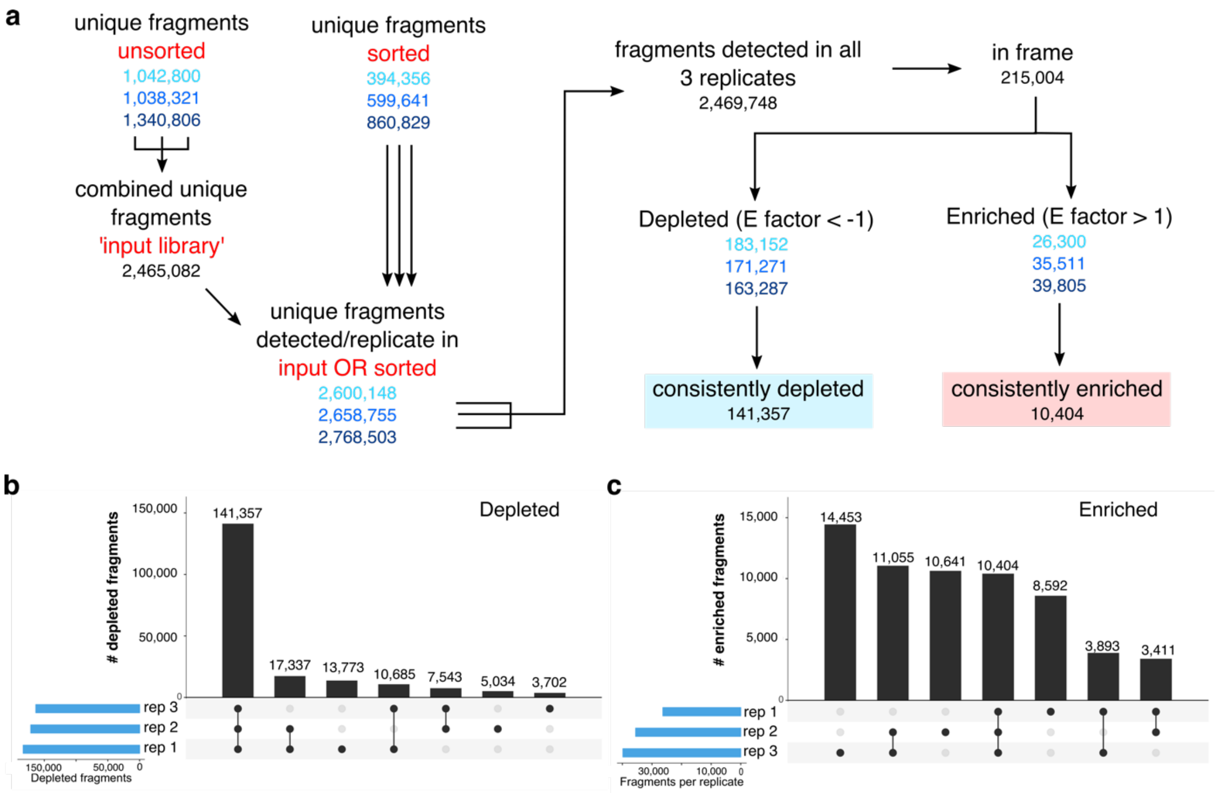


**Supplementary Fig. 9. Enriched/depleted fragment classification and replicate statistics in *P. pastoris* screens.** Fragment processing workflow for classification into enriched/depleted fragments, for our *P. pastoris* screens. As in Supplementary Fig. 3, the number of fragments in each of the three replicate screens is shown in different shades of blue. A fragment is considered detected in a replicate when it is detected in either the sorted sample of that experiment, and/or in the input library, which represents the total diversity of the starting yeast library before sorting to offset fragments missed due to insufficient depth of sequencing of unsorted samples. Consistently depleted (or enriched) fragments are in-frame fragments, detected in all three experiments, for which the E factor is < -1 (or > 1) in all experiments. **b-c**) Replicate experiment overlap. Left (b): depleted fragments. Right (c): enriched fragments.

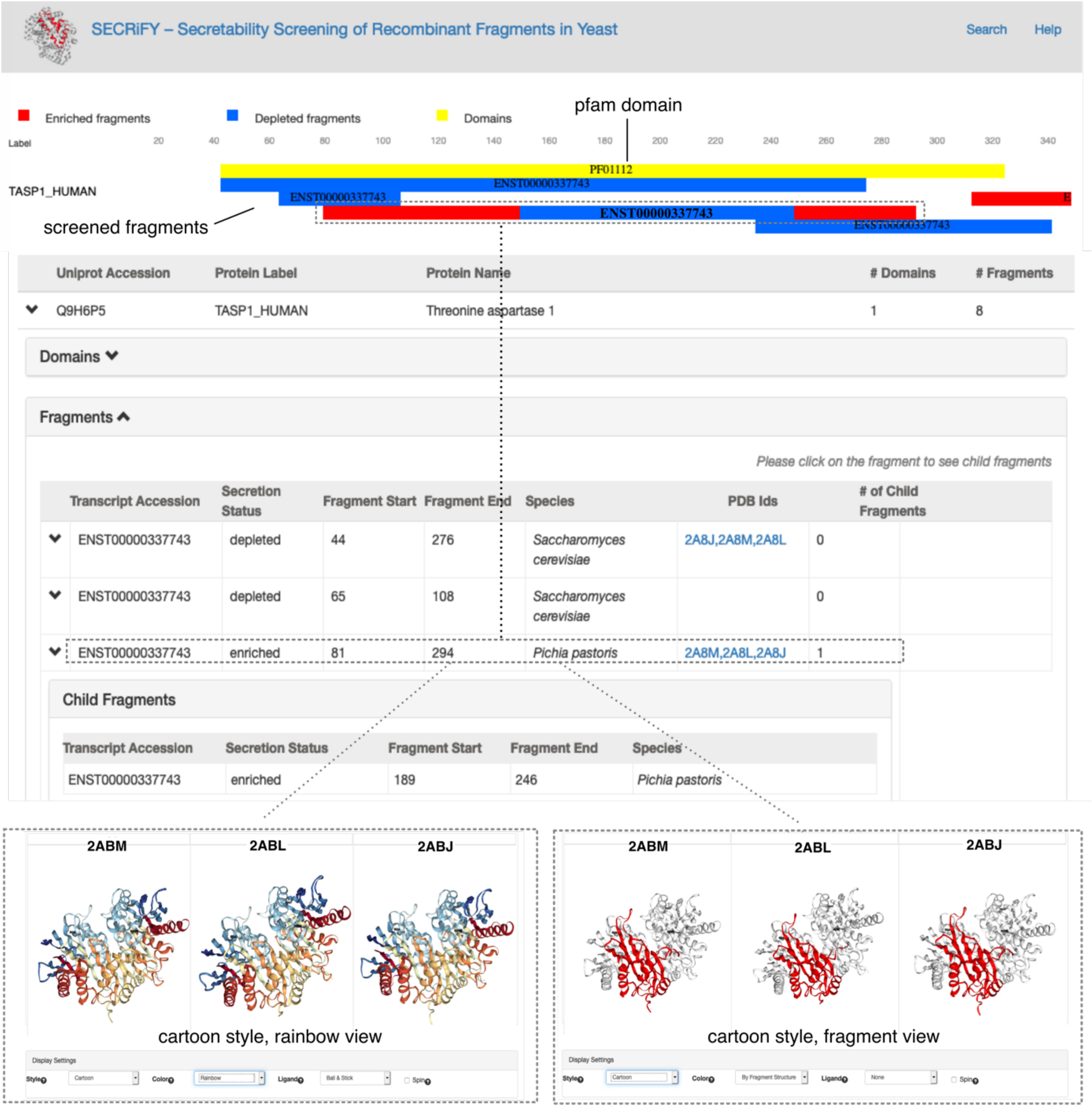


**Supplementary Fig. 10. Browsing secretability data using the SECRiFY website.** Users can survey which secretable or depleted fragments map to their protein of interest (here TASP1) or pfam domain of interest. The protein sequence and the SECRiFY-interrogated sequence fragments (blue for depleted, red for enriched) can be viewed using the sliding panel at the top of the page. Each fragment is labeled according to the Ensembl ID of the human transcript it mapped to. Yellow bars indicate known pfam domain regions. Pull-down panels give an overview of representative fragments (the longest fragment of a cluster of fragments with 100% overlap) and the corresponding child fragment(s) of the cluster, as well as their secretability status and the organism in which the screen was performed. Clicking on the PDB ID's to which the representative fragment mapped to enables a structural examination of the corresponding PDB files and mapped fragment in a variety of display styles (bottom insets left and right).

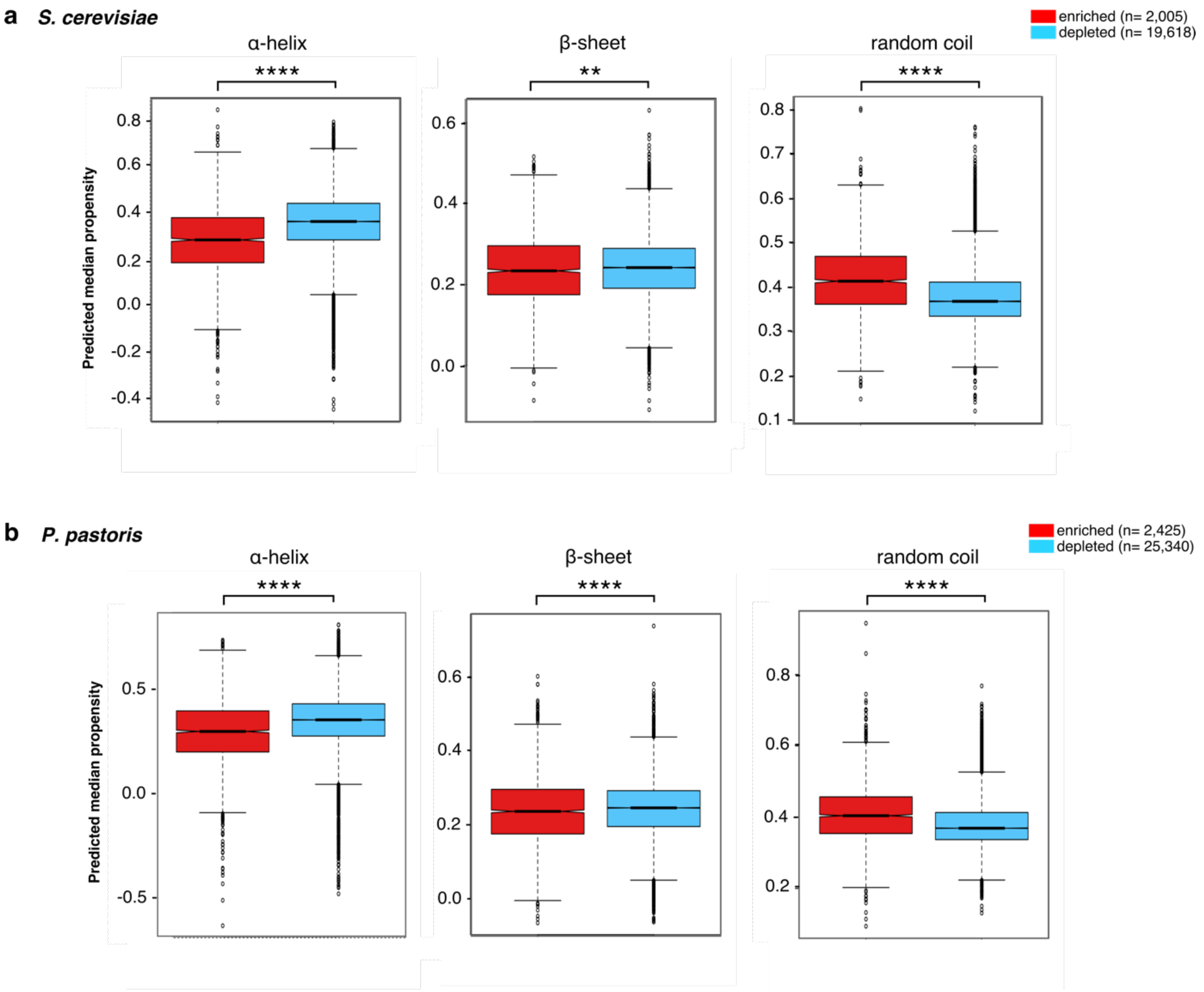


**Supplementary Fig. 11. Predicted helical, sheet and random coil propensity of subsets of consolidated enriched (secretable) and depleted fragments.** Box plots indicate the distribution of the median helix, sheet or coil propensity of amino acid residues, summarized per consolidated fragment. Whiskers reflect the maximum value or the respective quartile value times 1.5 the interquartile range, whichever is less. The notch displays a confidence interval based on the median plus/minus 1.57 times the interquartile range divided by the square root of the number of points. If the notches of two boxes do not overlap, this is strong evidence that their medians differ significantly. In both organisms, secretable fragments tend to have less α-helical tendencies (*S. cerevisiae:* p= $2.946*10^{-124}$*, P. pastoris:* p= $2.371*10^{-76}$*,* Mann-Whitney-Wilcoxon test), similar β-sheet propensities (*S. cerevisiae*: p= $1.262*10^{-03}$, *P. pastoris*: p= $3.360*10^{-05}$, Mann-Whitney-Wilcoxon test), and higher random coil frequencies that depleted fragments (*S. cerevisiae*: p= $1.993*10^{-127}$, *P. pastoris*: p= $1.098*10^{-85}$, Mann-Whitney-Wilcoxon test). Panel **a** was reproduced from Fig. 3 in the main text to compare with the fragments expressed in *P. pastoris* in **b**.

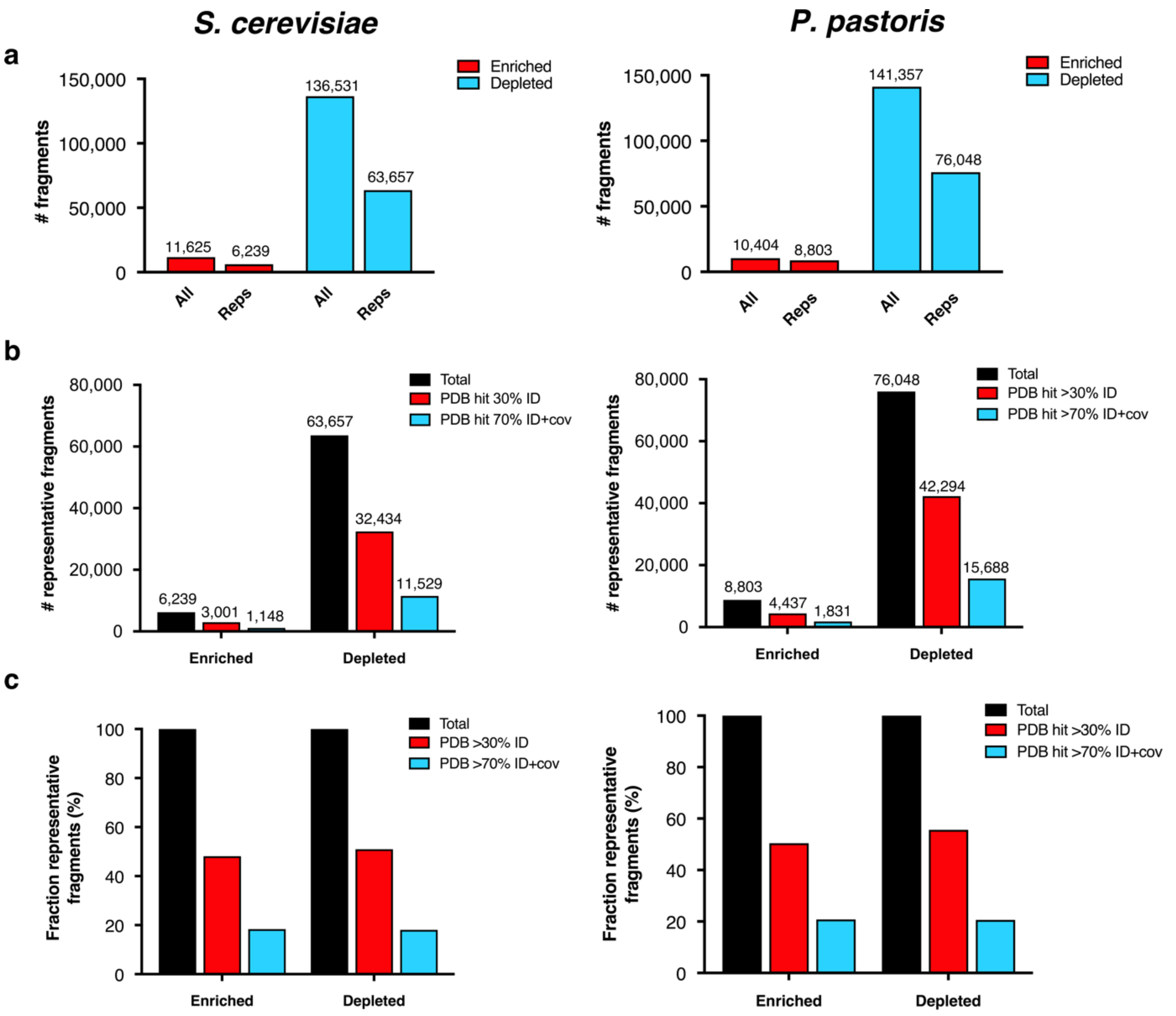


**Supplementary Fig. 12. Mapping of representative fragments to PDB. a)** To reduce the redundancy in the data before mapping to PDB, fragments that are 100% overlapping with longer fragments were clustered to a single representative fragment (i.e. the longest fragment). Graphs show the absolute number of fragments before and after clustering in *S. cerevisiae* (left) and *P. pastoris* (right) datasets. Depleted sets have a higher fraction of redundant fragments and were reduced to about 50% after clustering. Reps= representative fragments. **b**) Roughly 50% of the representative fragments match a sequence with a structure in PDB with ≥ 30% sequence identity, and roughly a third of those with ≥ 70% sequence identity and ≥ 70% query coverage. Data for *S. cerevisiae* (left) and *P. pastoris* (right) fragments. **c**) Same as in b but with fractional instead of absolute numbers.

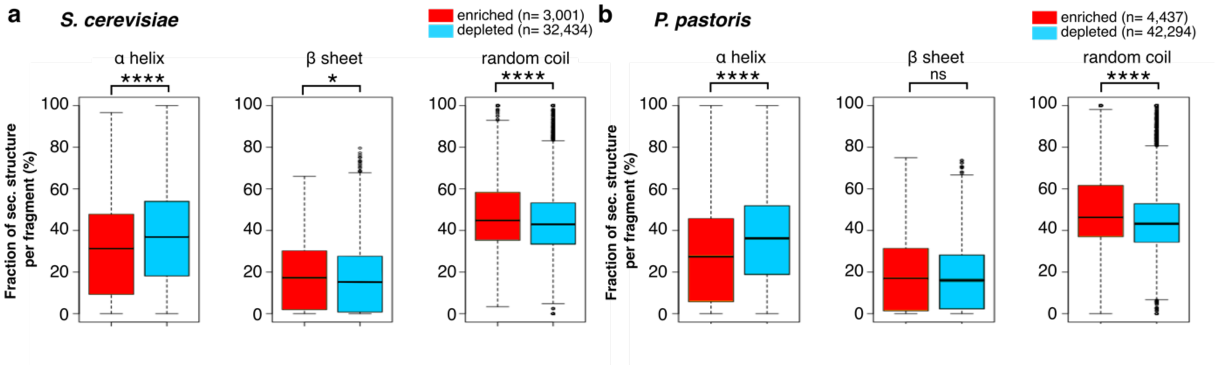


**Supplementary Fig. 13. Enriched fragments on average have a lower average α-helical but higher coiled coil content.** Percentage of residues in secondary structure elements in representative enriched and depleted fragments that mapped to PDB structures in both screens. Depleted fragments tend to have a higher helical propensity (*S. cerevisiae:* p= $2.35*10^{-8}$*, P. pastoris:* p= $3.12*10^{-22}$*,* Mann-Whitney-Wilcoxon test), whereas enriched fragments tend to have a higher coil tendency (*S. cerevisiae:* p= $1.24*10^{-4}$, *P. pastoris:* p= $2.55*10^{-14}$, Mann-Whitney-Wilcoxon test). Differences in β-sheet propensity are less pronounced (*S. cerevisiae*: p= 0.02, *P. pastoris:* p= 0.15*,* Mann-Whitney-Wilcoxon test). Panel **a** was reproduced from Fig. 3 in the main text to compare with the fragments expressed in *P. pastoris* in **b**.

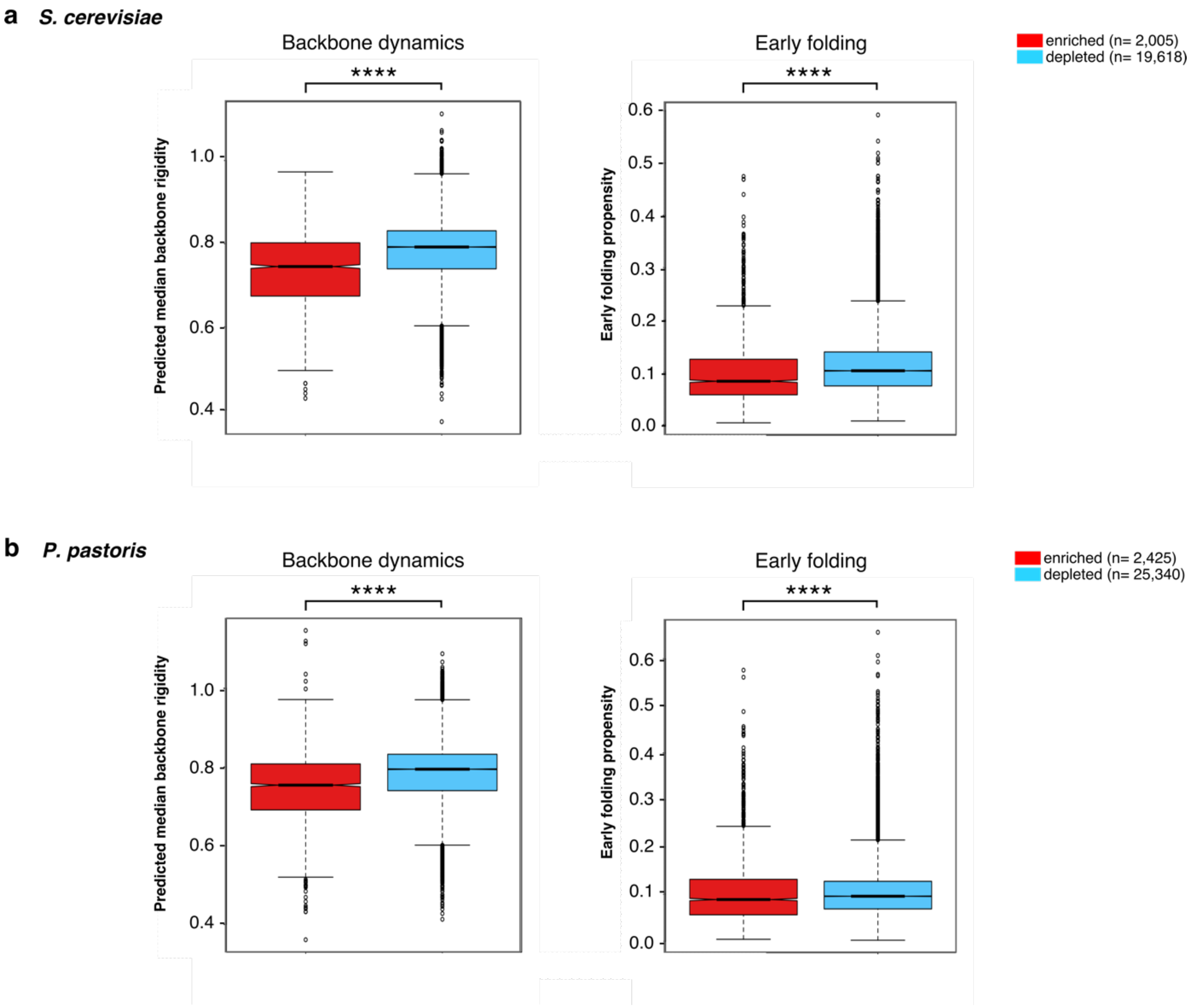


**Supplementary Fig. 14. Backbone dynamics and early fold predictions.** Notched box plots indicate median predicted backbone rigidity or early fold propensity of each consolidated fragment. In terms of backbone dynamics, enriched fragments score lower on the rigidity scale and seem thus to be more flexible than depleted fragments (*S. cerevisiae*: p= $5.076*10^{-118}$, *P. pastoris*: p= $2.498*10^{-93}$, Mann-Whitney-Wilcoxon test). The median propensity for early folding is much more similar between enriched and depleted fragments, despite significant differences in the propensity distribution (*S. cerevisiae*: p= $2.283*10^{-52}$, *P. pastoris*: p= $1.597*10^{-10}$, Mann-Whitney-Wilcoxon test). Panel **a** (backbone dynamics) was reproduced from Fig. 3 in the main text to compare with the fragments expressed in *P. pastoris* in **b**.

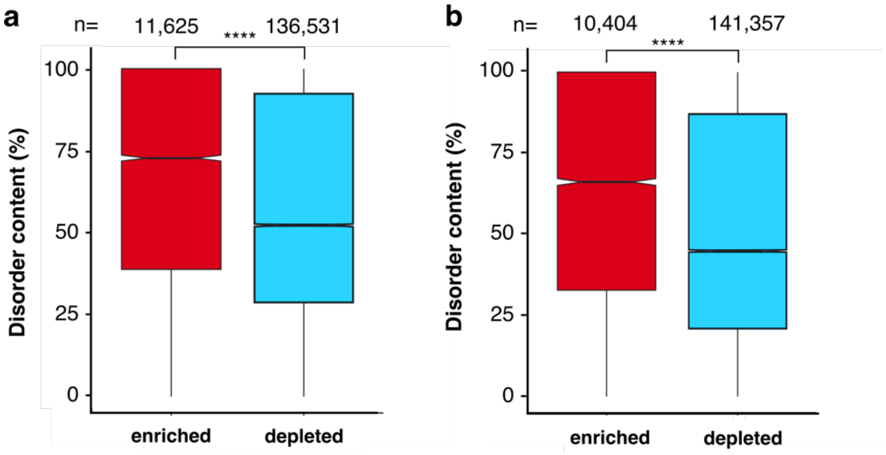


**Supplementary Fig. 15. RAPID protein disorder content predictions in *S. cerevisiae* (a) and *P. pastoris* (b).** Secretable fragments are predicted to have a higher average disorder content compared to depleted fragments in both organisms. Panel **a** is the same as in Fig. 3 but was reproduced here along the boxplot for the *P. pastoris* screen for comparison. Mann-Whitney-Wilcoxon test, **** $p < 2.2*10^{-16}$.

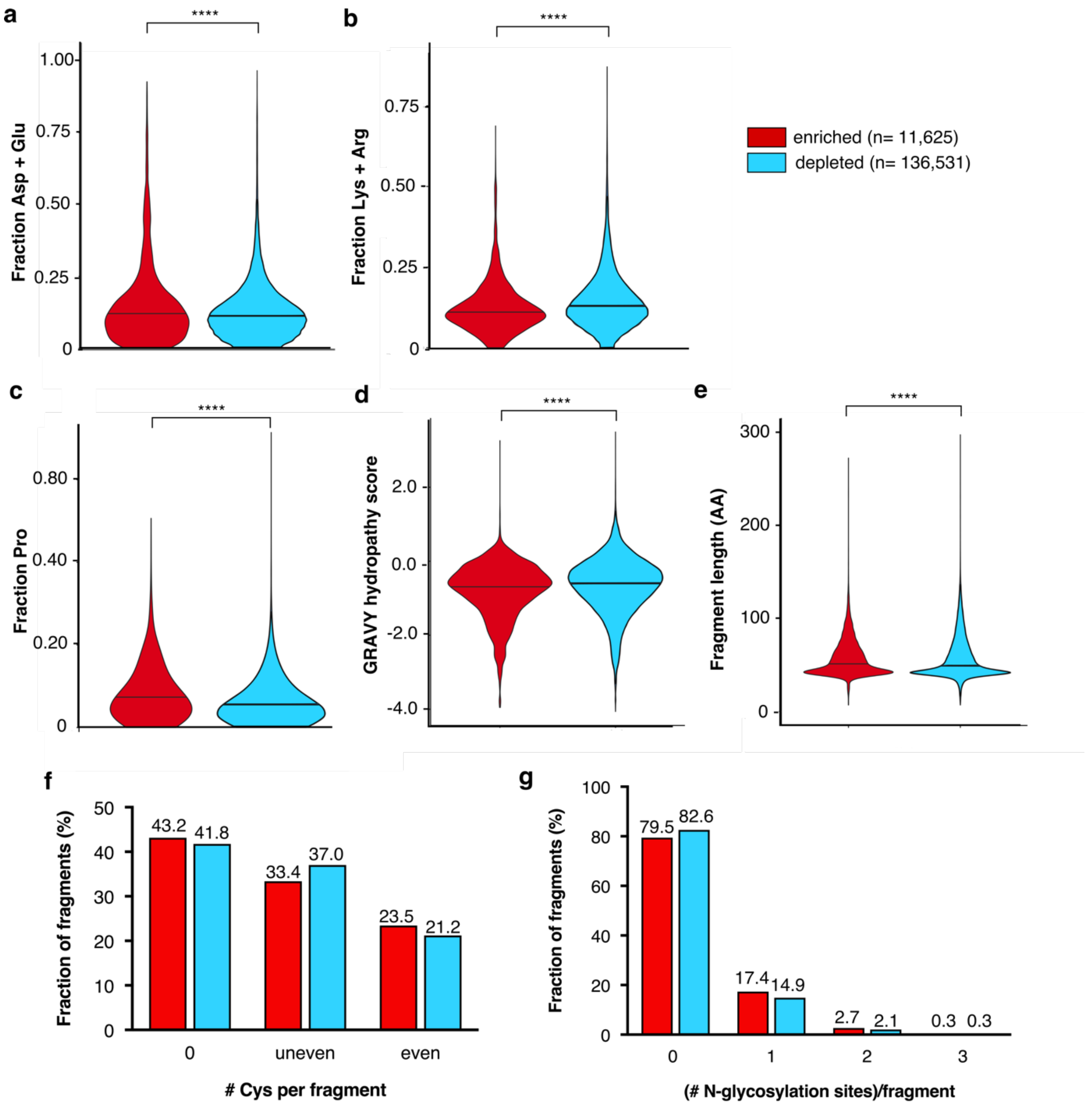


**Supplementary Fig. 16. Select patterns in human fragments displayed by *S. cerevisiae*. a**) Enriched fragments have a slightly larger proportion of fragments with a higher fraction of negatively charged amino acids Asp and Glu per fragment. **b**) The fraction of positively charged amino acids Lys and Arg is slightly lower in enriched fragments, which could be related to ribosome stalling on polybasic residues. **c**) The fraction of Pro is higher in enriched fragments. d) Enriched fragments tend to be less hydrophobic, as indicated with a slightly lower overall GRAVY hydropathy score. **e**) Length distributions in enriched and depleted fragments. In all violin plots (**a-e**), the black line indicates the median. Distributions were compared using the Mann-Whitney-Wilcoxon test. **** $p<2.2*10^{-16}$. **f**) Considering the oxidative environment in the ER, fragments that require paired Cys for folding could have been more prominent in enriched (secretable) fragments, but differences are only minor. **g**) Considering the important role of N-glycosylation for the folding of many secretory proteins, the number of potential N-glycosylation sites (Asn-X-Ser/Thr with X not Pro) was examined in enriched and depleted fragments, but no large differences are observed.

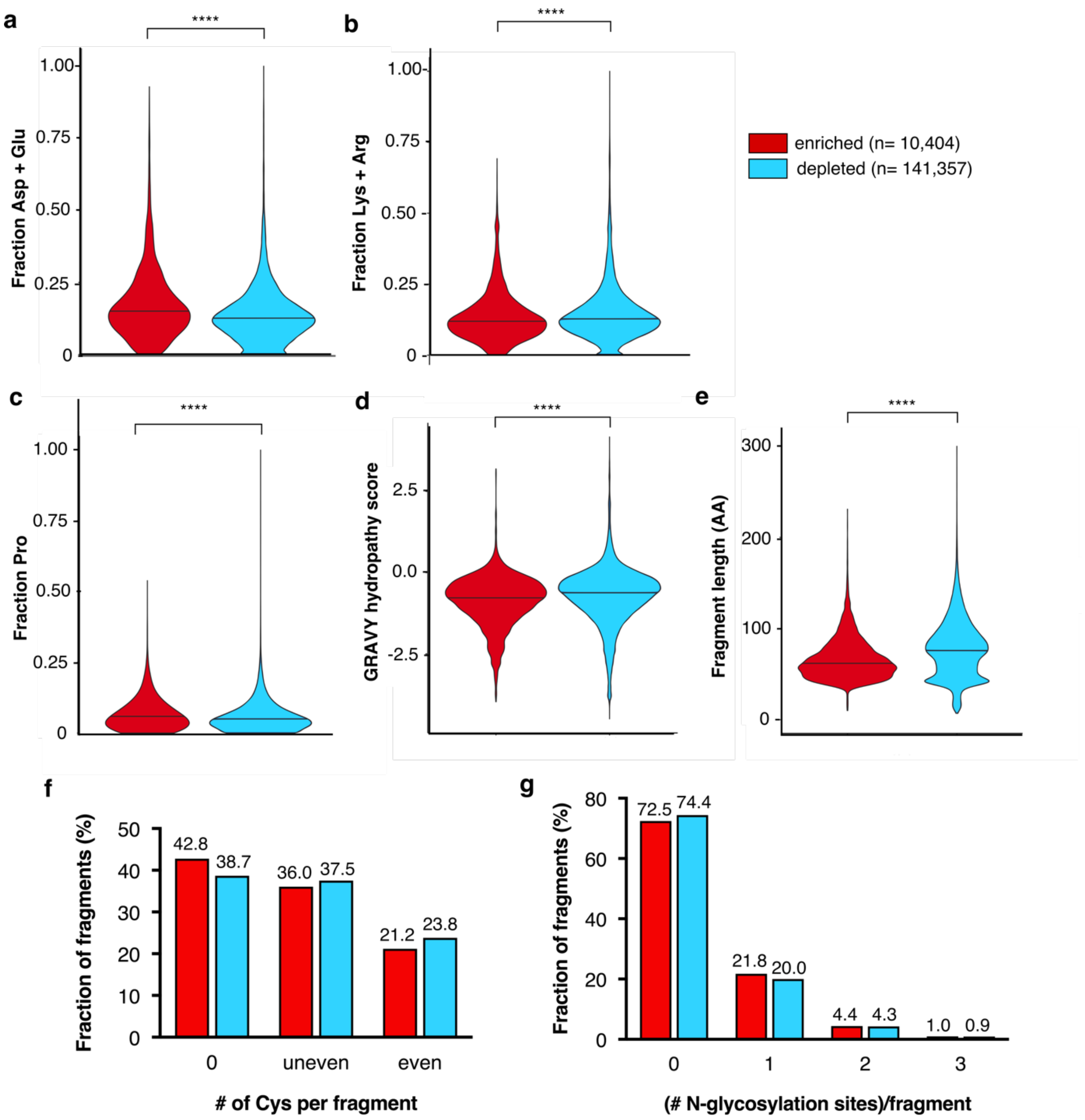


**Supplementary Fig. 17. Select patterns in human fragments displayed by *P. pastoris*.** We observed similar patterns in our *S. cerevisiae* screens (**Supplementary Fig. 15**): enriched fragments tend to have a slighty larger fraction of negatively charged amino acids (**a**) and Pro (**c**) (also residues that are often enriched in disordered proteins), less positively charged amino acids (**b**), less hydrophobic overall (**d**), and smaller in length (**e**). There is no clear difference in the number of Cys pairs (**f**) nor potential N-glycosylation sites (**g**). Black lines in violin plots indicate the median. Distributions were compared using the Mann-Whitney-Wilcoxon test. **** $p<2.2*10^{-16}$.

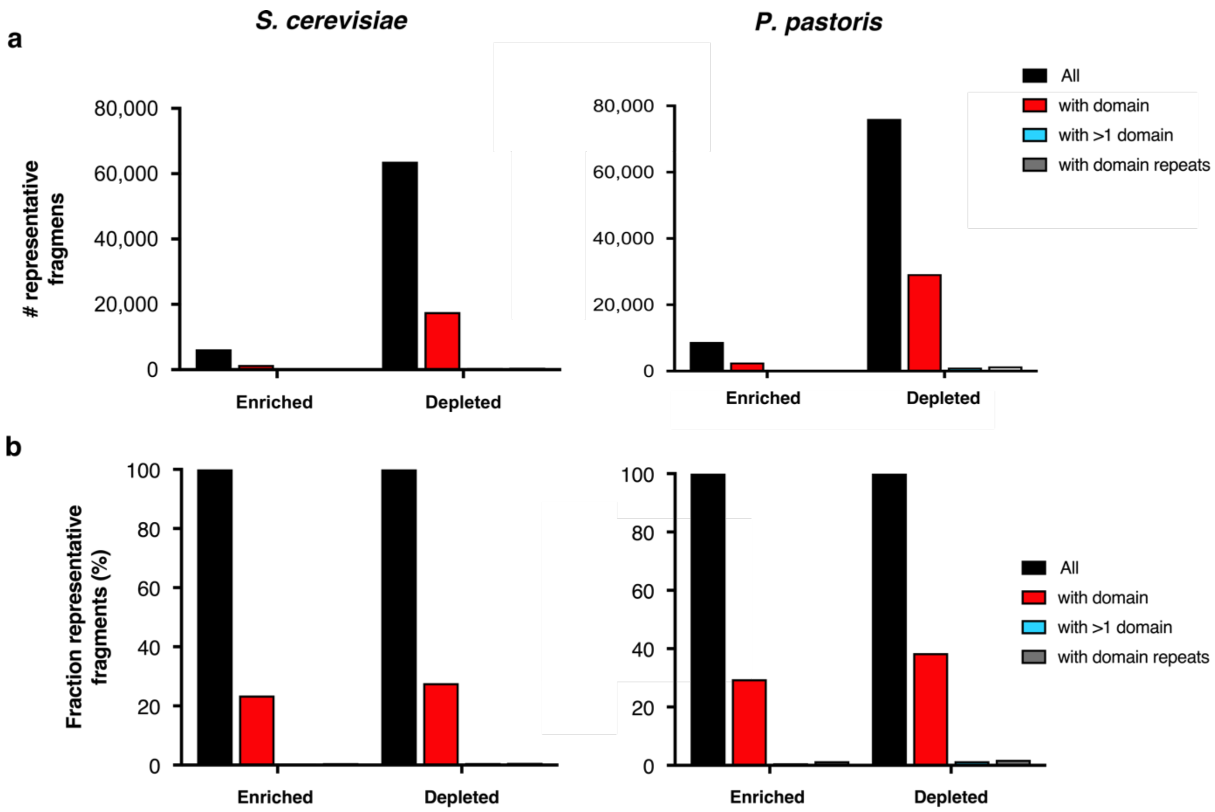


**Supplementary Fig. 18. Pfam domains in enriched and depleted fragments**. Number (a) and fraction (b) of representative fragments with one or more hits with Pfam domains. Left: *S. cerevisiae* screens, right: *P. pastoris* screens.

### Supplementary tables

| sample | | # reads | Fraction of reads (%) |
|---|---|---|---|
| **total** | | 214,773,952 | 100.00 |
| **passing filter (PF)** | | 199,092,922 | **92.70** |
| **PF + barcode identified** | unsorted 1 | 41,979,118 | 21.09 |
| | unsorted 2 | 41,527,810 | 20.86 |
| | unsorted 3 | 46,443,265 | 23.33 |
| | sorted 1 | 19,426,627 | 9.76 |
| | sorted 2 | 18,508,105 | 9.30 |
| | sorted 3 | 19,448,837 | 9.77 |
| | **total** | 187,333,764 | **94.09** |

| sample | fraction of reads remaining after (%) | |
|---|---|---|
| | **Illumina sequence removal** | **FLAG/V5 trimming** |
| unsorted 1 | 95.09 | 95.3 |
| unsorted 2 | 94.26 | 94.9 |
| unsorted 3 | 89.39 | 93.8 |
| sorted 1 | 92.04 | 94.7 |
| sorted 2 | 90.43 | 94 |
| sorted 3 | 89.82 | 94.2 |
| **total** | **91.84** | **94.48** |

**Supplementary Table 1.** ***S. cerevisiae*** **secretability screening sequencing and trim statistics.** Left: number of reads (twice the number of pairs) obtained after Illumina sequencing of 3 replicate samples. Data was summarized over four sequencing chip lanes. Of the 200 million reads passing the filter (PF), about 95% were barcode identified. Right: read statistics after trimming. In a first round, all reads matching Illumina flow cell or primer binding sequences were removed (on average about 9% of reads). Next, all FLAG+SfiI and V5+PacI sequences were trimmed off, and untrimmed or very short (< 20bp) reads were discarded, leaving roughly 95% of those sequences for mapping.

| | | unsorted 1 | unsorted 2 | unsorted 3 | sorted 1 | sorted 2 | sorted 3 |
|---|---|---|---|---|---|---|---|
| | input total reads | 38,051,552 | 37,155,742 | 38,926,446 | 16,940,044 | 15,729,448 | 16,455,338 |
| **R 1** | # reads | 19,025,776 | 18,577,871 | 19,463,223 | 8,470,022 | 7,864,724 | 8,227,669 |
| | mapped (%) | 86.43 | 86.46 | 86.54 | 90.62 | 91.29 | 88.61 |
| | unambiguous (%) | 18.87 | 18.45 | 18.25 | 15.25 | 14.94 | 14.33 |
| | ambiguous (%) | 67.56 | 68.02 | 68.30 | 75.37 | 76.35 | 74.29 |
| **R 2** | # reads | 19,025,776 | 18,577,871 | 19,463,223 | 8,470,022 | 7,864,724 | 8,227,669 |
| | mapped (%) | 86.53 | 86.29 | 86.37 | 90.50 | 91.19 | 88.50 |
| | unambiguous (%) | 18.83 | 18.41 | 18.22 | 15.25 | 14.97 | 14.33 |
| | ambiguous (%) | 67.43 | 67.88 | 68.15 | 75.25 | 76.23 | 74.16 |
| **total** | mated pairs (%) | 84.48 | 84.51 | 84.61 | 88.30 | 88.97 | 86.43 |
| | bad pairs (%) | 0.86 | 0.83 | 0.84 | 1.45 | 1.36 | 1.33 |
| | properly paired (%) | 84.48 | 84.50 | 84.60 | 88.30 | 88.97 | 88.56 |
| | singletons (%) | 1.05 | 1.04 | 1.01 | 0.81 | 0.92 | 0.80 |

**Supplementary Table 2. *S. cerevisiae* secretability screen mapping statistics**. Reads passing the filter (PF) and barcode identified reads were mapped to the hg38 transcriptome of known protein-coding genes using BBMap. The many ambiguous read mappings (due to the presence of multiple transcripts per gene) were assigned to the single best mapped transcript for read counting. This strategy is favored over mapping to a transcriptome of only canonical transcripts as it improves the fraction of mapped reads. Downstream fragment counting was further performed using properly-paired reads only. R 1: read 1, R 2: read 2.

| sample | # proper read pairs | # fragments |
|---|---|---|
| unsorted 1 | 16,072,901 | 991,563 |
| unsorted 2 | 15,699,215 | 1,020,569 |
| unsorted 3 | 16,466,734 | 1,026,175 |
| sorted 1 | 7,479,200 | 162,329 |
| sorted 2 | 6,996,856 | 172,142 |
| sorted 3 | 7,110,914 | 184,576 |
| **total** | 69,825,820 | // |

| sample | Average coverage per covered base | Bases covered (%) | Combined bases covered (%) |
|---|---|---|---|
| unsorted 1 | 155.18 | 27.40 | 27.55 |
| sorted 1 | 356.67 | 3.74 | |
| unsorted 2 | 155.82 | 26.18 | 26.34 |
| sorted 2 | 272.65 | 4.39 | |
| unsorted 3 | 167.5 | 25.29 | 25.46 |
| sorted 3 | 259.21 | 4.53 | |

**Supplementary Table 3.** ***S. cerevisiae*** **secretability screening fragment counts and transcriptome coverage statistics.** Left: number of properly mapped pairs and the corresponding number of different cDNA fragments obtained per sample. Right: calculated canonical transcriptome coverage. The fraction of canonical transcriptome bases covered (‘Based covered’) concerns only those bases covered with minimum 3 properly paired reads. The combined fraction reflects the fraction of the canonical transcriptome covered per sample. Note that these are estimates, as about 5% less reads can be mapped to the canonical transcriptome, and for fragments longer than 250 bp, the gap between the paired ends is not sequenced.

| Sample | median FLAG | rCV FLAG | median V5 | rCV V5 | unsorted count (FPTM) | sorted count (FPTM) | log2 FC |
|---|---|---|---|---|---|---|---|
| pSSD-R2 clone 1 | 1320.00 | 88.10 | 3353.00 | 104.00 | 60.97 | 1212.70 | 4.31 |
| pSSD-R2 clone 2 | 4717.00 | 92.60 | 3586.00 | 79.90 | 0.62 | 30.75 | 5.63 |
| pSSD-R2 clone 3 | 3586.00 | 83.30 | 3765.00 | 94.50 | 436.14 | 8614.56 | 4.30 |
| pSSD-R2 clone 4 | 7821.00 | 82.30 | 3521.00 | 96.00 | 102.66 | 2432.08 | 4.57 |
| pSSD-R2 clone 5 | 2951.00 | 91.10 | 4014.00 | 91.60 | 13.69 | 324.90 | 4.57 |
| pSSD-R2 clone 6 | 1246.00 | 111.00 | 3663.00 | 102.00 | 11.82 | 157.77 | 3.74 |
| pSSD-R2 clone 7 | 5184.00 | 90.70 | 2183.00 | 101.00 | 71.55 | 1262.17 | 4.14 |
| pSSD-R2 clone 8 | 2573.00 | 90.70 | 3881.00 | 97.60 | 0.00 | 8.02 | NA |
| pSSD-R2 clone 9 | 1619.00 | 92.10 | 2915.00 | 99.40 | 12.44 | 104.29 | 3.07 |
| pSSD-R2 clone 10 | 385.00 | 92.60 | 3070.00 | 85.70 | 0.00 | 239.33 | NA |
| pSSD-R2 clone 11 | 5594.00 | 82.00 | 3070.00 | 90.10 | 4.98 | 1218.04 | 7.93 |
| pSSD-R2 clone 12 | 444.00 | 95.60 | 1938.00 | 83.20 | 0.00 | 1.34 | NA |
| pSSD-R2 clone 13 | 796.00 | 114.00 | 2435.00 | 81.80 | 4.36 | 98.94 | 4.51 |
| pSSD-R2 clone 14 | 607.00 | 77.30 | 1519.00 | 94.50 | 16.80 | 1139.16 | 6.08 |
| pSSD-R2 clone 15 | 2604.00 | 80.80 | 2924.00 | 94.10 | 2.49 | 12.03 | 2.27 |
| pSSD-R2 clone 16 | 421.00 | 132.00 | 479.00 | 155.00 | 46.04 | 318.22 | 2.79 |
| pSSD-R2 clone 17 | 371.00 | 132.00 | 2751.00 | 86.80 | NA | NA | NA |
| pSSD-R2 clone 18 | 2503.00 | 81.40 | 3033.00 | 92.70 | 1.87 | 13.37 | 2.84 |
| pSSD-R2 clone 19 | 235.00 | 128.00 | 4100.00 | 84.00 | 42.93 | 1542.95 | 5.17 |
| pSSD-R2 clone 20 | 4804.00 | 94.50 | 4125.00 | 94.50 | 13.07 | 426.52 | 5.03 |
| pSSD-R2 clone 21 | 449.00 | 94.90 | 2802.00 | 88.60 | 115.10 | 1669.96 | 3.86 |
| pSSD-R2 clone 22 | 1950.00 | 95.90 | 4789.00 | 89.60 | 97.68 | 3199.54 | 5.03 |
| pSSD-R2 clone 23 | 692.00 | 70.40 | 3597.00 | 92.10 | 65.33 | 315.54 | 2.27 |
| pSSD-R2 clone 24 | 1835.00 | 109.00 | 3468.00 | 112.00 | 9.33 | 644.45 | 6.11 |
| pSSD-R2 clone 25 | 325.00 | 76.00 | 4051.00 | 119.00 | 138.74 | 5404.32 | 5.28 |
| pSSD-R2 clone 26 | 604.00 | 123.00 | 3184.00 | 91.70 | 0.62 | 2625.95 | 12.04 |
| pSSD-R2 clone 28 | 3005.00 | 94.20 | 3353.00 | 85.90 | 0.00 | 1.34 | NA |
| pSSD-R2 clone 29 | 3941.00 | 78.40 | 3965.00 | 93.60 | 0.00 | 20.06 | NA |
| pSSD-R2 clone 30 | 1442.00 | 95.40 | 3500.00 | 85.20 | 5.60 | 33.43 | 2.58 |
| pSSD-R2 clone 31 | 305.00 | 71.70 | 3686.00 | 92.70 | 11.20 | 175.15 | 3.97 |
| pSSD-R2 clone 32 | 1041.00 | 95.90 | 4292.00 | 96.30 | 0.00 | 1.34 | NA |
| pSSD-R2 clone 33 | 9025.00 | 82.00 | 3953.00 | 89.10 | 121.94 | 7039.52 | 5.85 |
| pSSD-R2 clone 34 | 2488.00 | 96.90 | 3313.00 | 94.70 | 74.04 | 1603.11 | 4.44 |
| pSSD-R2 clone 35 | 1391.00 | 148.00 | 2897.00 | 102.00 | NA | NA | NA |
| pSSD-R2 clone 36 | 2299.00 | 116.00 | 3005.00 | 103.00 | 44.80 | 2859.93 | 6.00 |
| pSSD-R2 clone 37 | 367.00 | 123.00 | 3663.00 | 96.00 | 57.86 | 1163.23 | 4.33 |
| pSSD-R2 clone 38 | 222.00 | 92.20 | 4214.00 | 86.90 | 54.75 | 119.00 | 1.12 |
| pSSD-R2 clone 39 | 429.00 | 71.00 | 3033.00 | 92.30 | 79.01 | 3549.84 | 5.49 |
| pSSD-R2 clone 40 | 551.00 | 105.00 | 3881.00 | 97.80 | 135.01 | 807.57 | 2.58 |
| pSSD-R2 clone 41 | 3697.00 | 111.00 | 4398.00 | 92.50 | 2.49 | 22.73 | 3.19 |
| pSSD-R2 clone 42 | 482.00 | 64.80 | 3042.00 | 97.30 | 54.13 | 1160.55 | 4.42 |
| pSSD-R2 clone 43 | 4506.00 | 92.90 | 2924.00 | 95.90 | 16.18 | 197.88 | 3.61 |
| pSSD-R2 clone 44 | 742.00 | 91.50 | 3204.00 | 91.80 | 149.94 | 3187.51 | 4.41 |
| pSSD-R2 clone 45 | 1575.00 | 98.50 | 3343.00 | 92.40 | 0.00 | 1.34 | NA |
| pSSD-R2 clone 46 | 2130.00 | 116.00 | 4452.00 | 92.60 | 1.24 | 215.26 | 7.43 |
| pSSD-R2 clone 47 | 5344.00 | 106.00 | 3364.00 | 113.00 | 0.62 | 74.87 | 6.91 |

**Supplementary Table 4. Randomly picked sorted clones display and fragment counts.** 47 random single colonies were picked after two round sorting (replicate 1) for single-clone flow cytometric assessment of display. Values represent median FLAG or median V5 signal of the non-negative population. rCV= robust coefficient of variation. The identity of the clones was determined by individual Sanger sequencing of the plasmid. The corresponding counts in unsorted and sorted samples as determined by sequencing are also shown. FPTM= fragments per ten million. Clone 17 and 35 were not detected in either unsorted or sorted sequenced samples.

| Hit # | RefSeq or Ensembl ID | Gene Symbol | Fragment length (AA) | Expected MW FLAG-frag-V5 (kDa) |
|---|---|---|---|---|
| 1 | NM_018403 | DCP1A | 154 | 19.55 |
| 2 | NM_004568, | SERPINB6 | 77 | 12.38 |
| 3 | NM_001348 | DAPK3 | 73 | 11.67 |
| 4 | NM_023007 | JMJD4 | 107 | 8.93 |
| 5 | NM_005977 | RNF6 | 58 | 9.36 |
| 6 | NM_002503 | NFKBIB | 111 | 15.48 |
| 7 | NM_006702 | PNPLA6 | 77 | 11.32 |
| 8 | NM_001569 | IRAK1 | 60 | 9.74 |
| 9 | NM_005587 | MEF2A | 58 | 9.25 |
| 10 | NM_014303 | PES1 | 45 | 8.36 |
| 11 | NM_003870 | IQGAP1 | 49 | 8.47 |
| 12 | NM_001625 | AK2 | 67 | 9.97 |
| 13 | NM_001746 | CANX | 47 | 8.71 |
| 14 | NM_015254 | ANKS1A | 43 | 7.69 |
| 15 | NM_001416 | EIF4A1 | 71 | 10.86 |
| 16 | ENST00000553917 | RALGAPA1 | 131 | 17.46 |
| 17 | NM_015509 | NECAP1 | 93 | 13.02 |
| 18 | NM_024923 | NUP120 | 69 | 10.55 |
| 19 | NM_001456 | FLNA | 61 | 9.05 |
| 20 | NM_022845 | CBFB | 118 | 16.84 |

**Supplementary Table 5. Identity and expected molecular weight of the random fragments picked from sorted cells after secretion screening.** Fragments were sequenced using the Sanger method, and mapped against human RefSeq transcripts for identification using BLASTn. Fragment length was calculated excluding the FLAG and V5 tags. Together, the N-terminal FLAG tag and C-terminal V5 add an additional 29 AA to the fragment (roughly 3 kDa). Note that fragment 16 maps to the C-terminus of the EIF4A1 ORF and the start of the annotated 3' UTR, and is in a different frame than the annotated protein.

| | | Enriched | Depleted |
|---|---|---|---|
| replicate 1 | # of fragments | 20,635 | 147,334 |
| | fraction of all (%) | 12.12 | 86.55 |
| replicate 2 | # of fragments | 21,480 | 145,806 |
| | fraction of all (%) | 12.62 | 85.65 |
| replicate 3 | # of fragments | 20,223 | 147,147 |
| | fraction of all (%) | 11.88 | 86.44 |
| **concordant** | # of fragments | 11,625 | 136,531 |
| | fraction of all (%) | **6.83** | **80.21** |

**Supplementary Table 6. *S. cerevisiae* enrichment and depleted fragment concordance between replicate screens.** From the 170 226 common in-frame fragments, roughly 20,000 fragments, or 12.21%, are detected as enriched after sorting in each replicate, and 86.22% as depleted. About 6.83%, or 11,625 fragments, are consistently enriched in all three replicates, and 80.21% consistently depleted. The remaining 12.98% of fragments are discordant between replicates, or below the set cut-offs for enrichment/depletion classification.

| sample | | # reads | fraction of reads (%) |
|---|---|---|---|
| total | | 725,021,482 | 100.00 |
| passing filter (PF) | | 674,935,720 | **93.09** |
| PF + barcode identified | unsorted 1 | 180,876,078 | 26.80 |
| | unsorted 2 | 170,163,740 | 25.21 |
| | unsorted 3 | 156,047,813 | 23.12 |
| | sorted 1 | 42,305,875 | 6.27 |
| | sorted 2 | 40,564,218 | 6.01 |
| | sorted 3 | 51,868,098 | 7.68 |
| | **total** | 641,825,822 | **95.09** |

| sample | fraction of reads (%) remaining after | |
|---|---|---|
| | Illumina sequence trimming | FLAG/V5 trimming |
| unsorted 1 | 99.46 | 96.22 |
| unsorted 2 | 99.49 | 97.09 |
| unsorted 3 | 99.46 | 95.75 |
| sorted 1 | 99.35 | 96.44 |
| sorted 2 | 99.35 | 96.83 |
| sorted 3 | 99.63 | 97.64 |
| **total** | **99.46** | **96.66** |

**Supplementary Table 7. *P. pastoris* secretability screening sequencing and trimming statistics.** Left: number of reads (twice the number of pairs) obtained after Illumina sequencing of 3 replicate samples. Data was summarized over four lanes. Of the roughly 675 million reads passing the Illumina quality filter (PF), about 95% were barcode identified. Right: read statistics after trimming. In a first round, all sequences matching Illumina flow cell or primer binding sequences were removed (on average less than 1% of sequences). Next, all FLAG+SfiI and V5+PacI sequences were trimmed off, and untrimmed or very short (< 20bp) reads were discarded, leaving roughly 97% of sequences for mapping.

| | | unsorted 1 | unsorted 2 | unsorted 3 | sorted 1 | sorted 2 | sorted 3 |
|---|---|---|---|---|---|---|---|
| | input total reads | 173,937,472 | 164,365,482 | 148,609,312 | 40,534,608 | 39,021,504 | 50,455,798 |
| **R 1** | # reads | 86,968,736 | 82,182,741 | 74,304,656 | 20,267,304 | 19,510,752 | 25,227,899 |
| | mapped (%) | 81.02 | 81.16 | 80.31 | 82.67 | 82.82 | 83.74 |
| | unambiguous (%) | 21.72 | 21.44 | 20.84 | 20.17 | 20.59 | 20.29 |
| | ambiguous (%) | 59.30 | 59.72 | 59.48 | 62.49 | 62.23 | 63.45 |
| **R 2** | # reads | 86,968,736 | 82,182,741 | 74,304,656 | 20,267,304 | 19,510,752 | 25,227,899 |
| | mapped (%) | 80.72 | 80.88 | 80.03 | 82.39 | 82.54 | 83.46 |
| | unambiguous (%) | 21.63 | 21.36 | 20.75 | 20.11 | 20.52 | 20.22 |
| | ambiguous (%) | 59.08 | 59.52 | 59.28 | 62.29 | 62.05 | 63.24 |
| **total** | mated pairs (%) | 77.92 | 78.12 | 77.26 | 79.88 | 79.89 | 80.93 |
| | bad pairs (%) | 0.96 | 0.94 | 0.99 | 0.85 | 0.95 | 0.93 |
| | properly paired (%) | 77.92 | 78.17 | 77.26 | 79.88 | 79.89 | 80.93 |
| | singletons (%) | 1.99 | 1.91 | 1.92 | 1.80 | 1.83 | 1.74 |

**Supplementary Table 8. *P. pastoris* secretability screening mapping statistics.** PF, barcode-identified and trimmed reads were mapped to the human hg38 transcriptome of known protein-coding genes using BBMap. The many ambiguous read mappings, owing to the multiple transcript isoforms per gene, were assigned to a single best mapped transcript for read counting. Downstream fragment counting was further performed using properly-paired reads only.

| sample | # proper pairs mapped | # fragments |
|---|---|---|
| unsorted 1 | 67,765,497 | 1,042,800 |
| unsorted 2 | 64,238,604 | 1,038,321 |
| unsorted 3 | 57,405,725 | 1,340,806 |
| sorted 1 | 16,189,765 | 394,356 |
| sorted 2 | 15,587,472 | 599,641 |
| sorted 3 | 20,416,465 | 860,829 |
| **total** | 241,603,528 | // |

| sample | Average coverage per covered base | Bases covered (%) | Combined bases covered (%) |
|---|---|---|---|
| unsorted 1 | 564.56 | 33.48 | 35.78 |
| sorted 1 | 195.64 | 22.24 | |
| unsorted 2 | 536.68 | 34.07 | 38.10 |
| sorted 2 | 146.19 | 29.84 | |
| unsorted 3 | 442.63 | 37.57 | 41.26 |
| sorted 3 | 168.36 | 34.20 | |

**Supplementary Table 9. *P. pastoris* secretability screening fragment counts and transcriptome coverage statistics.** Left: number of properly mapped read pairs and the corresponding number of different cDNA fragments obtained per sample. Right: to obtain an idea of the fraction of transcriptome bases covered in our library, coverage statistics were calculated for mappings against the hg38 transcriptome of canonical transcripts of known protein coding genes. On average, every base covered by at least one read is counted about 500 times in the unsorted samples, and +/- 150 times in sorted samples. About 30% of all transcriptome bases were covered with at least 3 reads in each sample, accounting for roughly 35-40% of the transcriptome per replicate (combined sorted and unsorted fragments). Note that these are estimates, as about 5% less reads can be mapped to the canonical transcriptome, and for fragments longer than 250 bp, the gap between the paired ends is not sequenced.

| | | Enriched | Depleted |
|---|---|---|---|
| **replicate 1** | # of fragments | 26,300 | 183,152 |
| | fraction of all (%) | 12.23 | 85.19 |
| **replicate 2** | # of fragments | 35,511 | 171,271 |
| | fraction of all (%) | 16.52 | 79.66 |
| **replicate 3** | # of fragments | 39,805 | 163,287 |
| | fraction of all (%) | 18.51 | 75.95 |
| **concordant** | # of fragments | 10,404 | 141,357 |
| | fraction of all (%) | **4.84** | **65.75** |

**Supplementary Table 10. *P. pastoris* enrichment and depleted fragment concordance between replicate screens.** From the 215 004 common in-frame fragments, between 12.2% and 18.5% are detected as enriched after sorting in each replicate, and between 76% and 85.2% as depleted. About 4.84%, or 10 404 fragments, are consistently enriched in all three replicates, and 65.75% consistently depleted. The remaining 29.41% of fragments are discordant between replicates, or below the set cut-offs for enrichment/depletion classification.

| | Human | | Baker's yeast | |
|---|---|---|---|---|
| | **Disordered** | **Not disordered** | **Disordered** | **Not disordered** |
| **Secretory** | 831 | 7412 | 22 | 317 |
| **Whole proteome** | 3674 | 16699 | 924 | 5797 |

**Supplementary Table 11. Disordered proteins in human and yeast secretome and proteome according to fractional disorder content.** Secretory proteins here are defined as being annotated with a signal peptide in Uniprot. The number of disordered proteins is calculated as the number of proteins with a disorder content >35% as calculated with RAPID. In human, the fraction of disordered secretory proteins is significantly lower than across the whole proteome (10.1% vs 18.0%, one-sided Fisher exact test, $p<2.2*10^{-16}$). A similar trend is observed in baker's yeast (6.5% vs 13.8%, one-sided Fisher exact test, $p=2.46*10^{-5}$).

| | Human | | Baker's yeast | |
|---|---|---|---|---|
| | **Disordered** | **Not disordered** | **Disordered** | **Not disordered** |
| **Secretory** | 2384 | 5859 | 91 | 248 |
| **Whole proteome** | 11390 | 8983 | 2899 | 3822 |

**Supplementary Table 12. Disordered proteins in human and yeast secretome and proteome according to number of disordered amino acids.** Secretory proteins here are defined as being annotated with a signal peptide in Uniprot. The number of disordered proteins here is calculated as the number of proteins with more than 50 disordered amino acids calculated with RAPID. In human, the fraction of disordered secretory proteins is significantly lower than across the whole proteome (28.9% vs 55.9%, one-sided Fisher exact test, $p<2.2*10^{-16}$). A similar trend is observed in baker's yeast (26.8% vs 43.1%, one-sided Fisher exact test, $p=9.44*10^{-10}$).

| Architecture | *S. cerevisiae* | | | | | *P. pastoris* | | | | |
|---|---|---|---|---|---|---|---|---|---|---|
| | # Enr. | # Depl. | Fraction Enr. | Fraction Depl. | Fold change | # Enr. | # Depl. | Fraction Enr. | Fraction Depl. | Fold Change |
| 3 Solenoid | 1 | 2 | 0.001 | 0.000 | 3.902 | 3 | 8 | 0.002 | 0.001 | 2.943 |
| 5-Stranded Propeller | 3 | 9 | 0.004 | 0.001 | 2.601 | 1 | 12 | 0.001 | 0.001 | 0.654 |
| 6 Propellor | 3 | 12 | 0.004 | 0.002 | 1.951 | 3 | 48 | 0.002 | 0.004 | 0.490 |
| Distorted Sandwich | 6 | 27 | 0.007 | 0.004 | 1.734 | 15 | 52 | 0.011 | 0.005 | 2.264 |
| Beta Complex | 7 | 32 | 0.009 | 0.005 | 1.707 | 9 | 41 | 0.007 | 0.004 | 1.723 |
| Sandwich | 63 | 353 | 0.077 | 0.055 | 1.393 | 137 | 801 | 0.100 | 0.075 | 1.342 |
| Ribbon | 11 | 65 | 0.013 | 0.010 | 1.321 | 42 | 181 | 0.031 | 0.017 | 1.821 |
| Up-down Bundle | 49 | 295 | 0.060 | 0.046 | 1.296 | 101 | 644 | 0.074 | 0.060 | 1.231 |
| 3-Layer(aba) Sandwich | 210 | 1458 | 0.255 | 0.227 | 1.124 | 241 | 2236 | 0.177 | 0.209 | 0.846 |
| 7 Propellor | 21 | 147 | 0.026 | 0.023 | 1.115 | 27 | 215 | 0.020 | 0.020 | 0.986 |
| Roll (Alpha-Beta) | 27 | 198 | 0.033 | 0.031 | 1.064 | 108 | 762 | 0.079 | 0.071 | 1.112 |
| Alpha-Beta Complex | 44 | 331 | 0.053 | 0.052 | 1.037 | 71 | 557 | 0.052 | 0.052 | 1.000 |
| Roll (mainly Beta) | 37 | 280 | 0.045 | 0.044 | 1.031 | 108 | 762 | 0.079 | 0.071 | 1.112 |
| Irregular | 3 | 23 | 0.004 | 0.004 | 1.018 | 32 | 65 | 0.023 | 0.006 | 3.863 |
| Beta Barrel | 33 | 259 | 0.040 | 0.040 | 0.994 | 63 | 405 | 0.046 | 0.038 | 1.221 |
| 2-Layer Sandwich | 146 | 1180 | 0.177 | 0.184 | 0.965 | 236 | 1818 | 0.173 | 0.170 | 1.019 |
| Alpha-Beta Barrel | 24 | 206 | 0.029 | 0.032 | 0.909 | 28 | 232 | 0.021 | 0.022 | 0.947 |
| Single Sheet | 6 | 56 | 0.007 | 0.009 | 0.836 | 19 | 107 | 0.014 | 0.010 | 1.394 |
| Alpha/alpha barrel | 4 | 38 | 0.005 | 0.006 | 0.821 | 2 | 23 | 0.001 | 0.002 | 0.682 |
| 3-Layer(bba) Sandwich | 6 | 57 | 0.007 | 0.009 | 0.821 | 3 | 82 | 0.002 | 0.008 | 0.287 |
| Alpha-Beta Horseshoe | 2 | 20 | 0.002 | 0.003 | 0.780 | 3 | 59 | 0.002 | 0.006 | 0.399 |
| Orthogonal Bundle | 83 | 861 | 0.101 | 0.134 | 0.752 | 151 | 1567 | 0.111 | 0.146 | 0.756 |
| Trefoil | 4 | 44 | 0.005 | 0.007 | 0.709 | 8 | 74 | 0.006 | 0.007 | 0.848 |
| Alpha Horseshoe | 10 | 333 | 0.012 | 0.052 | 0.234 | 35 | 530 | 0.026 | 0.049 | 0.518 |
| Box | 0 | 0 | 0 | 0 | 0 | 2 | 7 | 0.001 | 0.001 | 2.242 |

**Supplementary Table 13. CATH domain architectures of representative enriched and depleted fragments mapping to PDB structures in *S. cerevisiae* and *P. pastoris*.** While most architectures are roughly equally represented in both enriched and depleted fragments, a few architectures are found over- or underrepresented in secretable (enriched) fragments. Of those that were sampled sufficiently (detected at least in 0.5% of fragments in enriched or depleted sets), the Alpha Horseshoe architecture is substantially underrepresented in secretable fragments, while Beta Complex and Distorted Sandwiches are overrepresented in both yeasts. Fold change = fraction enriched / fraction depleted. # = number of occurrences of a particular architecture in representative enriched (Enr.) or depleted (Depl.) fragments mapping to one or multiple PDB structures. Fraction Enr. or fraction Depl. = # Enr. or Depl. / total number of all architecture occurrences in Enr. or Depl.

| Dataset | # of positives | # of negatives | total |
|---|---|---|---|
| *S. cerevisiae* | 6,194 | 68,062 | 74,256 |
| *P. pastoris* | 8,010 | 110,001 | 118,011 |

**Supplementary Table 14**. **Final dataset composition for (deep) machine learning training**. For both the *S. cerevisiae* and *P. pastoris* datasets, all sequences shorter than 50 amino acids were removed.

| Method | *S. cerevisiae* | *P. pastoris* |
|---|---|---|
| Polarity | 0.751 | 0.756 |
| Hydrophobicity | 0.764 | 0.746 |
| Average area buried | 0.767 | 0.741 |
| Buried residues | 0.760 | 0.766 |
| Bulkiness | 0.760 | 0.751 |
| Molar refraction | 0.764 | 0.765 |
| Recognition factors | 0.760 | 0.742 |
| Molecular weight | 0.772 | 0.756 |
| Transmembrane tendency | 0.776 | 0.757 |
| HPLC | 0.769 | 0.761 |
| Ensemble | 0.781 | 0.772 |

**Supplementary Table 15. Classification results using gradient boosted trees**. The AUROC over a ten-fold cross-validation experiment is listed for the individual classifiers, as well as the ensemble.

| Layer | Details |
|---|---|
| convolutional layer (+ ReLU) | 100 filters of size 3 |
| dropout layer | p = 0.2 |
| max pooling layer | pool size 4 |
| convolutional layer (+ ReLU) | 100 filters of size 7 |
| dropout layer | p = 0.2 |
| max pooling layer | pool size 4 |
| convolutional layer (+ ReLU) | 100 filters of size 7 |
| dropout layer | p = 0.2 |
| variable to fixed length transformation method (if applicable) | |
| fully-connected layer | 32 neurons |
| output layer (sigmoid) | 1 neurons |

**Supplementary Table 16. Hyperparameters for the final deep learning architecture**. Hyperparameters were determined by means of a grid search.

| Method | # of parameters | *S. cerevisiae* | *P. pastoris* |
|---|---|---|---|
| (baseline) zero padding only | 185K | 0.777 (0.004) | 0.767 (0.001) |
| global max pooling | 150K | 0.779 (0.002) | 0.768 (0.001) |
| *K*-max pooling | 162K | 0.778 (0.002) | 0.768 (0.001) |
| bidirectional GRU | 330K | 0.777 (0.003) | 0.768 (0.001) |

**Supplementary Table 17. Classification results using convolutional neural networks**. The average AUROC over ten ten-fold cross-validation experiments is listed, along with the standard deviation.
